# f-BGM enables fungi-specific genome mining in high accuracy and interpretability

**DOI:** 10.1101/2025.07.08.662937

**Authors:** Yiran Zhou, Dong Liu, Zequan Huang, Xingsi Xie, Wenhan Lin, Liangren Zhang, Aili Fan, Zhenming Liu

**Author notes:** Corresponding authors: Zhenming Liu,; Aili Fan. These authors contributed equally to this study (Yiran Zhou, Dong Liu).

## Abstract

Emerging artificial intelligence (AI)-based genome mining methods have revolutionized the paradigm of bacterial secondary metabolite (SM) discovery. Fortunately, recent data accumulation of fungal biosynthetic gene clusters (BGC) fairly offers opportunities for systematic development and evaluation of fungi-specific pipelines. In this work, we proposed a deep learning framework termed as f-BGM specifically for fungal genome mining. By designing a novel self-attention-based architecture to augment inter-domain associations in local genomic contexts, f-BGM exhibits superior performance over existing AI-based methods in both in-distribution and out-of-distribution benchmark tests for BGC detection. Further analyses demonstrate that f-BGM is of decent interpretability on deciphering single-domain and -protein importance, as well as inter-domain partnership. By establishing additional binary classification models, f-BGM also achieves high-quality identification of core enzymes within given BGCs. Finally, case studies of f-BGM-driven genome mining in marine fungi uncovers biosynthetic potential underestimated by the rule-based method antiSMASH, as supported by experimental and computational validation.

## Introduction

Secondary metabolites (SM) serve as a precious resource for drug discovery, of which the fungal SMs constitute an essential part. Statistically, approximately 25% of the approved natural drugs are fungi-sourced [1], such as penicillin [2], ciclosporin [3], pravastatin [4] and so on [5, 6], underlining their importance.

In microbes, the biosynthesis of SMs is usually mediated by genomically co-localized genes encoding necessary enzymes for the assembly and tailoring of metabolic scaffolds, also termed as biosynthetic gene clusters (BGCs) [7]. Rapid development of biotechnologies and computer science provides new insights for genome-driven SM discovery [8]. For example, by deciphering gene clusters with biosynthetic potential using *in silico* genome mining methods, combined with heterologous expression of these clusters in genetically tractable host, corresponding SMs can be isolated and elucidated from the post-fermented extracts of engineered hosts [9]. Since this strategy only relies on genomic data as source material, it can serve as a universal solution for discovering SMs and establishing BGC-SM linkages, circumventing crucial limitations of conventional native organism-required paradigm trapped in the following situations: (1) The native organisms are inaccessible; (2) In standard laboratory environment the SMs are low-expressed in native organisms [9]. Therefore, as its first step, how to achieve high-quality genome mining is obviously a fundamental problem.

Existing methods for *de novo* genome mining can be roughly classified into two categories: (1) Rule-based methods. One representative is the well-known antiSMASH [10], where the protein domains encoded by the input genomes are first identified and then a series of predefined rules, such as hit scores of specific domains, co-occurrent relationship of specific domain-domain pairs and so on, are used for BGC detection. Such methods are effective for discovering typical BGCs. However, the prior knowledge-dependent manner limits their generalization capability on novel BGC types; (2) Artificial intelligence (AI)-based methods. This category of methods aims to acquire biosynthetic signatures via automatic model training rather than relying on rigid handcrafted rules. The robust adaptivity has recruited growing scientific interests in recent years [11–13].

However, existing AI-based methods are largely developed for bacteria but rarely for fungi [11–13]. One contributing factor to the predicament is the data scarcity of well-annotated fungal BGCs. In the previously largest-scale BGC database, MIBiG (version 3.1) [7], only 440 BGC entries are curated, which is insufficient for systematic algorithm design and evaluation. Fortunately, the limitation has been properly mitigated with the recent release of FunBGCs, a fungi-specific BGC database [14]. First, the FunBGCs database achieves a significant expansion in data volume (768 entries). More importantly, the dataset underwent manual correction on BGC borders and membership, further ensuring the data quality.

From a methodological perspective, current methods always employ sequence-based algorithmic frameworks such as hidden Markov model (HMM), bidirectional long-short term memory (Bi-LSTM) and conditional random field (CRF) to model genomic contexts, combined with feature engineering strategies using evolutionarily conserved protein domains [11–13]. However, the straightforward design is less powerful to capture inter-domain and inter-gene cooperative relationship within BGCs, which is important for characterizing biosynthetic behaviors and improving model interpretability. Self-attention mechanism is a deep learning architecture originally designed for natural language processing problems [15]. However, in recent years, its applications have been extended to biomedical fields such as prediction of macromolecular folding conformations [16, 17], drug-target interactions [18], conditional drug design [19] and so on [20, 21] due to its context-aware modeling manner by explicit attention weights measuring inter-unit associations.

In the current work, by leveraging a novel self-attention-based module explicitly modeling inter-domain associations in local genomic contexts, as well as an advanced protein representation method [22], we proposed a deep learning algorithm specifically for the fungal genome mining research termed as Fungal Biosynthetic Genome Miner (f-BGM). The following content will first introduce the overall architecture of f-BGM and its benchmarking performance against existing algorithms in BGC detection. Then the results of ablation experiments and interpretability analyses will be presented to understand the key components and decision mechanisms of f-BGM. After further validating its performance in core enzyme identification, the complete f-BGM pipeline will be demonstrated through practical genome mining research in marine fungi, regarding the rule-based antiSMASH as control.

## Results

### Overview of f-BGM model

The f-BGM model aims to detect BGCs from input genomic contexts at first, then identify core enzymes encoded by the predicted BGCs (Fig. 1). For the BGC detection purpose, three sequential neural network modules were designed (Fig. 1a): (1) Self-attention-based module for short-range information interaction among consecutive ORFs (SRM); (2) LSTM-based module for long-range message passing across whole genomic contexts (LRM); (3) Output module to generate ORF-level probabilistic scores of BGC membership.

**Figure 1.**
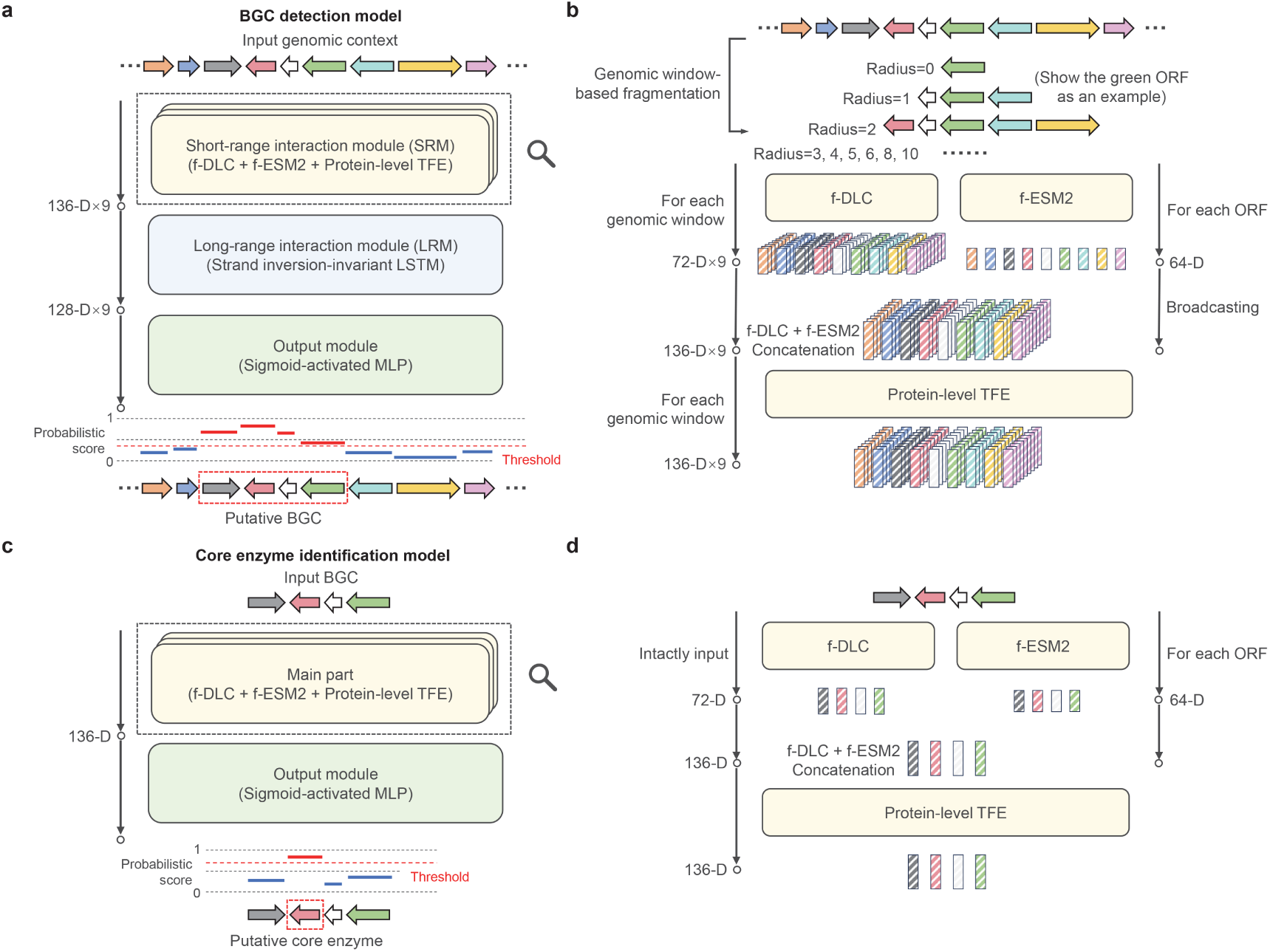
Overview of f-BGM model. **a** The BGC detection model receives genomic contexts as input and outputs ORF-level probabilistic scores representing BGC membership. The entire process consists of three sequential steps: (1) self-attention-based short-range information interaction (by SRM), (2) long-short-term memory (LSTM)-based long-range information passing (by LRM) and (3) probabilistic score generation (by OM). **b** Detailed illustration of the SRM. SRM integrates two pretrained models respectively capturing (1) inter-domain locally co-occurrent relationship in fungal genome (f-DLC) and (2) amino acid patterns within fungal protein sequences (f-ESM2). The input genomic contexts are first fragmented into multiple sizes of genomic windows and fed into f-DLC to enable domain-level information interaction and generate window-wise ORF-level embeddings, meanwhile the ORF-encoded protein sequences are passed into f-ESM2. The f-DLC-output embeddings and broadcasted f-ESM2-output embeddings are further concatenated and fed into an additional transformer encoder for protein-level information interaction (protein-level TFE). **c** The core enzyme identification model receives BGCs as input and outputs ORF-level probabilistic scores representing the presence or absence of target core enzymes. **d** The main part of core enzyme identification model mimics the SRM architecture in (**b)** but without genomic window-based fragmentation on the input BGCs due to their determined borders.

SRM is the core component of the BGC detection model (Fig. 1b), where two pretrained models respectively acquiring the knowledge of (1) inter-Domain Locally Co-occurrent relationship in Fungal genome (f-DLC) and (2) amino acid patterns within fungal protein sequences (f-ESM2), are employed. f-DLC was innovatively proposed by this work, which adopts a specially designed multi-layer self-attention architecture to model the intricate domain-to-domain relationship within single protein, domain-to-domain relationship within multiple proteins and domain-to-protein relationship (See ‘Methods and materials’; Supplemental Fig. S1-S2). Given limited number of consecutive ORFs (≤26) as input, f-DLC can generate 72-dimensional (72-D) ORF-level embeddings encoding the inter-domain associations; f-ESM2 is a derivate of the well-known protein language model ESM2 (150M-parameter version) [22] with the modifications of (1) output dimensionality reduction and (2) fungi-specific fine-tuning (See ‘Methods and materials’). For input protein sequences, f-ESM2 can generate 64-D representation embeddings. The complete pipeline of SRM is as follows: (1) For the given genomic context, construct a series of genomic windows by binning each ORF with flanking ORFs. Here the windows are set in nine different sizes with the radiuses of 0, 1, 2, 3, 4, 5, 6, 8 and 10; (2) Feed the genomic windows into f-DLC, thereby nine 72-D embeddings reflecting inter-domain interactivity in multi-scale neighborhoods will be generated for each ORF; (3) Input the ORF-encoded protein sequences into f-ESM2 to generate 64-D representation embeddings; (4) Broadcast the f-ESM2-output embeddings and perform f-DLC + f-ESM2 embedding concatenation for each genomic window (136-D); (5) Feed the fusion embeddings under each window size into an additional transformer encoder (TFE) to enable protein-level information interaction, the resulting embeddings (136-D) will be passed to downstream LRM.

LRM assigns independent LSTM neural networks to process the SRM-output embedding sequences under different genomic window sizes. Here the LSTM blocks are configured with strand direction-independent adaptations. That is to say, the predictive consistency can be maintained regardless of the directions of ORF arrangements (See ‘Methods and materials’). The output embeddings of individual LSTM blocks are 128-D.

Finally, the LRM-output embeddings are concatenated across genomic windows (128-D×9) and projected into probabilistic scores by a sigmoid-gated multi-layer perceptron (MLP).

Due to the data scarcity of genomic contigs with clearly labeled BGC membership, the BGC detection model was actually trained on synthetic genomic contexts generated by random concatenation between positive gene cluster samples (i.e., the BGCs) and constructed negative gene cluster samples, as previously described (See ‘Methods and materials’) [11–13].

The FunBGCs dataset additionally provides core enzyme annotations such as non-reducing polyketide synthase (NR-PKS), non-ribosomal peptide synthetase (NRPS), canonical class I terpene cyclase (TC-Class1) and so on that directly reveal BGCs’ biosynthetic behaviors. Based on this information, we summarized seven major families of core enzymes existing in ≥30 BGC samples, including PKS, NRPS, TC, PT, PKS-NRPS, chimeric TS and PPPS (Supplemental Table S1). Correspondingly, seven binary classification models were separately trained to identify respective core enzyme family within given BGCs (Fig. 1c). The main part of the core enzyme identification model adopts an SRM-similar self-attention-based design, where f-DLC, f-ESM2 and a protein-level TFE are integrated. However, it receives intact BGC samples as input without genomic window-based fragmentation due to their determined borders (Fig. 1d). The resulting ORF-level embeddings (136-D) are finally projected into probabilistic scores representing the presence or absence of target core enzymes (Fig. 1c).

### f-BGM outperforms existing algorithms in fungal BGC detection

Four existing algorithms including ClusterFinder [11], DeepBGC [12], GECCO [13] and TOUCAN [23] were reimplemented as baselines for the fungal BGC detection problem (See ‘Methods and materials’). The performance of f-BGM and baselines was benchmarked through two tasks: (1) Real BGC detection, where the algorithms were evaluated on real genomic contigs with annotated BGCs (Supplemental Table S2-S4); (2) Simulated genomic context mining, where synthetic genomic contexts were used for algorithm evaluation (See ‘Methods and materials’). In addition, three dataset partitioning schemes including one in-distribution and two out-of-distribution schemes were introduced to define training/validation/test sets. Under the in-distribution scheme, the training/validation/test sets were generated by completely random partitioning on the FunBGCs dataset at a 4:1:1 ratio, ensuring their distributional homogeneity. This configuration covers most real-world application scenarios. In contrast, the out-of-distribution schemes aim to engineer distributional shifts between the datasets used for model development (i.e., the training/validation sets) and evaluation (i.e., the test set), which place more emphasis on algorithms’ generalization capability. The introduced out-of-distribution schemes include: (1) Cross-SM-class scheme, where the FunBGCs dataset was partitioned into six subsets (Cluster 1-6) based on the chemical diversity of BGCs’ representative SM products (See ‘Methods and materials’; Supplemental Fig. S3). Under the scheme, algorithms’ performance was evaluated in a rotational cross-validation manner supplemented by balanced subsampling to systematically interrogate algorithms’ capability in identifying uncharacterized biosynthetic novelty (See ‘Methods and materials’); (2) Cross-dataset scheme, where the BGCs from different data sources, FunBGCs and MIBiG, were allocated to the training/validation sets and the test set, respectively. The training and validation sets were randomly assigned at a 4:1 ratio. Under both in-distribution and out-of-distribution schemes, the constructed negative gene cluster samples were co-grouped with their associated query BGCs. Algorithms’ performance metrics were calculated at ORF level and averaged over 16 rounds of *de novo* experiments to mitigate influences of random factors such as training/validation/test dataset partitioning.

For the real BGC detection task, it should be noted that the ORFs located outside the known BGCs cannot be arbitrarily regarded as negatives because of their potential involvement in uncharacterized biosynthetic regions. The uncertainty in ground truth means conventional evaluation methods such as receiver operating characteristic curve (ROC curve) and precision-recall curve (P-R curve) are inapplicable here. To address this issue, we proposed the recall of known positives at specified top ratio as a robust performance metric (See ‘Methods and materials’). Considering in most application scenarios only the top highly scored ORFs are preferred for further investigation, here we focused on three standard top ratios including 1%, 5% and 10% for recall calculation (recall_TR=1%_, recall_TR=5%_ and recall_TR=10%_), among which recall_TR=5%_ was prioritized as the principal metric due to the moderate background size, while recall_TR=1%_ and recall_TR=10%_ served as secondary metrics. Moreover, given the ORF number across genomic contigs is extremely imbalanced (Q1=136, median=612, Q3=1,286) (Supplemental Table S4), the ORFs included for performance evaluation per contig were constrained by a specified number to avoid biases to the overlength contigs. To be more specific, for the contigs whose ORF number exceeding the threshold, the excessive ORFs distant from known BGCs were excluded for metric calculation. Here a series of ORF number thresholds including 64, 128, 256, 512 and 1,024 were enumerated to cover more situations. The results show that f-BGM outperforms the baseline algorithms in general regardless of dataset partitioning schemes (Fig. 2a; Supplemental Fig. S4a-b). Especially, when the ORF number threshold equals moderate 256, the comparison between f-BGM and the best-performing baselines (always GECCO) demonstrates recall_TR=5%_ improvements of 0.035 (0.748±0.041 vs 0.713±0.027), 0.032 (0.694±0.009 vs 0.662±0.008) and 0.038 (0.692±0.009 vs 0.654±0.014) in-distribution, cross-SM-class and cross-dataset benchmark tests, respectively (Fig. 2a). However, in occasional cases the superiority might be fluctuant. For example, in the cross-dataset benchmark test with the ORF number threshold of 512, f-BGM slightly underperforms DeepBGC in terms of recall_TR=10%_ metric (0.930±0.011 vs 0.933±0.008) (Supplemental Fig. S4b).

**Figure 2.**
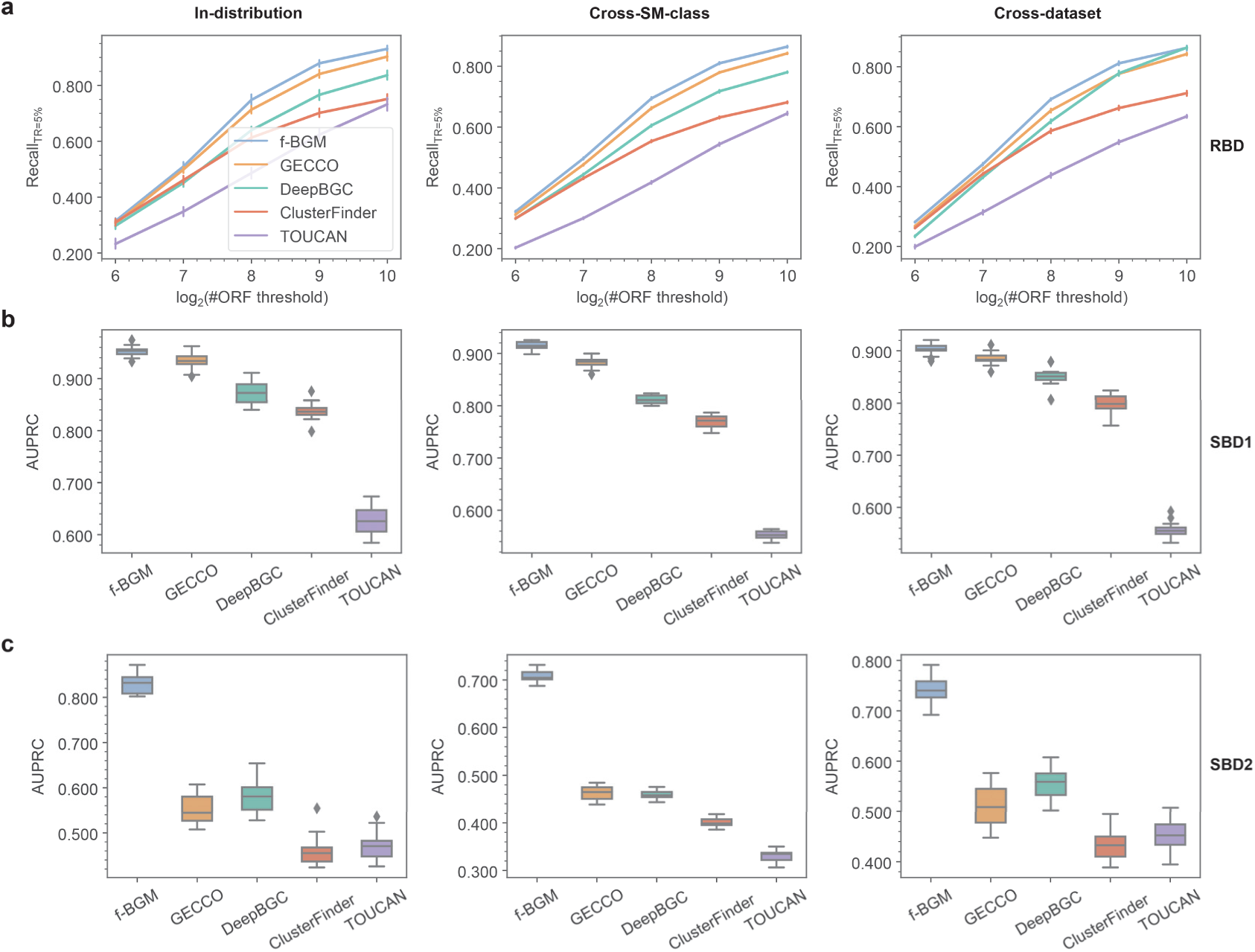
Performance of f-BGM in BGC detection benchmark tests. **a** Performance comparison among f-BGM and four baseline algorithms (ClusterFinder, DeepBGC, GECCO and TOUCAN) with regard to Recall_TR=5%_ in real BGC detection tasks under in-distribution, cross-SM-class and cross-dataset schemes. To avoid biases to overlength contigs, the ORFs included for analyses per contig are limited by a specified maximal number by removing excessive ORFs distant from the known positives. Here the ORF number thresholds of 64, 128, 256, 512 and 1024 are enumerated to cover more situations (x-axis). **b, c** Performance comparison among f-BGM and four baseline algorithms with regard to AUPRC in (**b)** standard simulated BGC detection and (**c)** challenging simulated BGC detection tasks under in-distribution, cross-SM-class and cross-dataset schemes. RBD: real BGC detection; SBD1: standard simulated BGC detection; SBD2: challenging simulated BGC detection.

For the simulated BGC detection task, the randomness introduced by the construction process of synthetic genomic contexts results in a broader diversity of gene arrangements, thus properly addressing current data scarcity of BGC-annotated real genomic contigs. In addition, the clearly labeled negative ORFs enable standard performance metrics such as areas under ROC curve (AUROC) and P-R curve (AUPRC) and F1-score. Here we selected AUPRC as the principal metric due to its sensitivity to false positives in class-imbalanced scenarios and comprehensiveness for performance evaluation, whereas AUROC and F1-score were secondarily referred. As expected, the results still confirm the superiority of f-BGM (Fig. 2b; Supplemental Fig. S4c-d). Specifically, f-BGM respectively achieves minimum (1) AUPRC improvements of 0.019 (0.953±0.010 vs 0.934±0.015), 0.033 (0.915±0.008 vs 0.882±0.010) and 0.018 (0.903±0.011 vs 0.885±0.012), (2) AUROC improvements of 0.002 (0.994±0.004 vs 0.992±0.003), 0.007 (0.988±0.002 vs 0.981±0.002) and 0.002 (0.983±0.003 vs 0.981±0.002) and (3) F1-score improvements of 0.022 (0.897±0.012 vs 0.875±0.015), 0.032 (0.861±0.009 vs 0.829±0.012) and 0.012 (0.845±0.012 vs 0.833±0.010) over baselines under individual dataset partitioning schemes. However, we noted the performance gains are less significant (AUPRC improvements <0.050). One explanation is that the negative gene cluster samples incorporated in the synthetic genomic contexts were constrained by a maximal BGC similarity of 0.5 (Type-1 negative samples, see ‘Methods and materials’), so that the positive-vs-negative discrimination might be of low difficulty and insufficient to reflect methodological differences among the algorithms. Therefore, we additionally orchestrated a challenging version of simulated BGC detection task by introducing another BGC-similar type of negative samples (Type-2 negative samples, see ‘Methods and materials’), as previous research suggested [12]. All the algorithms underwent from-scratch training for a new round of benchmark tests (Fig. 2c; Supplemental Fig. S4e-f). On the one hand, the involvement of the type-2 negative samples degrades all algorithms’ performance as expected. But on the other hand, f-BGM could maximally resist the negative influences. As a result, f-BGM significantly outperforms the baselines, where (1) AUPRC gaps of at least 0.247 (0.829±0.020 vs 0.582±0.039), 0.246 (0.708±0.013 vs 0.462±0.015) and 0.185 (0.741±0.028 vs 0.556±0.028), (2) AUROC gaps of at least 0.034 (0.981±0.003 vs 0.947±0.006), 0.040 (0.961±0.002 vs 0.921±0.004) and 0.016 (0.944±0.005 vs 0.928±0.005) and (3) F1-score gaps of at least 0.177 (0.763±0.017 vs 0.586±0.028), 0.165 (0.677±0.009 vs 0.512±0.011) and 0.132 (0.689±0.026 vs 0.557±0.017) were respectively observed under individual dataset partitioning schemes. Given the genes with some extent of BGC homogeneity contribute a large proportion of false positives in practical genome mining research, the results confirm f-BGM’s efficacy to identify and exclude such challenging negative gene clusters over the baselines. But unfortunately, the retrained f-BGM models fail to perform better in real BGC detection (Supplemental Fig. S5), suggesting the artificial construction pipeline of type-2 negative samples might induce shifts on real-world data distribution. Therefore, the downstream research still applied the previous model version. With expansion of open-access data repositories, the practical values of f-BGM would be further enhanced in future by training on real genomic contigs with complete ground truth instead of synthetic genomic contexts.

Overall, the benchmark tests jointly demonstrate the promising accuracy and generalization capability of f-BGM in fungal BGC detection.

### Ablation experiments reveal major contribution of f-DLC to f-BGM’s BGC detection performance

Given the decent performance of f-BGM in BGC detection, we further carried out ablation experiments to better understand the contribution of its individual components. Specifically, four f-BGM variants in absence of (1) f-DLC, (2) f-ESM2, (3) f-DLC pretraining and (4) LRM were trained and compared with the full model through in-distribution BGC detection tasks (Supplemental Fig. S6). First, all four variants underperform the full model in different degree, indicating each component’s contribution to the overall performance. To be more specific, for the real BGC detection task and the standard simulated BGC detection task with only type-1 negative samples, the variant without f-DLC perform worst with regard to respective primary evaluation metrics (i.e., recall_TR=5%_ and AUPRC), in contrast to slight performance reduction of the other three variants (Supplemental Fig. S6c/f). And for the challenging simulated BGC detection task involving type-2 negative samples, the performance of the variant without f-DLC pretraining also drastically decrease (Supplemental Fig. S6i). The results jointly demonstrate the pivotal role of f-DLC in f-BGM, and proper initialization of f-DLC parameters by pretraining is crucial for excluding BGC-similar negatives.

Considering that most existing algorithms only use domain features as input, we further focused on the f-BGM variant without f-ESM2, which also relies on Pfam domain information only. Despite the absence of protein sequence-based information presented by f-ESM2, f-BGM still outperforms the baselines (Supplemental Fig. S7), highlighting its state-of-the-art (SOTA) capability in deciphering latent inter-domain patterns within BGCs.

### f-BGM captures inter-domain and inter-gene associations of biological significance

In the SRM of f-BGM BGC detection model, the self-attention mechanism deployed in f-DLC (including single protein domain-level TFE and multi-protein domain-level TFE) and the protein-level TFE explicitly models inter-domain and inter-protein associations via softmax-defined attention weights, respectively. Therefore, to understand the acquired knowledge of f-BGM, we conducted a series of analyses to uncover the patterns underlying the attention weights, where the models built for the in-distribution prediction task were used. The attention weights were extracted by reinputting intact gene cluster samples of the test set into SRM without genomic window-based fragmentation, and then underwent robust z-score normalization to ensure cross-sample comparability (See ‘Methods and materials’). The self-loop attention connections were removed to avoid confounding factors.

First, assuming the BGCs contain more closely inner-associated submodules than the negative gene cluster samples, the top 1%, 5% and 10% of attention weights of individual TFEs within each sample were averaged and compared. Note that here the analytic targets towards single protein domain-level TFE are individual ORFs rather than whole gene cluster samples. As expected, the positive-derived top-ranked weights are significantly higher than those from negatives at both domain and protein levels (Fig. 3a), suggesting the behavior differences of f-BGM in positive-vs-negative discrimination.

**Figure 3.**
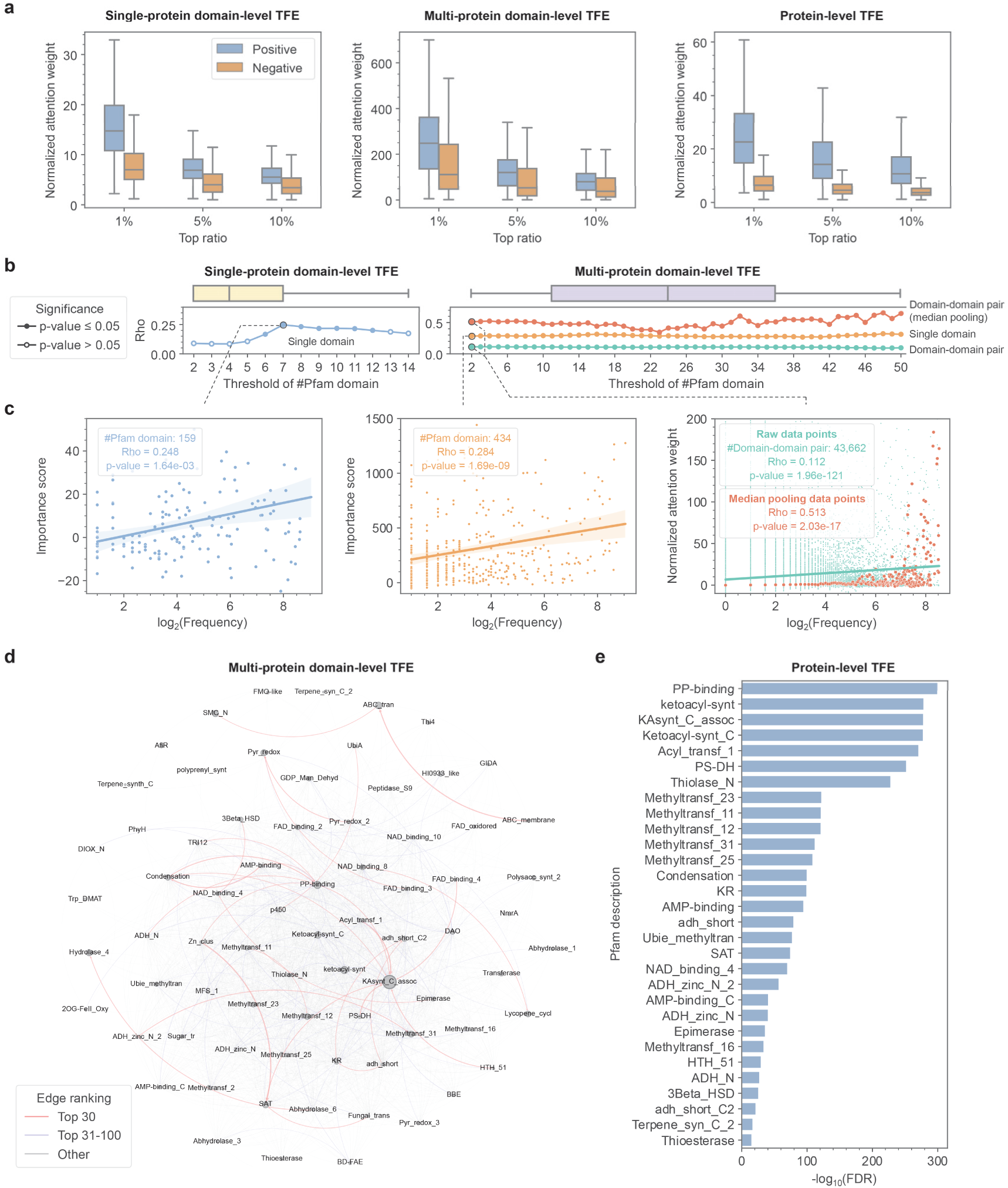
Interpretability analysis of f-BGM in BGC detection. **a** TFE-wise positive-vs-negative comparison among gene cluster samples with regard to inner attention weights at top ratios of 1%, 5% and 10%. **b** Spearman correlation analyses for single domains and domain-domain pairs, where the correlation coefficients rho (y-axis) between BGC-containing frequencies and f-BGM attention weights/attention weight-based importance scores are depicted with regard to minimal Pfam domain number required for the analytic objects (x-axis). The statistical significance of each data point is shown by solid/hollow style. The horizonal box plots show domain number distributions of single proteins and BGCs, respectively. **c** Examples of the positively correlative relationship shown in (**b)** at specific domain number threshold (7 for single-protein domain-level TFE and 2 for multi-protein domain-level TFE). **d** Attention weight network organized from 2,048 highest-frequency domain-domain pairs (among 74 domains) in BGCs. The node sizes and edge widths reflect single-domain importance and inter-domain attention weights, respectively. The edges are also color-coded by their attention weight ranking. **e** Fisher’s exact tests reveal top 30 Pfam domains enriched in the most important proteins of BGCs defined by f-BGM protein-level attention weights.

Further, we investigated whether the attention weights within BGCs are biased to specific subsets of domains or proteins. First at domain level, the weights received by each domain underwent subsequential cross-ORF and cross-BGC averaging to measure their overall importance under the f-BGM definition. Motivated by the hypothesis that the presence frequency of one domain in BGCs is indicative of its biosynthetic relevance, we conducted Spearman correlation analyses between the attention weight-based importance scores and the BGC-containing frequencies. The correlation coefficients rho and statistical significance information were depicted with regard to thresholds of minimal Pfam domain number required for the analytic targets (Fig. 3b-c). For the analysis towards single-protein TFE, positively correlative relationship was observed and it is especially significant (p-value≤0.05) when focusing on the top 25% of ORFs containing ≥7 domains (rho=0.248). From the multi-protein perspective, the positive correlation is robust regardless of the domain number thresholds (median rho=0.286), preliminarily demonstrating the interpretability of f-BGM. Similar analysis was also performed on domain-domain pairs, and the positive trend still maintains (median rho=0.107). Notably, after median data pooling in terms of presence frequency, the correlation coefficient is further augmented (median rho=0.504). Inspired by the above observations, we further screened high-frequency domain-domain pairs co-occurrent in ≥5% of the BGCs and depicted a macroscopic attention weight network containing 74 nodes and 2,048 edges (Fig. 3d; Supplemental Table S5-S6). First, highly weighted ‘KAsynt_C_assoc’- and ‘PP-binding’-targeting attention connections are enriched in the network, in consistent with their top-ranked importance scores. As an essential component of beta-ketoacyl synthase (KS), ‘KAsynt_C_assoc’ specifically exists in most PKSs (94.40%) [24]. And ‘PP-binding’ widely exists in both PKS and NRPS, accounting for 69.06% of the known BGCs [14, 24, 25]. The top-ranked attention connections also contain some other inter-domain associations with clear biosynthetic signatures. For example, the paired ‘AMP-binding’ and ‘Condensation’ (ranking 4th) represent the biosynthesis of NRPs [25]. The combination of ‘UbiA’ and ‘p450’ (ranking 28th) covers 19.23% of known terpene biosynthetic pathways [14] and contains an emerging UbiA-type of TCs [26, 27]. There are also domain-domain pairs possibly related to biosynthesis-assistant functions. For example, paired ‘ABC_membrane’ and ‘ABC_tran’ (ranking 3rd) represent the basic constitution of ATP-binding cassette (ABC) transporters [28], which might be responsible for the export of SMs across cellular membranes [29]. The ternary topology of ‘adh_short’, ‘adh_short_C2’ and ‘KR’ (ranking 16th, 17th and 24th), combined with the indirect interaction between ‘3Beta_HSD’ and ‘Epimerase’ mediated by ‘NAD_binding_4’ (ranking 23rd and 25th) collectively define canonical short-chain dehydrogenase/reductase (SDR) superfamily, which is crucial for SM tailoring [30].

At protein level, for each BGC we prioritized the protein with the highest incoming attention weight for investigation, whereas the other proteins served as control. These two groups were compared in terms of domain components by one-sided Fisher’s exact tests. Consequently, we found the protein-level attention weight can be a robust indicator of core enzyme since quite a few domains over-represented in the high attention group are of core biosynthetic relevance (Fig. 3e). For example, the combinations of (1) ‘ketoacyl-synt’, ‘Ketoacyl-synt_C’, ‘KAsynt_C_assoc’ and ‘Acyl_transf_1’, (2) ‘Condensation’, ‘AMP-binding’ and ‘AMP-binding_C’ and (3) ‘Terpene_syn_C_2’ are strongly indicative of PKSs [24], NRPSs [25] and partial TCs [31, 32], respectively.

In conclusion, the f-BGM-derived attention weights are of biological significance at both domain and protein levels. The decent interpretability could assist researchers to better understand single-domain and -protein importance, as well as inter-domain partnership of interested BGCs.

### f-BGM identifies core enzymes within BGCs in high accuracy

To benchmark the performance of f-BGM core enzyme identification model, a RF-based framework supplemented with Pfam domain features, which was commonly used for BGC-related binary classification [12, 13], was introduced as baseline (See ‘Methods and materials’). The benchmark tests involved seven major families of core enzymes with abundant BGC samples (≥30 BGCs) (Supplemental Table S1), where the training/validation/test sets were randomly assigned at a 4:1:1 ratio, and the performance metrics of 16 rounds of *de novo* experiments were averaged for algorithm evaluation. To adapt the BGC-level prediction of RF, the ORF-level probabilistic scores of f-BGM underwent max pooling for each BGC.

First, although the RF model is of naïve design, it still exhibits decent capability in core enzyme identification, where the average AUROCs and AUPRCs no less than 0.990 and 0.950 were widely observed (Fig. 4a; Supplemental Fig. S8). However, by investigating the challenging positive BGCs whose confidence scores lower than ≥2.5% of the negative BGCs, we noted RF might be less powerful in occasional cases, especially when deciphering chimeric BGCs with multiple core enzyme families (Fig. 4b). In contrast, f-BGM not only focuses on BGCs’ domain components, but also their inner associations (f-DLC) and sequence information (f-ESM2), which endow it with potential to process BGCs with more complex structures. As expected, a comparative analysis reveals that all the RF-resistant BGCs are prioritized by f-BGM when identifying most core enzyme families including PKS, NRPS, PT, PKS-NRPS, chimeric TS and PPPS (Fig. 4c), in consistent with its nearly perfect performance elucidated by AUPRCs (≥0.990 in average) (Fig. 4a). However, for the prediction of TC family, f-BGM fails to maintain the absolute superiority, where RF performs better on a subset of BGCs (11/30, 36.66%). Nevertheless, most comparative links (19/30, 63.33%) are still biased to f-BGM (Fig. 4d). Consequently, f-BGM achieves AUPRC improvement of 0.022 (0.973±0.024 vs 0.951±0.027) over RF. The consistent improvements of AUROC and F1-score confirm stable performance gains (Supplemental Fig. S8). Some representative examples prioritized by f-BGM over RF are illustrated in Supplemental Fig. S9.

**Figure 4.**
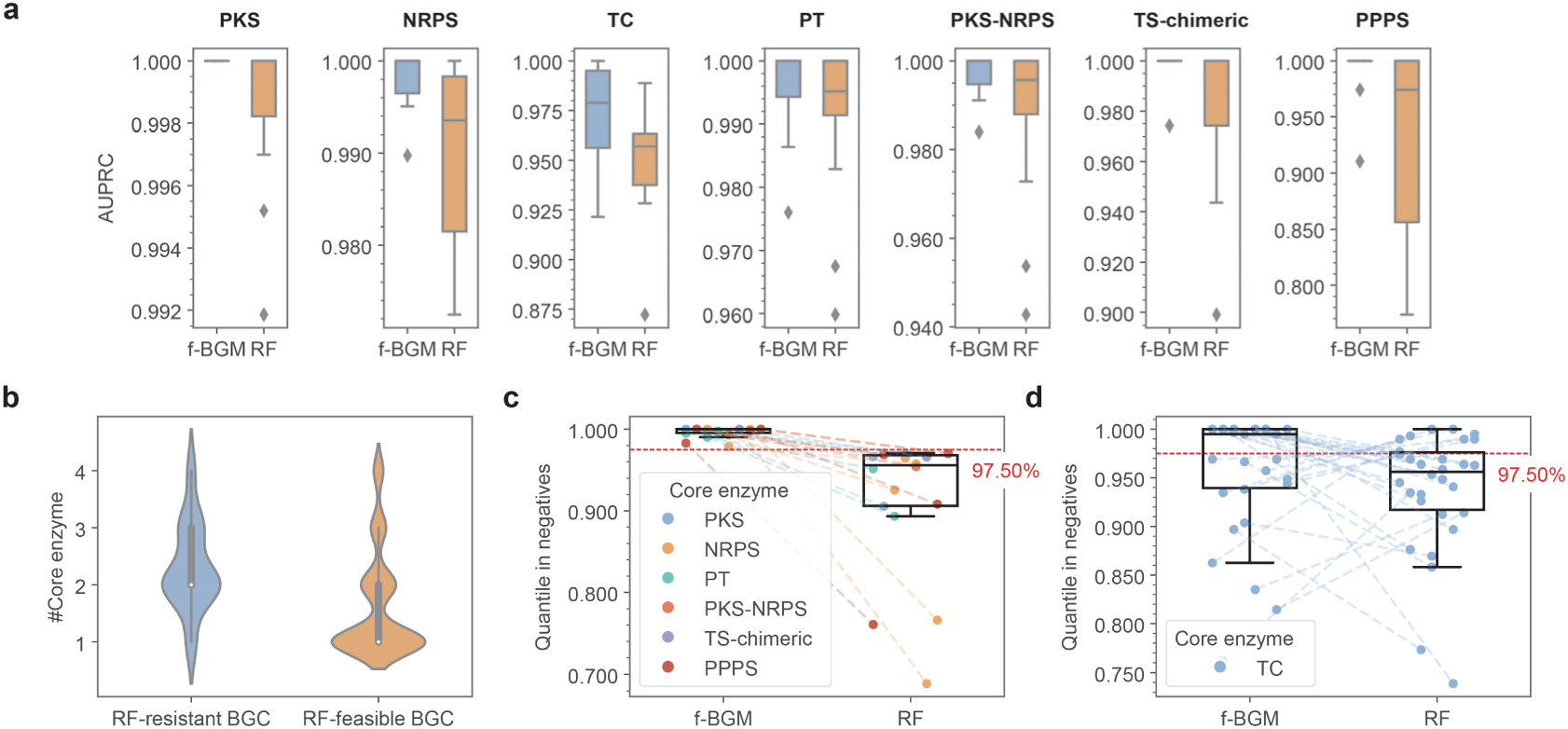
Performance of f-BGM in core enzyme identification benchmark tests. **a** Performance comparison between f-BGM and random forest (RF) in identifying seven major families of core enzymes with regard to AUPRC. **b** Distributional difference in core enzyme number between RF-resistant challenging (scored below ≥2.5% of negatives by RF) and RF-feasible BGCs. **c** Performance comparison between f-BGM and RF in identifying PKS, NRPS, PT, PKS-NRPS, chimeric TS and PPPS on challenging BGCs (scored below ≥2.5% of negatives) with regard to their relative ranks in negatives. Since all the BGCs are properly prioritized by f-BGM, only the BGCs underestimated by RF are plotted. **d** Performance comparison between f-BGM and RF in identifying TC on challenging BGCs with regard to their relative ranks in negatives. All the BGCs underestimated by any of f-BGM and RF are plotted.

At ORF level, f-BGM’s performance still maintains at an excellent level for most core enzymes (PKS, NRPS, PT, PKS-NRPS and chimeric TS). Despite the degraded AUPRCs for TC and PPPS identification relative to those at BGC level, the absolute values of 0.928 (±0.041) and 0.965 (±0.088) are still acceptable (Supplemental Fig. S10), confirming f-BGM’s capability in core enzyme localization beyond naïve presence-absence judgement.

The complete f-BGM framework was finally packaged as a Python-based command line toolkit to ensure its usability (See ‘Methods and materials’).

### Genome mining in marine fungi using f-BGM reveals BGCs out of antiSMASH-predefined rules

To adapt high-salinity and -pressure environments, marine fungi have evolved sophisticated biosynthetic pathways, representing an underexplored resource for SM discovery [33–35]. Using in-house genomic data of four marine fungal strains including *Aspergillus sclerotiorum* LZDX-33-4, *A. wentii* SX-4-1, *A. puulaauensis* F77, and *Talaromyces funiculosus* AN-49-2, we further conducted comparative genome mining research between f-BGM and the ubiquitously adopted rule-based method antiSMASH (version 7.1.0). For f-BGM, the putative BGCs were organized from the top highly scored ORFs at a relaxed ratio of 10%, combined with a false positive-controlling constraint of containing at least one ORF whose confidence score ≥0.3. antiSMASH was set to default ‘relaxed’ mode.

First, by comparing against the ORF number distribution of known BGCs, we noted f-BGM performs better on the determination of BGC borders, whereas antiSMASH is likely to overestimate BGCs’ genomic span (Fig. 5a). Then using least one common ORF as a criterion, we found f-BGM successfully hits 76.13% (236/310) of the antiSMASH-putative BGCs (Fig. 5b), confirming its basic functionality. We further focused on 25.70% (83/323) of the putative BGCs exclusively presented by f-BGM. First, 19.28% of these gene clusters are annotated with specific core enzymes, among which the enrichment of terpene biosynthetic potential (TC and chimeric TS) was observed (Fig. 5c). Indeed, unlike the relatively conserved core enzyme families such as PKS and NRPS, terpene synthases consist of phylogenetically divergent subtypes [36], bringing challenges for systematic extraction of detection rules. Here we prioritized a 4-ORF gene cluster C77 and a 6-ORF gene cluster C128 from the LZDX-33-4 genome for further investigation, of which the ORFs g4217 (TC) and g7212 (chimeric TS) were respectively identified as candidate core enzymes, in accordance with their highest importance defined by protein-level attention weights (Fig. 5d; Supplemental Fig. S11). For functional validation, these two gene clusters were heterologously expressed in *A. nidulans* (See ‘Supplemental experimental procedures’) [37]. Compared to negative control, four main products (**1**-**4**) and one product (**5**) were respectively observed in the C77- and C128-expressing strains via LC-/GC-MS analyses (Fig. 5e). Further NMR spectroscopy and optical rotation analysis elucidates **1-4** as sesquiterpenes and **5** as sesterfisherol (Fig. 5f; Supplemental Fig. S12-S32; Supplemental Table S7-S10). The remaining f-BGM-specific putative BGCs (80.72%) lack of usual core enzymes (Fig. 5c). To our best knowledge, such non-canonical instances always indicate sub-BGCs separated from the core regions [38–40]. Indeed, through BLAST analysis on protein sequences (Supplemental Fig. S33), we first noted that a 19-ORF gene cluster C123 from the AN-49-2 exhibits high homology with a validated sub-BGC (FunBGCs ID: FBGC00470) mediating the demethoxyviridin-biosynthesis (Supplemental Fig. S33b), which is featured with packed cytochrome P450 monooxygenase genes [39]. As for the other non-canonical gene clusters, despite the absence of direct homologous linkages with specific validated BGCs, ∼57.74% of their protein products share significant similarity (e-value≤1×10^−30^) with at least one protein in the whole FunBGCs dataset (Supplemental Fig. S33a). Further dimensionality reduction visualization on domain components illustrates the non-canonical putative BGCs are adjacent to non-canonical validated BGCs and prone to be sandwiched by canonical (validated/putative) BGCs and random sampled background (Fig. 5g), plausibly indicating their marginal biosynthetic potential. Therefore, f-BGM provides a broader perspective for discovering novel biosynthetic mechanisms.

**Figure 5.**
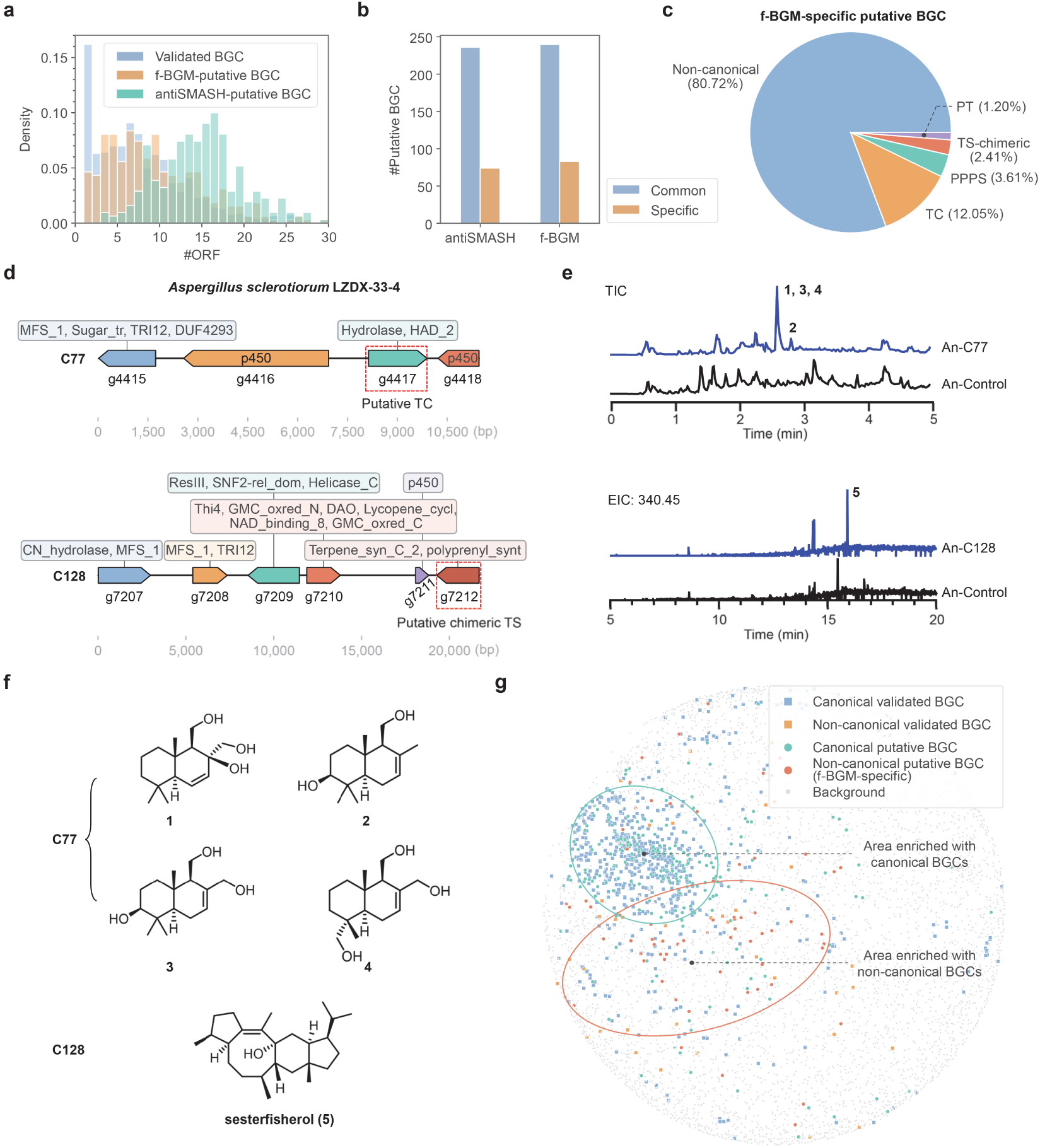
Comparative genome mining research in four marine fungi between f-BGM and antiSMASH. **a** Distributional difference in ORF number between f-BGM- and antiSMASH-putative BGCs, using validated BGCs as reference. **b** Number of gene clusters commonly and specifically predicted by f-BGM and antiSMASH. **c** Constitution of f-BGM-specific putative BGCs with regard to core enzyme families. The non-canonical part represents the gene clusters in absence of usual core enzyme families. **d** Prioritized gene clusters C77 and C128 from the LZDX-33-4 genome and their constitutions of Pfam domains. Both gene clusters are predicted of terpene biosynthetic potential, and the ORFs g4417 and g7212 are identified as putative core enzymes, respectively. **e** MS analysis reveals compounds **1**-**4** and **5** in *A. nidulans* heterologously expressed with C44 and C128, respectively. **f** Elucidated chemical structures of compounds **1**-**5**. **g** Domain component-based dimensionality reduction visualization indicates marginal biosynthetic potential of the non-canonical gene clusters exclusively predicted by f-BGM. Specifically, they exhibit clustering tendencies with non-canonical validated BGCs and tend to be sandwiched between canonical (validated/putative) BGCs and random sampled background.

## Discussion

Benefit by the accumulation of high-quality data of fungal BGCs, here we proposed an advanced deep learning framework f-BGM specifically for fungal genome mining. Different from the existing methods, f-BGM places more emphasis on capturing inter-domain and inter-gene associations via a self-attention-based model design. After rigorous evaluation through multiple benchmark tests, f-BGM was validated of SOTA performance in both (1) BGC detection in given genomes and (2) core enzyme identification in given BGCs. Besides the enhanced performance, its comprehensive interpretability in BGC detection is also remarkable. In spite some existing algorithms such as ClusterFinder [11] and GECCO [13] employ probabilistic modeling frameworks with explicit parameters (i.e., HMM and CRF), they are still less interpretable for the following reasons: (1) Both HMM and CRF simplify the cooperative relationship among multiple domains and ORFs within BGCs by first-order Markov property, where the hidden states (i.e., whether the domains or ORFs located in BGCs) are assumed associated with respective neighbor hidden states instead of broader receptive fields, which might introduce hypothesis-driven biases; (2) In GECCO, the CRF framework only uses linearly weighted domain features for ORF modeling. Therefore, its interpretability is only limited to single-domain importance revealed by the static post-training weights. In contrast, f-BGM’s architecture enables multi-level interpretability through dynamic calculation of attention weights. In this work, we presented the attention weights or attention weight-based importance scores are of biological significance no matter for domain-domain pairs, single domains or single proteins.

For further demonstration, we presented a purely f-BGM-driven workflow for marine fungal SM discovery. First, f-BGM recovers 76.13% of the gene clusters predicted by the rule-based method antiSMASH [10], positioning it as an alternative method for standard research. Not limited to this, f-BGM also detects a subset of gene clusters (25.70%) not annotated by antiSMASH, of which two gene clusters with candidate terpene synthases were prioritized and functionally validated by heterologous expression experiments. Moreover, there is also a larger part of non-canonical gene clusters without usual core enzymes. Methodologically, antiSMASH adopts a core gene-centered strategy combined with genomically flanking extension to detect BGCs, thus it is obviously powerless in locating such core enzyme-absent gene clusters. However, our result suggests the underestimated prevalence of such ‘non-canonical’ instances, which accounts for a promising proportion (20.74%, 67/323). More importantly, their homology with validated BGCs reveal by *in silico* analyses on member protein sequences and domain components confirms their plausibly biosynthetic roles rather than false positives. Overall, experimental and computational validation jointly demonstrates that f-BGM can push the boundaries of biosynthetic space exploration in fungi.

Another contribution of this work is the innovatively proposed architecture of f-DLC, which acquires the inter-domain co-occurrent relationship in local genomic contexts while considering domain-to-protein affiliations. Here the efficacy of f-DLC in BGC detection had been proved by ablation experiments on f-BGM. However, its applications can be further extended since it actually provides a fundamental representation methodology for functional gene clusters. For example, AI-driven prediction of SMs’ chemical structures [41] and bioactivities [42] based on genomic information has been preliminarily explored for bacteria, but such methods always organize BGCs into over-simplified representations such as purely domain-based feature vectors or token sequences without high-order interactions. By introducing f-DLC or its variants as foundation models, their performance is hopefully improved. Moreover, genomic contextualization for proteins has also been shown helpful for predicting general enzyme functions, elucidating *cis*-regulation relationship and so on [43], indicating extensive utility of f-DLC beyond the original genome mining purpose.

## Methods and materials

### Data source

The information of experimentally validated fungal BGCs including DNA sequences, member ORF locations and member protein sequences was retrieved from two existing databases, FunBGCs (downloaded on June 22, 2024) [14] and MIBiG (version 3.1) [7]. After deduplication on DNA sequences using Smith-Waterman algorithm, 735 FunBGCs- and 128 MIBiG-sourced BGCs were retained, respectively (Supplemental Table S2). FunBGCs was determined as the core dataset for f-BGM development due to its larger volume, whereas MIBiG served as the test set in a downstream cross-dataset BGC detection test.

EnsemblFungi (version 56) [44] and JGI MycoCosm (downloaded on July 3, 2024) [45] were considered as the primary data source of fungal genomes, where whole DNA sequences, genomic annotations and protein sequences of thousands of fungi strains identified by unique taxon IDs are curated. The whole genome data was used for (1) tracing BGCs’ genomic origins, (2) constructing negative gene cluster samples, (3) pretraining the f-DLC model and (4) fine-tuning the established ESM2 model (See below). For the (2)-(4) purpose, the data underwent strain-wise deduplication. Specifically, only the genome with most annotated ORFs per strain was retained, resulting in 985 EnsemblFungi- and 1,777 MycoCosm-sourced genomes, respectively (Supplemental Table S3).

The retrieved protein sequences were further annotated with Pfam domains [46, 47]. In detail, the information of 19,632 naturally conserved protein domains in HMM format was downloaded from the Pfam database (downloaded on March 7, 2024), and the *hmmscan* subprogram of Python package *PyHMMER* (version 0.8.1) [48, 49] was utilized to achieve HMM-based domain detection with an i-evalue cutoff 0.01.

### Tracing of BGC genomic origin

Due to the scarcity of genomic data annotated with experimentally validated BGCs, we tried to build BGC-genome linkages from scratch based on existing data. With zero tolerance to mismatches, the DNA sequences of 447/863 (51.80%) BGCs, were successfully traced to 344 contigs of 182 fungal genomes (Supplemental Table S4). The BGC-containing contigs enabled downstream benchmark tests of real BGC detection.

### Construction of negative gene clusters and synthetic genomic contexts

Two types of negative gene cluster samples, respectively termed as type-1 and type-2 negative samples, were constructed in this work. The type-1 negative samples are consecutive ORFs randomly sampled from fungal genomes in accordance with the ORF number distribution of known BGCs, which represent the information of genomic background. To avoid mis-inclusion of real BGCs, the cosine similarity of domain components between the type-1 negative samples and the known BGCs were constrained <0.5. In contrast, the type-2 negative samples constitute a challenging set with higher BGC similarity, which were constructed under a homologous replacement strategy originally proposed by DeepBGC [12]. Specifically, given a query BGC and a reference genome, each member ORF of the query BGC was randomly replaced by one of the most similar ORFs ranked in the top 1% of the reference genome. In addition, the selected negative ORFs were required from different contigs so that even though the constructed samples are of higher BGC similarity, they belong to unreal genomic context, confirming their negative roles. The type-2 negative samples were constrained by a maximal BGC similarity of 0.75. For all the negative samples, the member ORFs were required to be unique and located out of known BGC-containing contigs to avoid training-to-test information leakage.

By performing random strand inversion, shuffling and concatenation on the BGCs and the negative gene cluster samples, we further constructed synthetic genomic contexts to enable model training and downstream benchmark tests of simulated BGC detection. In this work two construction schemes were applied: (1) BGCs + only type-1 negative samples at a 1:16 ratio. This is the default configuration in this manuscript unless otherwise specified; (2) BGCs + type-1 negative samples + type-2 negative samples at a 1:12:4 ratio. This scheme was only applied in a challenging version of simulated BGC detection task. The synthetic genomic contexts were required with a proper number of ORFs ranging from 128 to 144. Moreover, to ensure the diversity of gene arrangement, abundant genomic contexts equaling half the number of gene cluster samples were constructed.

### Design and pretraining of f-DLC model

f-DLC is a specially designed language model for capturing inter-domain locally co-occurrent relationship in fungal genome and thereby generating meaningful ORF-level representation embeddings for given consecutive ORFs. f-DLC linearizes consecutive ORFs with a hierarchical domain-to-protein structure into a token sequence *S*:

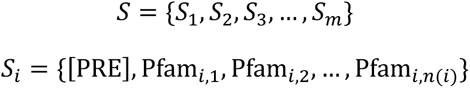

where *S*_*i*_ denotes the token sub-sequence of the *i*-th ORF, [PRE] is the prefix token to mark the beginning of sub-sequence, Pfam_*i,j*_ represents the *j*-th Pfam domain token identified in the protein encoded by the *i*-th ORF, *m* and *n*(*i*) are the ORF number of input and the domain number of *i*-th ORF, respectively. The tokens are further translated into 72-D embeddings consisting of three components: (1) Token-specific 64-D learnable embeddings; (2) 4-D embeddings linearly transformed from the information of domains’ relative location within proteins; (3) 4-D embeddings linearly transformed from the domain enrichment information in BGCs. It should be noted that f-DLC was not only designed for BGCs but also general genomic contexts, therefore the third embedding component encoding domain enrichment information was temporarily masked during the pretraining process and only enabled when establishing the downstream f-BGM model (See below). The f-DLC model contains four TFEs equipped with self-attention mechanism and achieves layer-specific inter-token information interaction by configuring different attention masks (Supplemental Fig. S1). Specifically, the first and the second TFEs respectively calculate on the inter-domain attention connections within single protein and multiple proteins, ensuring multi-scale domain-level information interaction. In addition, real-time domain-to-protein information aggregation is achieved by the attention mechanism from domain tokens to corresponding prefix tokens. The third TFE serves as a readout module for upstream extracted features, thus only the domain-to-protein attention connections are remained. The protein-level (ORF-level) embeddings for downstream use are exactly derived from this layer. The fourth TFE only serves the BERT-style self-supervised pretraining [50] of f-DLC, where the domain tokens of input sequences are partially masked, and then the model is guided to recover the masked tokens by an additional linear multi-classification header. For this purpose, a reversed attention mechanism is applied on the upstream generated protein-level embeddings to achieve information passing from prefix tokens to the subordinate domain tokens. During the pretraining process, an Adam optimizer with a learning rate of 5×10^−4^ was used to minimize the cross-entropy loss between the predicted and true domain tokens with masks. The training iteration number was set to 262,144 for exhausted parameter optimization (Supplemental Fig. S2). For each training iteration, 64 genomic windows containing 1-26 consecutive ORFs were randomly sampled from fungal genomes and fed into the model, with 15% of domain tokens masked. The genomic window size follows the ORF number range of known BGCs.

### Modifications and fine-tuning on ESM2 model

ESM2 is a well-known protein language model [22]. Theoretically, it was also trained following the BERT protocol by partially masking and recovering protein sequences [50], thereby it can capture the inter-residue associations and generate effective protein representation embeddings. Here the pretrained ESM2 model (150M-parameter version) was introduced with two modifications: (1) For downstream parameter efficiency, the dimensionality of original output representation embeddings was reduced from 640-D to 64-D by an additional linear layer; (2) Considering the original ESM2 model was trained on a universal protein dataset, fungi-specific fine-tuning was performed to specialize the model function. The fine-tuned model is termed as f-ESM2 in this manuscript. For memory-footprint optimization, here the parameters of original ESM2 were frozen and only the linear layer for dimensionality reduction was configured trainable. The fine-tuning process lasted for 2,048 iterations (Adam optimizer, learning rate=1×10^−3^). For each iteration, 32 protein sequences with a mask ratio of 15% were randomly sampled from the fungal genomes to constitute a training batch. The input sequences were required ≤1,024 amino acids, otherwise 1,024-aa subsequences would be randomly sliced out as representatives. However, when using f-ESM2 for formal inference (i.e., when establishing f-BGM), the overlength proteins were processed in another way to preserve the complete sequence information, specifically, fragmented into consecutive 1,024-aa subsequences with an overlap ratio of 75%, and the final representation embeddings were calculated by averaging the sub-embeddings.

### Establishment of f-BGM

Both f-DLC and f-ESM2 were integrated into f-BGM but with some modifications: (1) To reduce computation complexity, only the domain tokens existing in at least one BGCs in the training set were retained for f-DLC; (2) The domain enrichment information in BGCs was integrated into the input embeddings of f-DLC for semantic augmentation. Specifically, the presence frequency of each domain was compared between the training BGCs and background using one-sided Fisher’s exact test. The resulting p-values were ranked, min-max normalized across domains and further projected into 4-D embeddings by a trainable linear layer; (3) The f-ESM2 model is set to inference mode without gradient computation.

The BGC detection model was trained on the synthetic genomic contexts by minimizing the cross-entropy loss between output ORF-level probabilistic scores and true binary labels of BGC membership (Adam optimizer, learning rate=5×10^−4^). For each iteration, 64 genomic contexts were randomly selected for model training, meanwhile the validation set was used to monitor model convergence and trigger the early stopping mechanism to avoid overfitting. Specifically, if the performance metrics on validation data were no longer optimized for 64 iterations, then the training process would be interrupted. For the model training in absence of the type-2 negative gene cluster samples, the average recall at top ratios of 1%, 5% and 10% (See below) on the real genomic contigs was referred for early stopping. Otherwise, the AUPRC on the synthetic genomic contexts was used due to potential distributional difference between the type-2 negative samples and real-world data.

The optimization target of f-BGM core enzyme identification model was fitting ORF-level probabilistic scores with binary labels of whether the ORFs encoding target core enzymes. The training batch of each iteration consisted of 8 positive BGCs and 8 negative BGCs. And the validation AUPRC was monitored for early stopping. The other configurations including loss function, parameter optimizer, learning rate and patience for early stopping remain consistent with those of the BGC detection model.

The f-BGM model was entirely established under the deep learning framework PyTorch (version 2.3.1) [51] on an NVIDIA RTX A5500 GPU.

### Strand inversion-invariance of BGC detection model

To distinguish the proteins encoded by different DNA strands, positive and negative strands are always artificially defined for genomic contigs. However, for the BGC detection problem, ORFs’ confidences as BGC members should be independent with the strand definition. In other words, BGC detection result should be constant regardless of the directions of ORF arrangement, namely the strand inversion-invariance. This section will present the strand inversion-invariance of the two main modules of BGC detection model, namely SRM and LRM.

The SRM adopts a multi-TFE design. The TFE architecture consists of a self-attention block and a linear block. Under the definition of self-attention mechanism, the TFE-output embedding at each position only depends on the corresponding input embedding and its interaction partner embeddings:

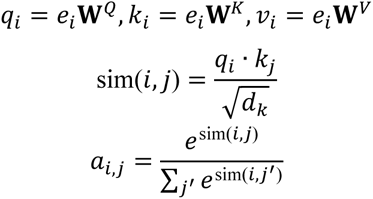

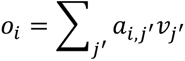

where *q*_*i*_, *k*_*i*_ and *v*_*i*_ represents the linearly transformed embeddings of the *i*-th input embedding *e*_*i*_ using learnable matrices **W**^*Q*^, **W**^*K*^and **W**^*V*^, *a*_*i,j*_ is the softmax-defined attention weight between the *i*-th and *j*-th tokens, and *o*_*i*_ is the final output embedding at position *i*. In SRM, the TFEs define inter-domain, domain-to-protein and inter-protein interactions in multi-scale neighborhoods. Since the relative positioning among the ORFs is constant, the interaction partners of each domain and protein are independent from the strand definition. Therefore, the strand inversion-invariance of SRM obviously holds.

The LRM applies traditional LSTM architecture to achieve global message passing. By default, the LSTM neural network cannot satisfy the inversion-invariant requirement, therefore we modified it with a novel forward propagation strategy. Specifically, given an embedding sequence *E* as input, the output embedding at position *i*, denoted as *e*_*i*_, is calculated as follows: (1) Pass *E* into LSTM in positive direction, and denote the output embedding at position *i* as *e*_*i*_^+^; (2) Pass *E* into LSTM in negative direction, and denote the output embedding at original position *i* as *e*_*i*_^−^; (3) Calculate the final *e*_*i*_ = *e*_*i*_^+^ + *e*_*i*_^−^. In this way, for the inversed *E*, the output embedding at original position *i*, denoted as, 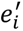, will satisfies:

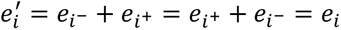

The strand inversion-invariance is thus proved.

### Baseline algorithms

ClusterFinder [11], DeepBGC [12], GECCO [13] and TOUCAN [23] were selected as baseline algorithms in downstream BGC detection benchmark tests. Here the baselines were reimplemented following the original literatures for convenience of customized training. The following text will briefly introduce basic principles of the baselines and key points for their reimplementation.

ClusterFinder uses HMM to model genomic contexts at domain level, where the domains are regarded as observation states, and the binary labels of whether the domains belong to BGCs are regarded as hidden states [11]. In this way, given a genomic context in form of domain sequence, the hidden states can be deduced by calculating posterior probabilities conditioning the observations. In this work the parameters of ClusterFinder, namely start probabilities, state transition matrices and emission probabilities of HMM, were determined by simple counting statistics on the training synthetic genomic contexts [11]. The Python package *hmmlearn* (version 0.2.8) was used for HMM implementation.

DeepBGC is the first deep learning algorithm for genome mining, which employs a Bi-LSTM network in combination with a domain-level vectorization strategy termed as Pfam2Vec [12]. The Pfam2Vec strategy regards genomic contigs and the subordinate Pfam domains as sentences and words, respectively. In this way, the well-known Word2Vec algorithm [52] originally designed for word vectorization in natural language could be applied for extension. The resulting embeddings could reflect inter-domain associations in fixed range (specified by a hyperparameter ‘window size’), thus are informative for BGC detection. In the DeepBGC pipeline, the input genomic contexts are first translated into sequences of Pfam2Vec embeddings, appended with two flag-bits representing protein borders. Then the embeddings are directly fed into the Bi-LSTM module for global message passing. After further projection by a sigmoid-gated linear header, the confidence scores of BGC membership will be generated at domain level. Here the Pfam2Vec embeddings were trained from scratch based on the fungal genomes, where the Python package *gensim* (version 4.2.0) was used for Word2Vec implementation. The hyperparameters including embedding dimensionality and window size were set to 128 and 8, respectively. The DeepBGC architecture was reimplemented under the PyTorch framework [51] and trained on the synthetic genomic contexts (Adam optimizer, learning rate=5×10^−4^, batch size=64). The number of training iterations was determined by validation-based early stopping mechanism.

Here for both ClusterFinder and DeepBGC, the output domain-level confidence scores were finally averaged into ORF level as the result. For the ORFs without identified domains, the confidence scores were linearly interpolated by flanking ORFs.

GECCO directly models genomic contexts at ORF level by linear CRF, where each ORF is characterized by presence or absence of the feature domains over- or under-represented in BGCs [13]. Here the feature domains were screened by two-sided Fisher’s exact tests as the original literature suggested [13], with a standard p-value cutoff of 0.05. The CRF model training was still based on the synthetic genomic contexts. For further improvements, we additionally optimized the regularization factors c1 and c2 through an exhausted grid search evaluated on the validation set, where the c1, c2 values of 0, 0.125, 0.25, 0.5, 1, 2 and 4 were enumerated (Supplemental Table S11). The CRF models were implemented using the Python package *sklearn-crfsuite* (version 0.9.10).

Distinct from the above sequence-based algorithms, TOUCAN only uses a binary classification model to detect BGCs. In detail, TOUCAN introduces three types of features to profile gene cluster samples: (1) K-mer protein subsequences; (2) High-specificity domains exclusively or largely exist in BGCs. Here these domains were redefined in a more quantizable manner by one-sided Fisher’s exact tests with a p-value cutoff 0.05; (3) GO functional annotations deduced by homologous proteins in the well-annotated SwissProt database [53]. With bag-of-words vectorization on these features, a classification model built from traditional machine learning algorithms such as RF, SVM and LR can be trained to distinguish BGCs from negative gene cluster samples. When performing BGC detection, the model will be used on a genomic window sliding through the input genomic contexts, and thereby assign confidence scores for all the ORFs after full-range scanning. In this work, we reimplemented RF-based TOUCAN models by means of the Python package *scikit-learn* (version 1.0.2) [54]. The most proper k-mer length of the protein subsequence features was optimized based on validation AUPRC (Supplemental Table S11). As for the other hyperparameters, the decision tree number of RF was set to standard 256, and the window for BGC detection was set with 5,000-aa coverage and 2,500-aa sliding step. The window size mimics the median length of known BGCs.

For the core enzyme identification problem, an algorithmic framework commonly used for BGC-related binary classification, which combines RF with Pfam domain-based bag-of-words vectorization [12, 13], was reimplemented by *scikit-learn* [54] as baseline. The models accommodate 256 decision trees to fit BGC-level presence-absence labels of target core enzymes.

### Cross-SM-class BGC detection benchmark tests

BGCs’ representative SM products of the FunBGCs dataset were clustered by MACCSKeys-based K-means algorithm and an inertia-based elbow method was used to find the most proper cluster number. As a result, six BGC subsets with SM chemical diversity were determined (Supplemental Fig. S3a-b). The highest-frequency core enzymes of each subset are illustrated in Supplemental Fig. S3c. In the cross-SM-class benchmark tests, algorithms’ prediction results were generated in a rotational cross-validation manner to cover all six subsets. To be more specific, each subset is assigned as the test set in turn, meanwhile the other subsets jointly constitute the training/validation sets. The prediction results on each subset were further aggregated for systematic evaluation. Considering the BGC number of each subset is imbalanced (Supplemental Fig. S3c), here we applied a balanced subsampling strategy to avoid biases towards the larger subsets. In more detail, the final performance metrics were averaged over 512 subsampling rounds. For each round, equal number of genomic contexts were randomly sampled from each subset and then merged for single-round metric calculation. Here the number of subsampled genomic contexts per round was set to 2 and 4 for the real BGC detection task and the simulated BGC detection task, respectively

### Performance metrics

Algorithm performance for the BGC detection problem was evaluated at ORF level. For the real BGC detection task, we only focused on the recall (also termed as sensitivity) of known positives due to the uncertainty of negatives. By defining the predictive positives as the top highly scored ORFs at specified ratios, the recalls were calculated as follows:

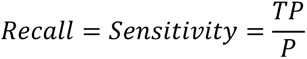

where *TP* and *P* represent the number of true positives and total positives, respectively.

For the simulated BGC detection task with clearly labeled negatives, the confusion matrix can be fully determined by a prediction score threshold differentiating the predictive positives and negatives, which enables the calculation of specificity and precision:

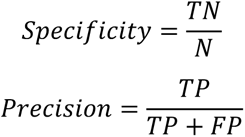

where *TN*, *FP* and *N* represent the number of true negatives, false positives and total negatives, respectively. We further depicted ROC curve reflecting the relationship between 1-specificity and sensitivity, and P-R curve focusing on the relationship between recall and precision, with the prediction score threshold increasing from 0 to 1. The metrics AUROC and AUPRC were calculated for systematic performance evaluation. In addition, the data points of P-R curve were enumerated to calculate F1-scores, and the best value was retained as another performance metric:

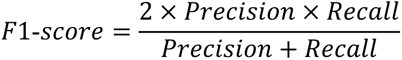

For all metrics values closer to 1 were preferred.

As for the core enzyme identification problem, the BGCs with and without target core enzymes were regarded as positives and negatives, respectively. Then the metrics AUROC, AUPRC and F1-score could be calculated at BGC level for algorithm comparison. It should be noted that the BGC-level prediction scores of f-BGM were processed from original ORF-level scores by max pooling, which sacrifices the locational information of core enzymes.

During the entire process from model development to evaluation, quite a few steps introduced randomness, such as the construction of negative gene cluster samples and synthetic genomic contexts, the assignment of training/validation/test sets, the multi-round subsampling-based evaluation and so on. Therefore, to counteract the randomness-induced variations, all the performance metrics were averaged over 16 rounds of *de novo* experiments. Moreover, since the training of deep learning-based algorithms is more sensitive to random seeds, the probabilistic scores of f-BGM and DeepBGC used for evaluation were averaged from four independently trained models.

### Normalization of attention weights

Due to the fixed sum constraint of softmax definition, the spatial expansion of TFE’s receptive field will induce attenuation of individual attention weights in random expectation. Therefore, to ensure comparability among the gene cluster samples containing different number of domains or proteins, the attention weights of each sample were normalized by a robust version of z-score:

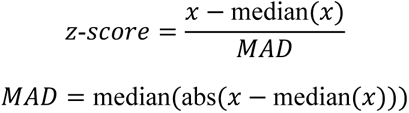

where *x* represents the full list of values to be normalized.

### Development of f-BGM toolkit

To extend the usability of f-BGM, we developed a Python-based command-line toolkit, where three main functions including (1) BGC detection (using the models built under in-distribution scheme), (2) core enzyme identification and (3) attention weight visualization are provided. To improve predictive robustness, all the models built in 16 rounds of *de novo* experiments are employed to generate averaged results. The toolkit has been released at https://github.com/bingozhyr/f-BGM.

## Acknowledgements

This study was supported by the National Key R&D Program of China (2022YFC2804900), the Beijing Natural Science Foundation (7244465, 7242195), the Beijing AI Health Cultivation Project (Z221100003522022), the National Natural Science Foundation of China (22277006, 82173733, 82473836), and the Ningbo Key Science and Technology Development Program (2022Z144).

## Notes

https://github.com/bingozhyr/f-BGM

https://figshare.com/articles/software/f-BGM/29396423

### Competing Interest Statement

The authors have declared no competing interest.

## References

1. Patridge, E., et al., An analysis of FDA-approved drugs: natural products and their derivatives. Drug Discov Today, 2016. 21(2): p. 204–7.

2. Yip, D.W. and V. Gerriets, Penicillin, in StatPearls. 2025: Treasure Island (FL) ineligible companies. Disclosure: Valerie Gerriets declares no relevant financial relationships with ineligible companies.

3. Patocka, J., et al., Cyclosporine A: Chemistry and Toxicity - A Review. Curr Med Chem, 2021. 28(20): p. 3925–3934.

4. Adams, S.P., et al., Pravastatin for lowering lipids. Cochrane Database Syst Rev, 2023. 9(9): p. CD013673.

5. Cohen, J.A., et al., Oral fingolimod or intramuscular interferon for relapsing multiple sclerosis. N Engl J Med, 2010. 362(5): p. 402–15.

6. Aris, P., et al., Griseofulvin: An Updated Overview of Old and Current Knowledge. Molecules, 2022. 27(20).

7. Terlouw, B.R., et al., MIBiG 3.0: a community-driven effort to annotate experimentally validated biosynthetic gene clusters. Nucleic Acids Res, 2023. 51(D1): p. D603–D610.

8. Harvey, A.L., R. Edrada-Ebel, and R.J. Quinn, The re-emergence of natural products for drug discovery in the genomics era. Nat Rev Drug Discov, 2015. 14(2): p. 111–29.

9. Chiang, C.Y., M. Ohashi, and Y. Tang, Deciphering chemical logic of fungal natural product biosynthesis through heterologous expression and genome mining. Natural Product Reports, 2023. 40(1): p. 89–127.

10. Blin, K., et al., antiSMASH 7.0: new and improved predictions for detection, regulation, chemical structures and visualisation. Nucleic Acids Res, 2023. 51(W1): p. W46–W50.

11. Cimermancic, P., et al., Insights into secondary metabolism from a global analysis of prokaryotic biosynthetic gene clusters. Cell, 2014. 158(2): p. 412–421.

12. Hannigan, G.D., et al., A deep learning genome-mining strategy for biosynthetic gene cluster prediction. Nucleic Acids Res, 2019. 47(18): p. e110.

13. Carroll, L.M., et al., Accurate de novo identification of biosynthetic gene clusters with GECCO. bioRxiv, 2021: p. 2021.05.03.442509.

14. Tang, J. and Y. Matsuda, Discovery of fungal onoceroid triterpenoids through domainless enzyme-targeted global genome mining. Nature Communications, 2024. 15(1).

15. Vaswani, A., et al. Attention Is All You Need. 2017. arXiv:1706.03762 DOI: 10.48550/arXiv.1706.03762.

16. Jumper, J., et al., Highly accurate protein structure prediction with AlphaFold. Nature, 2021. 596(7873): p. 583–589.

17. Shen, T., et al., Accurate RNA 3D structure prediction using a language model-based deep learning approach. Nat Methods, 2024. 21(12): p. 2287–2298.

18. Zhang, P., et al., DeepMGT-DTI: Transformer network incorporating multilayer graph information for Drug-Target interaction prediction. Comput Biol Med, 2022. 142: p. 105214.

19. Wang, J.K., et al., Multi-constraint molecular generation based on conditional transformer, knowledge distillation and reinforcement learning. Nature Machine Intelligence, 2021. 3(10): p. 914–922.

20. Hao, M., et al., Large-scale foundation model on single-cell transcriptomics. Nat Methods, 2024. 21(8): p. 1481–1491.

21. Qiang, B., et al., Bridging the gap between chemical reaction pretraining and conditional molecule generation with a unified model. Nature Machine Intelligence, 2023. 5(12): p. 1476–1485.

22. Lin, Z., et al., Evolutionary-scale prediction of atomic-level protein structure with a language model. Science, 2023. 379(6637): p. 1123–1130.

23. Almeida, H., et al., TOUCAN: a framework for fungal biosynthetic gene cluster discovery. NAR Genom Bioinform, 2020. 2(4): p. lqaa098.

24. Cox, R.J. and T.J. Simpson, Fungal type I polyketide synthases. Methods Enzymol, 2009. 459: p. 49–78.

25. Miller, B.R. and A.M. Gulick, Structural Biology of Nonribosomal Peptide Synthetases. Methods Mol Biol, 2016. 1401: p. 3–29.

26. Yang, Y.L., et al., Discovery and Characterization of a New Family of Diterpene Cyclases in Bacteria and Fungi. Angew Chem Int Ed Engl, 2017. 56(17): p. 4749–4752.

27. Li, X.L., et al., Rapid discovery and functional characterization of diterpene synthases from basidiomycete fungi by genome mining. Fungal Genet Biol, 2019. 128: p. 36–42.

28. Thomas, C. and R. Tampe, Structural and Mechanistic Principles of ABC Transporters. Annu Rev Biochem, 2020. 89: p. 605–636.

29. Viglas, J. and P. Olejnikova, An update on ABC transporters of filamentous fungi - from physiological substrates to xenobiotics. Microbiol Res, 2021. 246: p. 126684.

30. Persson, B. and Y. Kallberg, Classification and nomenclature of the superfamily of short-chain dehydrogenases/reductases (SDRs). Chem Biol Interact, 2013. 202(1-3): p. 111–5.

31. Yan, Y., et al., Resistance-gene-directed discovery of a natural-product herbicide with a new mode of action. Nature, 2018. 559(7714): p. 415–418.

32. Han, H., et al., Functional Characterization of Sesquiterpene Synthases and P450 Enzymes in Flammulina velutipes for Biosynthesis of Spiro [4.5] Decane Terpene. J Agric Food Chem, 2024.

33. Liu, M., et al., Potential of marine natural products against drug-resistant bacterial infections. Lancet Infect Dis, 2019. 19(7): p. e237–e245.

34. Xu, Z.L., et al., Equisetin is an anti-obesity candidate through targeting 11β-HSD1. Acta Pharmaceutica Sinica B, 2022. 12(5): p. 2358–2373.

35. Rao, Y., et al., *Identification of a natural PLA2 inhibitor from the marine fungus Aspergillus sp. c1* for MAFLD treatment that suppressed lipotoxicity by inhibiting the IRE-1alpha/XBP-1s axis and JNK signaling. Acta Pharm Sin B, 2024. 14(1): p. 304–318.

36. Luo, P., et al., Biosynthesis of fungal terpenoids. Nat Prod Rep, 2024. 41(5): p. 748–783.

37. Chiang, Y.M., et al., Development of Genetic Dereplication Strains in Aspergillus nidulans Results in the Discovery of Aspercryptin. Angew Chem Int Ed Engl, 2016. 55(5): p. 1662–5.

38. de Wit, P.J., et al., The genomes of the fungal plant pathogens Cladosporium fulvum and Dothistroma septosporum reveal adaptation to different hosts and lifestyles but also signatures of common ancestry. PLoS Genet, 2012. 8(11): p. e1003088.

39. Wang, G.Q., et al., Biosynthetic pathway for furanosteroid demethoxyviridin and identification of an unusual pregnane side-chain cleavage. Nat Commun, 2018. 9(1): p. 1838.

40. Zhao, S., et al., Elucidation of Palmarumycin Spirobisnaphthalene Biosynthesis Reveals a Set of Previously Unrecognized Oxidases and Reductases. Angew Chem Int Ed Engl, 2024. 63(23): p. e202401979.

41. Xu, T., et al., *DeepSeMS:* a large language model reveals hidden biosynthetic potential of the global ocean microbiome. bioRxiv, 2025: p. 2025.03.02.641084.

42. Walker, A.S. and J. Clardy, A Machine Learning Bioinformatics Method to Predict Biological Activity from Biosynthetic Gene Clusters. J Chem Inf Model, 2021. 61(6): p. 2560–2571.

43. Hwang, Y., et al., Genomic language model predicts protein co-regulation and function. Nat Commun, 2024. 15(1): p. 2880.

44. Martin, F.J., et al., Ensembl 2023. Nucleic Acids Res, 2023. 51(D1): p. D933–D941.

45. Grigoriev, I.V., et al., MycoCosm portal: gearing up for 1000 fungal genomes. Nucleic Acids Res, 2014. 42(Database issue): p. D699–704.

46. Mistry, J., et al., Pfam: The protein families database in 2021. Nucleic Acids Res, 2021. 49(D1): p. D412–D419.

47. Paysan-Lafosse, T., et al., InterPro in 2022. Nucleic Acids Res, 2023. 51(D1): p. D418–D427.

48. Eddy, S.R., Accelerated Profile HMM Searches. PLoS Comput Biol, 2011. 7(10): p. e1002195.

49. Larralde, M. and G. Zeller, PyHMMER: a Python library binding to HMMER for efficient sequence analysis. Bioinformatics, 2023. 39(5).

50. Devlin, J., et al. BERT: Pre-training of Deep Bidirectional Transformers for Language Understanding. 2018. arXiv:1810.04805 DOI: 10.48550/arXiv.1810.04805.

51. Paszke, A., et al. *PyTorch:* An Imperative Style, High-Performance Deep Learning Library. 2019. arXiv:1912.01703 DOI: 10.48550/arXiv.1912.01703.

52. Mikolov, T., et al. Efficient Estimation of Word Representations in Vector Space. 2013. arXiv:1301.3781 DOI: 10.48550/arXiv.1301.3781.

53. UniProt, C., UniProt: the Universal Protein Knowledgebase in 2025. Nucleic Acids Res, 2025. 53(D1): p. D609–D617.

54. Pedregosa, F., et al., Scikit-learn: Machine Learning in Python. 2011. 12: p. 2825–2830.

