## Supplemental information for "f-BGM enables fungi-specific genome mining in high accuracy and interpretability"

**Supplemental experimental procedures**

**General methods and materials**

Optical rotations were measured on an Autopol III automatic polarimeter. The nuclear magnetic resonance (NMR) data were acquired on 500 MHz or 600 MHz Bruker FTNMR spectrometer using DMSO-*d_6_* or MeOH-*d_4_* as an internal standard. Structural assignments were made with additional information from COSY, HSQC, HMBC, and NOESY experiments. The chromatographic substrates included Sephadex LH20 (GE Healthcare Bio-Sciences AB, Sweden) and ODS-A (12 nm, S-50 μm; YMC Co., LTD., Japan). High performance liquid chromatography (HPLC) was performed using an Alltech 426 pump equipped with an Alltech UVIS-200 detector (210 nm) and a semipreparative reversed-phase column (YMC-packed, C18, 5 μM, 20 mm × 250 mm).

Analytical grade chemicals and reagents were purchased from Fischer, Beijing Tong Guang Fine Chemicals Company (Beijing, China), and Beijing Chemical Works. HPLC grade reagents were purchased from Sigma. General molecular biology kits were used according to the manufacturer’s protocols. Analytical PCR was performed using Taq polymerase (Mei5 Biotechnology Co. Ltd) and preparative PCR for cloning procedures was performed using Phanta ® Max Super-Fidelity DNA polymerase (Vazyme Biotech Co., Ltd.). Restriction endonucleases were purchased from NEB (New England Biolabs). Yatalase was purchased from Takara and lysing enzymes were purchased from Sigma.

**Strains, media and culture conditions**

The marine fungus *Aspergillus sclerotiorum* LZDX-33-4 was grown on PDA (potato dextrose agar) with sea salt plates at 28 °C. For [mycelia](D:/%E8%BD%AF%E4%BB%B6%E5%AE%89%E8%A3%85%E5%8C%85/LenovoSoftstore/Install/wangyiyoudaocidian/8.10.3.0/resultui/html/index.html#/javascript:;) growth, the strain was cultured in PDB (potato dextrose broth) with sea salt medium at 28 °C for 2 days.

*Aspergillus* *nidulans* LO8030 were used for heterologous expression. The transformants (Table S14) and wild-type strain were grown on GMM medium (1.0% glucose, 50 mL/L 20 × nitrate salt solution, 1 mL/L trace element solution, 1.6% agar) at 37 °C for conidia production. For conidia germination, the strain was cultured in 30 mL LMM medium (1.0% glucose, 50 mL/L 20 × nitrate salt solution, 1 mL/L trace element solution, 0.5% yeast extract) at 28 °C with 150 rpm shaking for 12-16 h. The 20 × nitrate salt solution comprises (w/v) 12% NaNO_3_, 1.04% KCl, 1.04% MgSO_4_∙7H_2_O, and 3.04% KH_2_PO_4_. The trace element solution contains (w/v) 2.2% ZnSO_4_∙7H_2_O, 1.1% H_3_BO_3_, 0.5% MnCl_2_∙4H_2_O, 0.16% FeSO_4_∙7H_2_O, 0.16% CoCl_2_∙5H_2_O, 0.16% CuSO_4_∙5H_2_O, 0.11% (NH_4_)_6_Mo_7_O_24_∙4H_2_O, and 5% Na_4_EDTA.

*Escherichia coli* DH5α which was used for heat shock transformation, was cultured in lysogeny broth (LB) medium (1% NaCl, 1% tryptone, and 0.5% yeast extract) at 37 °C. Ampicillin (100 μg/mL) was added to the LB agar plates for selection.

**Genomic DNA isolation**

The mycelia of *Aspergillus sclerotiorum* LZDX-33-4 and *A. nidulans* were dried on filter paper and collected in 2 mL Eppendorf tubes. Glass beads (2.85 mm in diameter) and 400 μL of LETS buffer (10 mM Tris-HCl pH 8.0, 20 mM EDTA pH 8.0, 0.5% SDS, and 0.1 M LiCl) were added to the tubes. After vigorous mixing for 1 min, 300 μL LETS buffer were added. The solution was then treated with 700 μL phenol: chloroform: isoamyl alcohol (25: 24: 1). Genomic DNA was precipitated by addition of 900 μL of absolute EtOH. After centrifugation at 13,000 rpm for 30 min and washing with 70% EtOH, the obtained DNA was dissolved in 30 μL distilled H_2_O.

**PCR amplification, molecular cloning, plasmid construction, and heterologous expression in *A. nidulans***

Primers and plasmids used in this study are listed in Table S12 and Table S13, respectively. All primers were synthesized by Genewiz (Azanta Inc., China). PCR amplification was carried out by using Phanta ® Max Super-Fidelity DNA Polymerase (Vazyme Biotech Co.,Ltd. Nanjing, China). PCR thermal profiles were set following the manufacturer’s instructions. The gene fragments were cloned from gDNA of *A. sclerotiorum* LZDX-33-4. The plasmids for heterologous expression were constructed via homologous recombination in *E. coli* DH5α. For construction of the plasmids for heterologous expression in *A. nidulans*, the DNA fragments were obtained by PCR from the gDNA of *Aspergillus sclerotiorum* LZDX-33-4 using the primer pairs and assembled with the NheI-linearized pFAL1 expression vector (containing *gpdA* promoter, *trpC* terminator and uracil marker). The above fragments assembly experiments were performed using ClonExpress ® MultiS One Step Cloning Kit (Vazyme Biotech Co., Ltd) for propagation in *E. coli* DH5α. All plasmids were verified by sanger sequencings performed by Genewiz (Azanta Inc., China).

*Aspergillus nidulans* LO8030 was used as the recipient host. The protoplast preparation and transformation were performed as described previously [1]. pHZQ1-2 containing the C77 genes g4415-4418 were transformed into *A. nidulans* to create expression strain for compounds **1**-**4**. pHZQ4 containing the C128 genes g7211-7212 were transformed into *A. nidulans* to create expression strain for compound **5**.

**LC-MS**

LC-MS analysis was performed on an Agilent HPLC 1260 series system equipped with a Bruker microTOF QIII mass spectrometer by using an Agilent Eclipse XDB C18 column (5 μm, 4.6 × 150 mm) or Waters UPLC-MS system. The parameters of the spectrometer were set as the following: electrospray ionization, capillary voltage with 4.5 kV, collision energy with 8.0 eV. Sodium formate was used in each run for mass calibration. The masses were scanned in the range of *m/z* 100 ‒ 1500 in ESI^+^ and ESI^-^ modes. Data were evaluated with the Compass Data Analysis 4.2 software (Bruker Daltonik, Bremen, Germany).

**GC-MS**

Analysis of crude extract: GC-MS analysis was carried out on an Agilent 7890A GC System equipped with a B5975C MSD by using an Agilent 19091S-433(30 m, 0.25 mm, 0.25 µm). A flow rate of 1.5 mL min-1 of helium was used as carrier gas and electronic impact was 70 eV. Oven temperature was programmed at 50 °C hold for 4 min, from 50 °C to 280 °C in 11.5 min, and 280 °C hold for 5 min.

Analysis of compound **4**: GC-MS analysis was also carried out on an Thermo Trace-1300 ISQ-Mass System equipped with a Thermo TraceGOLD TG-5MS column (30 m, 0.25 mm, 0.25 µm). A flow rate of 2.0 mL min-1 of helium was used as carrier gas and electronic impact was 70 eV. Oven temperature was programmed at 50 °C hold for 3 min, from 50 °C to 280°C in 11.5 min, and 280 °C hold for 3 min. Data were acquisited evaluated with the Xcalibur software (Thermo Scientific, Massachusetts, USA).

**UPLC-DAD analysis**

UPLC analysis was performed on an Agilent UPLC series 1200 (Agilent Technologies) equipped with an Agilent Eclipse XDB-C18 column (5 μm, 4.6 × 150 mm). A linear gradient 10 to 100% ACN in H_2_O in 7 min was used at a flowrate of 0.4 mL/min. The column was then washed with 100% ACN for 2 min and equilibrated with 10% ACN for 1 min. Detection was carried out with a photodiode array detector from 190 to 600 nm.

**Semi-preparative HPLC**

The semi-prep-HPLC was conducted with a YMC-packed C18 column (5 μm, 10 × 250 mm). The products were eluted with different solvent gradients of ACN in H_2_O, with or without CF_3_CO_2_H, at a flowrate of 2 mL/min.

**Large-scale fermentation, extraction and isolation of secondary metabolites**

To obtain compounds **1**-**4** from THZQ-1, the strain was cultivated in 20 bottles of rice media, each containing 60 g rice and 55 mL water at 28 °C for 14 days. For metabolites extraction after large-scale fermentation, mycelia on rice media were extracted with EtOAc three times. After extraction, the crude extract (6.4 g) was subjected to a silica gel column, eluting with a gradient of cyclohexane-ethyl acetate (from 20:1 to 1:1). The fractions containing the desired products were further dried and subjected to Sephadex LH-20 column using dichloromethane-methanol (1:1) to obtain compound **2**, followed by semi-preparative HPLC with a mobile phase of ACN-H2O (9:20) containing 0.01% trifluoroacetic acid (TFA) to yield compounds **1**, **3**, and **4**.

To obtain compounds **5** from THZQ-4, the strain was cultivated in 20 bottles of rice media at 28 °C for 14 days. For metabolites extraction after large-scale fermentation, mycelia on rice media were extracted with EtOAc three times. After extraction, the crude extract (15.0 g) was subjected to a silica gel column, eluting with a gradient of cyclohexane-dichloromethane (from 10:1 to 1:1) to obtain compound **5** and 95% MeOH/H_2_O to obtain compound **5**.

**NMR and ECD**

The NMR data were recorded using a Bruker FTNMR spectrometer at 400/500/600 MHz. 2D spectra (COSY, HSQC and HMBC) were recorded using standard parameters. Samples were dissolved in deuterated as solvents indicated in the respective.

Optical rotations were measured on an Autopol III automatic polarimeter. UV spectra were detected on an Alltech UVIS-200 detector. The electronic circular dichroism (ECD) spectra were measured using a JASCO J-810/J-815 spectropolarimeter.

**Physiochemical properties of compounds 1-4**

Compound **1**: [α]22 D −20.9 (*c* 0.23, MeOH); ^13^C NMR and ^1^H NMR data, see Table S7-S8; HRESIMS *m/z* 219.1742 [M+H-2H_2_O]^+^ (calcd for C_15_H_23_O, 219.1749).

Compound **2**: [α]22 D −14.7 (*c* 0.1, MeOH); ^13^C NMR and ^1^H NMR data, see Table S7-S8; HRESIMS *m/z* 221.1913 [M+H-H_2_O]^+^ (calcd for C_15_H_25_O, 221.1905).

Compound **3**: [α]22 D −10.0 (*c* 0.04, MeOH); ^13^C NMR and ^1^H NMR data, see Table S7-S8; HRESIMS *m/z* 201.1651 [M+H-3H_2_O]^+^ (calcd for C_15_H_21_, 201.1643).

Compound **4**: [α]22 D −4.7 (*c* 0.1, MeOH); ^13^C NMR and ^1^H NMR data, see Table S7-S8; HRESIMS *m/z* 219.1742 [M+H-2H_2_O]^+^ (calcd for C_15_H_23_O, 219.1749)

Compound **5**: [α]25 D −80.0 (*c* 0.1, CHCl_3_); ^13^C NMR and ^1^H NMR NMR data, see Table S9-S10; EIMS *m/z* 340.45 [M-H_2_O]^+^.

**References**

1.	Chiang, Y.M., et al., *Development of Genetic Dereplication Strains in Aspergillus nidulans Results in the Discovery of Aspercryptin.* Angew Chem Int Ed Engl, 2016. **55**(5): p. 1662-5.

**Supplemental Tables**

**Supplemental Table S1. Core enzyme families summarized from FunBGCs-defined core enzyme subtypes**

| **Core enzyme family** | **FunBGCs-defined core enzyme subtype(s) (Frequency)** | **Frequency** |
| --- | --- | --- |
| PKS | NR-PKS (155), HR-PKS (146), T3PKS (12),  PR-PKS (8), PKS-like (1) | 291 |
| NRPS | NRPS (147), NRPS-like (31) | 174 |
| TC | TC (Class1) (55), TC (Pyr4) (54), TC (Tri5) (22),  TC (SHC/OSC) (10), TC (PbcA) (8), TC (AstC) (4),  TC (UbiA) (3), TC (AsR6) (2), TC (ABA3) (2) | 156 |
| PT | PT (DMATS) (70), PT (UbiA) (45), PT (PaxC) (15) | 118 |
| PKS-NRPS | PKS-NRPS (72), NRPS-PKS (14) | 86 |
| TS-chimeric | chimeric TS (33) | 33 |
| PPPS | PPPS (33) | 33 |

**Supplemental Table S2. BGC entries of FunBGCs and MIBiG included in this work**

See the attached Excel file.

**Supplemental Table S3. Genome entries of EnsemblFungi and JGI MycoCosm included in this work**

See the attached Excel file.

**Supplemental Table S4. BGC-genome linkages determined by zero-mismatched DNA sequence alignment**

See the attached Excel file.

**Supplemental Table S5. Nodes (domains) of attention weight network**

See the attached Excel file.

**Supplemental Table S6. Edges (domain-domain pairs) of attention weight network**

See the attached Excel file.

**Supplemental Table S7. ^13^C NMR data of compounds 1-4 (1 in DMSO-*d*6, 150 MHz; 2-4 in CD_3_OD, 100 MHz)**

| **No.** | **1** | **2** | **3** | **4** |
| --- | --- | --- | --- | --- |
|  | ***δ*_C_** | ***δ*_C_** | ***δ*_C_** | ***δ*_C_** |
| 1 | 38.0 | 37.5 | 37.4 | 38.8 |
| 2 | 18.4 | 26.7 | 26.7 | 17.8 |
| 3 | 41.3 | 78.3 | 78.1 | 35.2 |
| 4 | 32.9 | 38.3 | 38.3 | 37.1 |
| 5 | 53.9 | 49.6 | 49.3 | 42.4 |
| 6 | 133.4 | 23.7 | 22.8 | 22.8 |
| 7 | 127.9 | 122.3 | 124.9 | 124.9 |
| 8 | 72.3 | 133.5 | 137.0 | 137.1 |
| 9 | 55.1 | 56.8 | 54.3 | 54.3 |
| 10 | 36.5 | 35.4 | 35.2 | 35.0 |
| 11 | 57.7 | 59.7 | 59.8 | 59.9 |
| 12 | 68.7 | 20.7 | 65.4 | 65.6 |
| 13 | 15.7 | 13.4 | 13.5 | 16.8 |
| 14 | 33.0 | 27.2 | 27.2 | 70.3 |
| 15 | 22.2 | 14.4 | 14.4 | 14.1 |

**Supplemental Table S8. ^1^H NMR data of compounds 1-4 (1 in DMSO-*d*6, 600 MHz; 2-4 in CD_3_OD, 400 MHz)**

| **No.** | **1** | **2** | **3** | **4** |
| --- | --- | --- | --- | --- |
|  | ***δ*_H_** | ***δ*_H_** | ***δ*_H_** | ***δ*_H_** |
| 1 | 1.76, 1H, m | 2.07, 1H, dt (13.2, 3.6) | 2.12, 1H, m | 2.11, 1H, m |
|  | 1.09, 1H, m | 1.30, 1H, ddd (14.4, 13.2, 4.4) | 1.33, 1H, ddd (14.2, 13.2, 5.1) | 1.16, 1H, ddd (14.3, 13.2, 3.9) |
| 2 | 1.59, 1H, m | 1.68, 1H, m | 1.68, 1H, m | 1.69, 1H, m |
|  | 1.41, 1H, m | 1.63, 1H, m | 1.63, 1H, m | 1.55, 1H, m |
| 3 | 1.39, 1H, m | 3.20, 1H, dd (11.2, 4.6) | 3.22, 1H, dd (10.5, 5.3) | 1.66, 1H, dt (13.6, 3.2) |
|  | 1.13, 1H, m |  |  | 1.51, 1H, ddd (14.2, 13.6, 3.5) |
| 4 |  |  |  |  |
| 5 | 1.52,1H, dd (3.0, 2.2) | 1.20, 1H, dd (10.6, 6.3) | 1.26, 1H, dd (11.4, 5.0) | 1.61, 1H, dd (12.5, 4.6) |
| 6 | 5.59, 1H, dd (10.2,3.1) | 2.00, 2H, m | 2.06, 2H, m | 2.05, 1H, m |
|  |  |  |  | 1.94, 1H, m |
| 7 | 5.69, 1H, dd (10.2,2.0) | 5.47, 1H, brs | 5.80, 1H, brd (4.6) | 5.78, 1H, brs |
| 8 |  |  |  |  |
| 9 | 1.36, 1H, t (4.7) | 1.82, 1H, m | 2.04, 1H, m | 2.03, 1H, m |
| 10 |  |  |  |  |
| 11 | 3.73, 1H, dt (11.5, 4.7) | 3.81, 1H, dd (11.2, 3.1) | 3.86, 1H, dd (11.1, 2.1) | 3.87, 1H, dd (11.0, 2.5) |
|  | 3.56, 1H, dt (11.5, 4.7) | 3.58, 1H, dd (11.2, 6.6) | 3.64, 1H, dd (11.2, 7.2) | 3.64, 1H, dd (11.2, 7.5) |
| 12 | 3.26, 2H, d (6.0) | 1.77, 3H, brs | 4.27, 1H, d (12.5) | 4.27, 1H, d (12.6) |
|  |  |  | 3.98, 1H, d (12.5) | 3.97, 1H, d (12.6) |
| 13 | 0.87, 3H, s | 0.83, 3H, s | 0.82, 3H, s | 0.85, 3H, s |
| 14 | 0.88, 3H, s | 0.98, 3H, s | 0.87, 3H, s | 3.33, 1H, d (11.2) |
|  |  |  |  | 3.03, 1H, d (11.2) |
| 15 | 0.83, 3H, s | 0.86, 3H, s | 0.86, 3H, s | 0.86, 3H, s |

**Supplemental Table S9. Comparison of the ^13^C NMR data of compound 5 to sesterfisherol (in CDCl_3_, 150 MHz)**

| **No.** | **5** | **sesterfisherol** | **No.** | **5** | **sesterfisherol** |
| --- | --- | --- | --- | --- | --- |
|  | ***δ*_C_** | ***δ*_C_** |  | ***δ*_C_** | ***δ*_C_** |
| 1 |  | - | 14 | 43.9 | 44.1 |
| 2 | 44.7 | 44.9 | 15 | 41.4 | 41.6 |
| 3 |  | - | 16 | 41.1 | 41.2 |
| 4 |  | - | 17 | 28.4 | 28.5 |
| 5 | 26.4 | 26.4 | 18 | 47.1 | 47.3 |
| 6 | 43.8 | 43.9 | 19 | 31.5 | 31.6 |
| 7 | 38.4 | 38.5 | 20 |  | - |
| 8 | 29.0 | 29.2 | 21 | 15.1 | 15.2 |
| 9 | 32.0 | 32.0 | 22 | 17.8 | 17.9 |
| 10 | 143.0 | 143.1 | 23 | 18.5 | 18.7 |
| 11 | 133.0 | 133.0 | 24 | 22.6 | 22.7 |
| 12 | 78.5 | 78.5 | 25 | 24.0 | 24.1 |
| 13 | 39.2 | 39.3 |  |  |  |

**Supplemental Table S10. Comparison of the ^1^H NMR data of compound 5 to sesterfisherol (in CDCl_3_, 600 MHz)**

| **No.** | **5** | **sesterfisherol** | **No.** | **5** | **sesterfisherol** |
| --- | --- | --- | --- | --- | --- |
|  | ***δ*_H_** | ***δ*_H_** |  | ***δ*_H_** | ***δ*_H_** |
| 1 |  |  | 14 | 1.89, 1H, m | 1.89, 1H, m |
| 2 | 1.39, 1H, m | 1.40, 1H, m | 15 | - | - |
| 3 |  |  | 16 | 1.44, 1H, m | 1.44, 1H, m |
| 4 |  |  |  | 1.14, 1H, m | 1.13, 1H, m |
| 5 | 1.42, 1H, m | 1.41, 1H, m | 17 | 1.82, 1H, m | 1.83, 1H, m |
|  | 1.02, 1H, m | 1.00, 1H, m |  | 1.54, 1H, m | 1.54, 1H, m |
| 6 | 3.66, 1H, brs | 3.66, 1H, brs | 18 | 1.66, 1H, m | 1.65, 1H, m |
| 7 | 2.12, 1H, m | 2.12, 1H, m | 19 | 1.59, 1H, m | 1.58, 1H, m |
| 8 | 1.68, 1H, m | 1.68, 1H, m | 20 | 0.87, 3H, d (7.1) | 0.87, 3H, d (7.2) |
|  | 1.29, 1H, m | 1.27, 1H, m | 21 | 0.94, 3H, d (6.7) | 0.94, 3H, d (6.7) |
| 9 | 2.32, 1H, m | 2.33, 1H, m | 22 | 1.70, 3H, s | 1.70, 3H, s |
|  | 2.17, 1H, m | 2.16, 1H, m | 23 | 0.80, 3H, s | 0.80, 3H, s |
| 10 | - | - | 24 | 0.83, 3H, d (6.5) | 0.83, 3H, d (6.0) |
| 11 | - | - | 25 | 0.90, 3H, d (6.2) | 0.90, 3H, d (6.0) |
| 12 | - | - |  |  |  |
| 13 | 2.14, 1H, m | 2.13, 1H, m |  |  |  |
|  | 1.57, 1H, m | 1.57, 1H, m |  |  |  |

**Supplemental Table S11. Optimized hyperparameters of GECCO and TOUCAN in BGC detection benchmark tests**

| **Trained with**  **type-2 negative gene**  **cluster samples?** | **Algorithm** | **Dataset partitioning scheme** | **Subtask** | **Optimized hyperparameter combination** |
| --- | --- | --- | --- | --- |
| No | GECCO | In-distribution | - | c1: 0, c2: 1 |
|  |  | Cross-SM-class | Testing cluster: 1 | c1: 0.25, c2: 1 |
|  |  |  | Testing cluster: 2 | c1: 0.125, c2: 2 |
|  |  |  | Testing cluster: 3 | c1: 0, c2: 0.5 |
|  |  |  | Testing cluster: 4 | c1: 0, c2: 1 |
|  |  |  | Testing cluster: 5 | c1: 0.125, c2: 0 |
|  |  |  | Testing cluster: 6 | c1: 0, c2: 0.125 |
|  |  | Cross-dataset | - | c1: 0.25, c2: 0.125 |
|  | TOUCAN | In-distribution | - | k-mer: 7 |
|  |  | Cross-SM-class | Testing cluster: 1 | k-mer: 6 |
|  |  |  | Testing cluster: 2 | k-mer: 6 |
|  |  |  | Testing cluster: 3 | k-mer: 6 |
|  |  |  | Testing cluster: 4 | k-mer: 6 |
|  |  |  | Testing cluster: 5 | k-mer: 6 |
|  |  |  | Testing cluster: 6 | k-mer: 6 |
|  |  | Cross-dataset | - | k-mer: 6 |
| Yes | GECCO | In-distribution | - | c1: 0.5, c2: 0 |
|  |  | Cross-SM-class | Testing cluster: 1 | c1: 0.125, c2: 0 |
|  |  |  | Testing cluster: 2 | c1: 1, c2: 0 |
|  |  |  | Testing cluster: 3 | c1: 0.25, c2: 0 |
|  |  |  | Testing cluster: 4 | c1: 0.5, c2: 0 |
|  |  |  | Testing cluster: 5 | c1: 0.5, c2: 0 |
|  |  |  | Testing cluster: 6 | c1: 0, c2: 0.125 |
|  |  | Cross-dataset | - | c1: 0.25, c2: 0 |
|  | TOUCAN | In-distribution | - | k-mer: 6 |
|  |  | Cross-SM-class | Testing cluster: 1 | k-mer: 4 |
|  |  |  | Testing cluster: 2 | k-mer: 6 |
|  |  |  | Testing cluster: 3 | k-mer: 4 |
|  |  |  | Testing cluster: 4 | k-mer: 4 |
|  |  |  | Testing cluster: 5 | k-mer: 4 |
|  |  |  | Testing cluster: 6 | k-mer: 4 |
|  |  | Cross-dataset | - | k-mer: 4 |

**Supplemental Table S12. Primers used in this study**

| **Primers** | **Sequence (5’-3’)** |
| --- | --- |
| C77-gpdA+3MFS-F | CCAAGAACCTTTAATCCCGGAATATCACTCTTCAGCA |
| C77-3MFS+TrpC-R | CTATTAAATCAGCTAGCCAGTATTCATACCGGGCCAT |
| C77-gpdA+DRI-F | CCAAGAACCTTTAATCCTCTCTGCACTGTCATCCC |
| C77-5P450+TrpC-R | CTATTAAATCAGCTAGCCAGAGCAACGCATGAAAGC |
| C77-gpdA+5p450-F | CCAAGAACCTTTAATCGAATGACACGAGCAGTGATG |
| C128-pgdA+STS-F | CCAAGAACCTTTAATCGATGGAGAATGTTTGGCGATATTC |
| C128-5P450+TrpC-R | CTATTAAATCAGCTAGCGTGCAGTCTTGCCCACTAC |

**Supplemental Table S13. Plasmids constructed in this study**

| **Plasmids** | **Description** |
| --- | --- |
| pHZQ-1 | C77-STS-P450 in pFAL1; uracil, uridine (UU) |
| pHZQ-2 | C77-MFS in pFAL3; riboflavin (R) |
| pHZQ-3 | C77-P450 in pFAL4; pyridoxine (P) |
| pHZQ-4 | C128-STS-P450 in pFAL1; uracil, uridine (UU) |

**Supplemental Table S14. Heterologous expression strain constructed in this study**

| **Strains** | **Description** |
| --- | --- |
| THZQ-1 | *A. nidulans* harboring pHZQ-1; medium with riboflavin and pyridoxine |
| THZQ-2 | *A. nidulans* harboring pHZQ-2; medium with riboflavin and pyridoxine |
| THZQ-3 | *A. nidulans* harboring pHZQ-3; medium with riboflavin and pyridoxine |
| THZQ-4 | *A. nidulans* harboring pHZQ-4; medium with riboflavin and pyridoxine |

**Supplemental Figures**

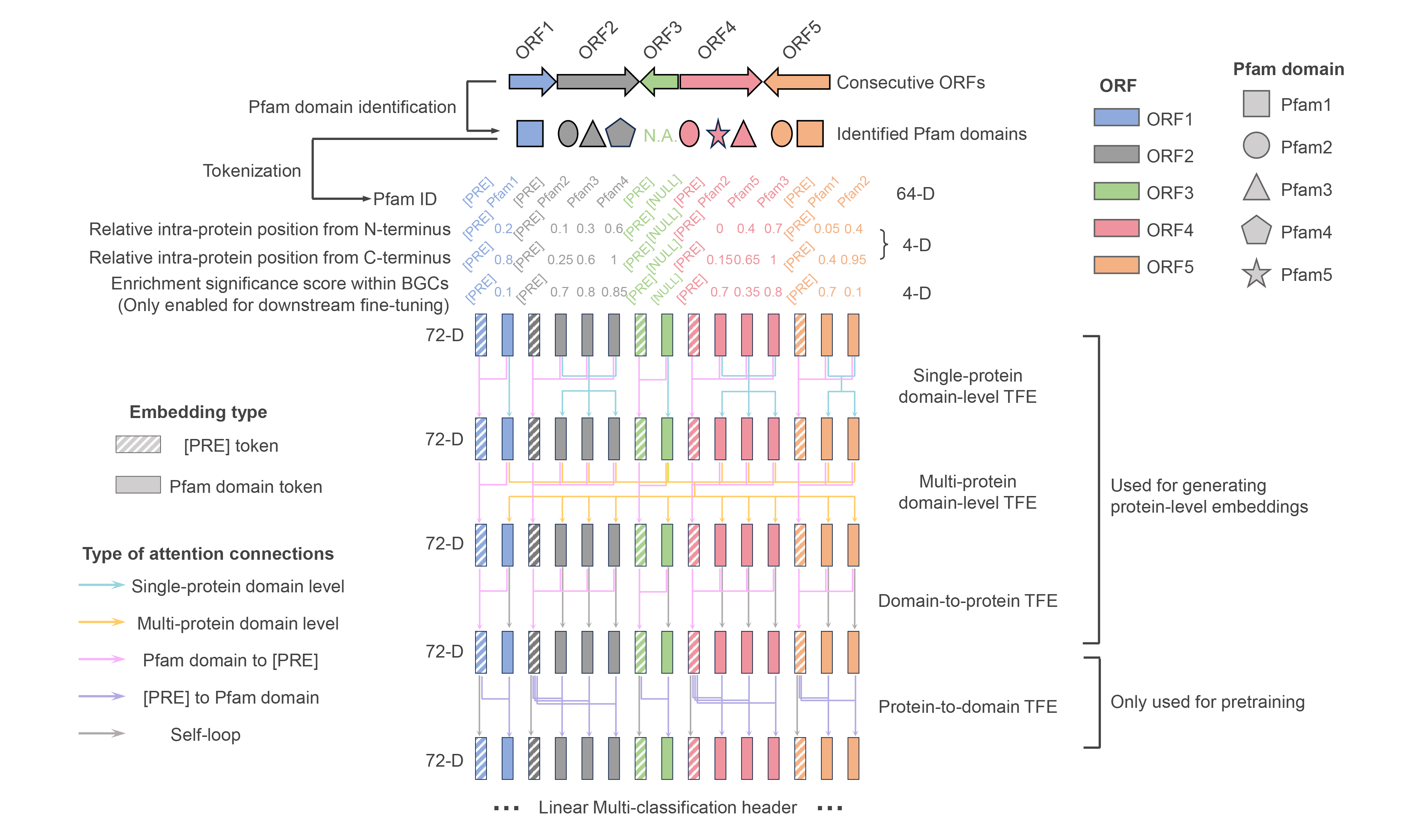

**Supplemental Figure S1. Overview of f-DLC model**

f-DLC receives ≤26 consecutive ORFs as inputs. First, f-DLC transforms the ORFs into linearized token sequences with a hierarchical domain-to-protein structure, which consists of Pfam domain tokens prefixed with [PRE] token for each ORF. Then the tokens are translated into 72-D embeddings consisting of three components: (1) 64-D domain-specific learnable embeddings; (2) 4-D embeddings encoding domains’ relative positioning information in protein; (3) 4-D embeddings encoding domains’ enrichment information in BGC (only enabled during downstream establishment of f-BGM). Next, the embeddings are passed through four sequential TFEs with different attention masks to achieve layer-wise information interaction. Specifically, the first and second TFE mainly focus on inter-domain information interaction within single protein and multiple proteins, respectively. Meanwhile the attention connections from domain tokens to corresponding prefix tokens are enabled for real-time domain-to-protein information aggregation. The third TFE only retains the domain-to-protein attention mechanism to fully summarize the upstream extracted features and thereby generate meaningful protein-level embeddings for downstream use. The fourth TFE reversely calculates on the attention connections from prefix tokens to domain tokens to satisfy domain-level BERT-style pretraining, where 15% of the input domain tokens are masked and the model is guided to recover these tokens by a downstream multi-classification header.

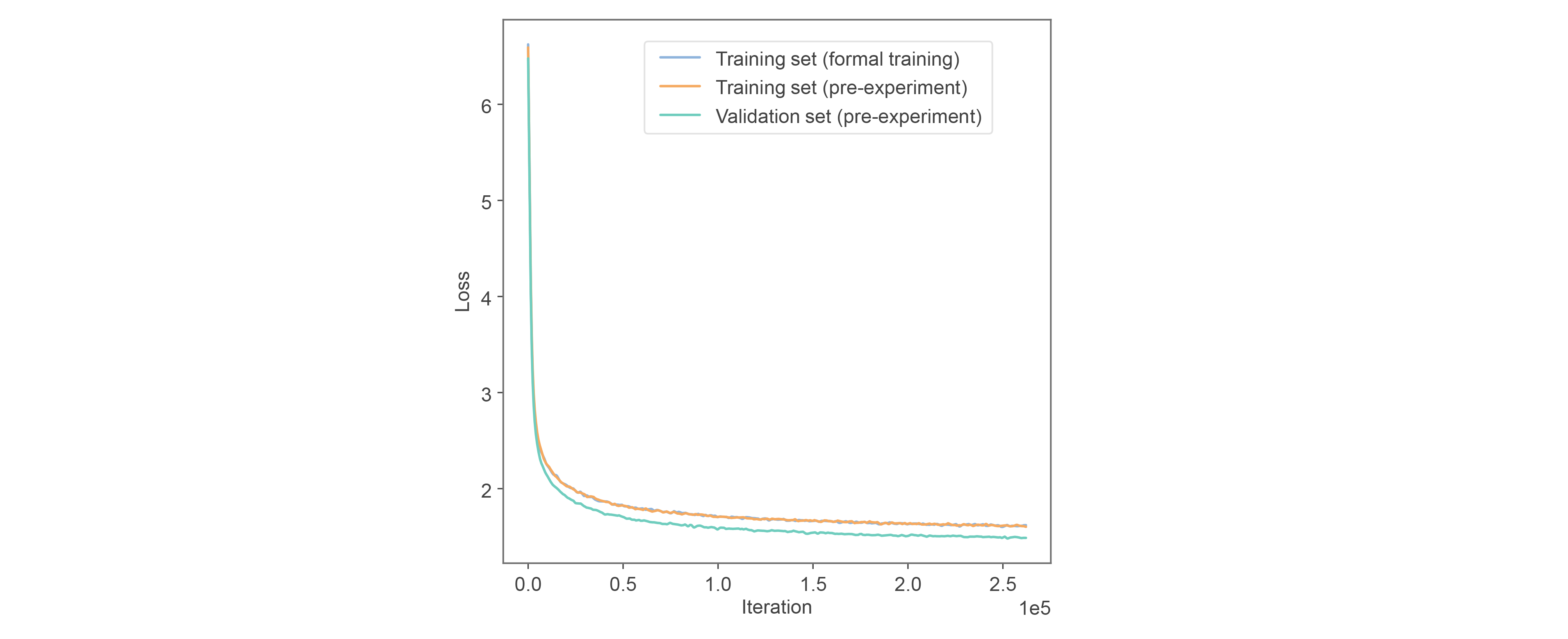

**Supplemental Figure S2. Loss convergence trend of f-DLC**

Loss convergence trend of f-DLC in pre-experiments and formal training. During the pre-experiments, the fungal genomes were randomly split into training and validation set at an 8:1 ratio. The convergence trend of validation loss proves that f-DLC is free of overfitting, supporting the formal training based on the whole genome dataset.

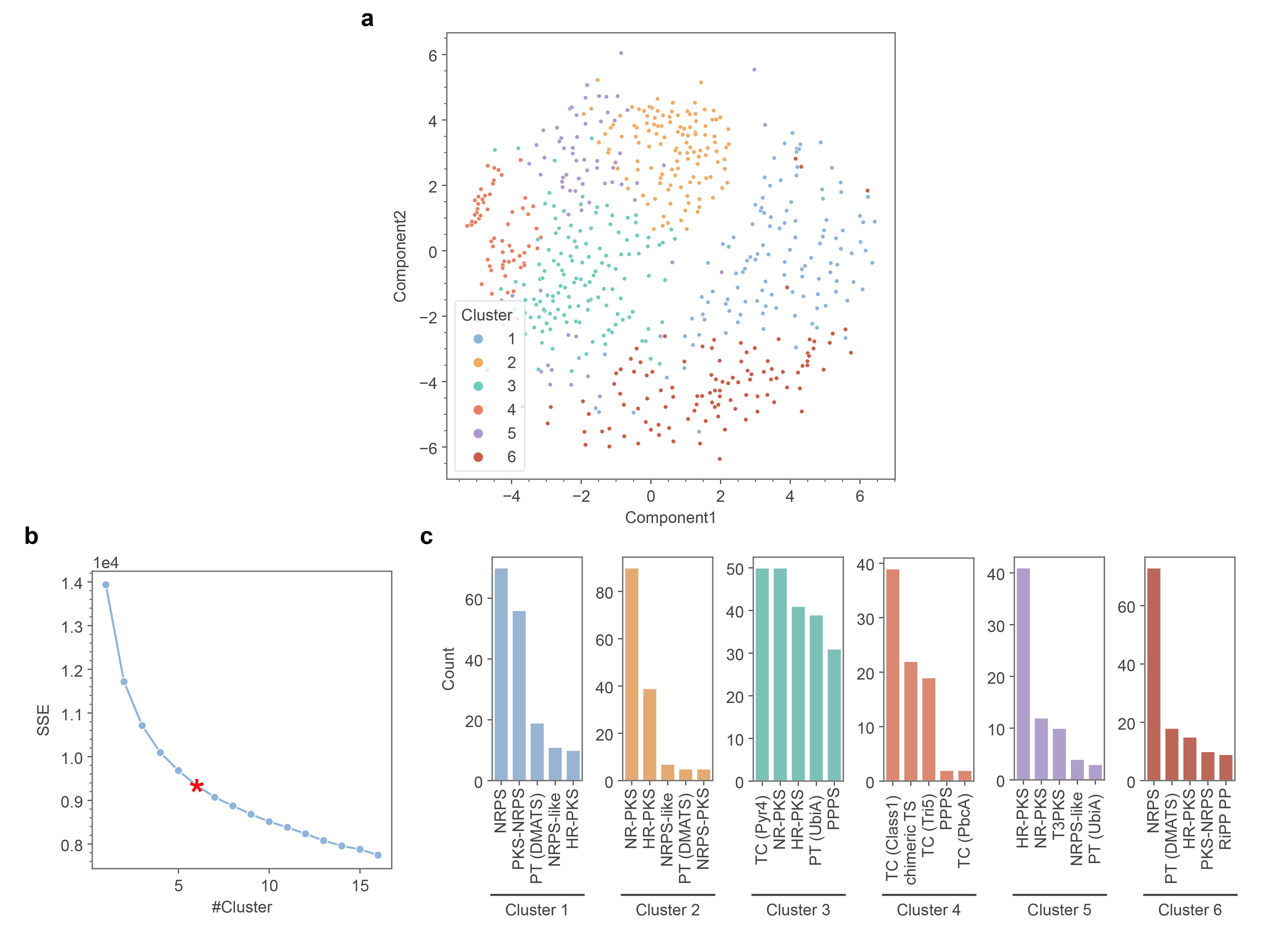

**Supplemental Figure S3. Cross-SM-class partitioning on FunBGCs dataset**

**a** Dimensionality reduction visualization of six BGC subsets defined by MACCSKeys-based K-means clustering on BGCs’ representative SM products. **b** Elbow method suggests 6 as a proper number for BGC clustering. **c** Highest-frequency core enzymes (following the FunBGCs’ definitions) of each BGC subset.

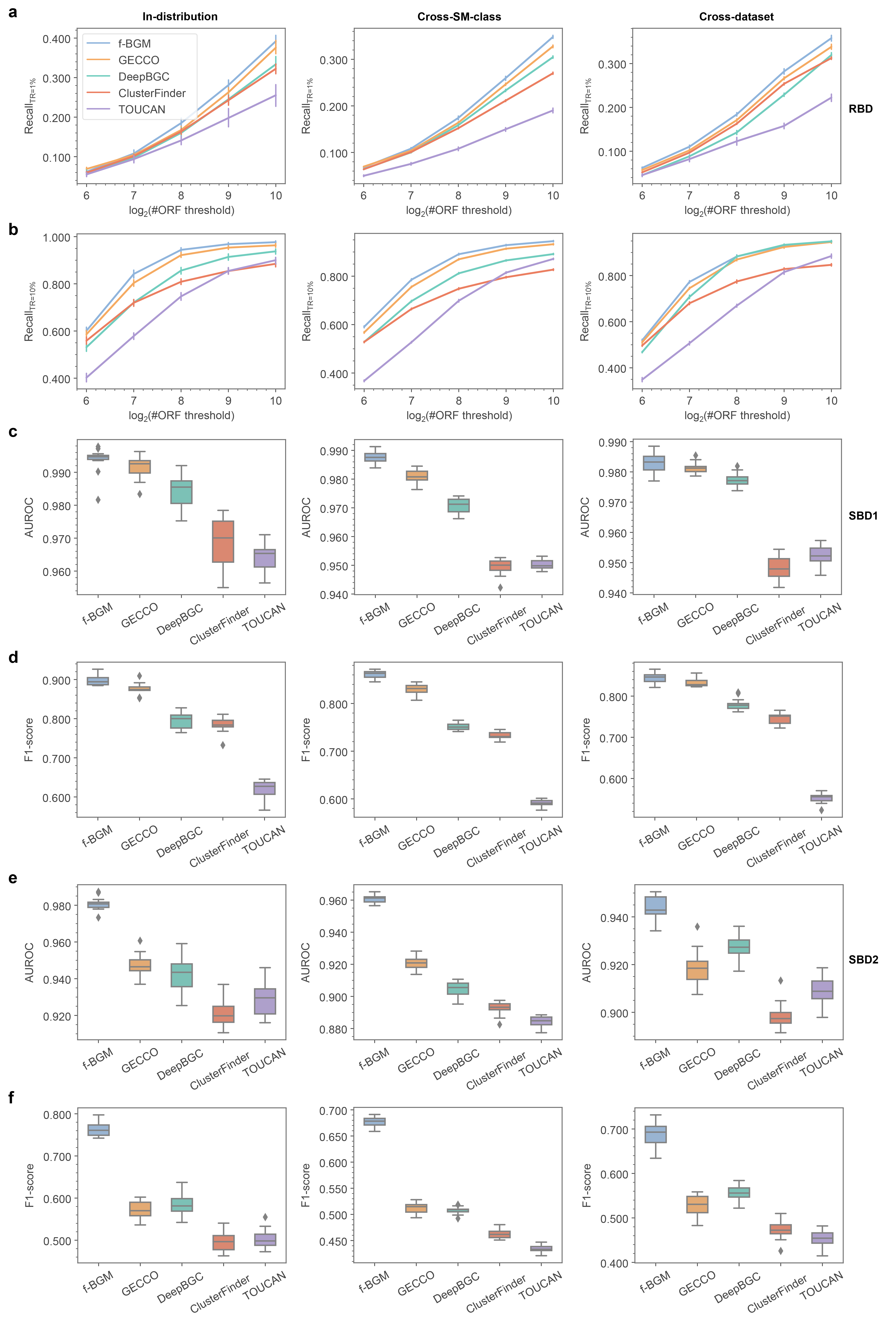

**Supplemental Figure S4. Performance of f-BGM in BGC detection benchmark tests (secondary metrics)**

**a, b** Performance comparison among f-BGM and four baseline algorithms with regard to (**a**) Recall_TR=1%_ and (**b**) Recall_TR=10%_ in real BGC detection tasks under in-distribution, cross-SM-class and cross-dataset schemes. **c, d** Performance comparison among f-BGM and four baseline algorithms with regard to (**c**) AUROC and (**d**) F1-score in standard simulated BGC detection tasks under in-distribution, cross-SM-class and cross-dataset schemes. **e, f** Performance comparison among f-BGM and four baseline algorithms with regard to (**e**) AUROC and (**f**) F1-score in challenging simulated BGC detection tasks under in-distribution, cross-SM-class and cross-dataset schemes. RBD: real BGC detection; SBD1: standard simulated BGC detection; SBD2: challenging simulated BGC detection.

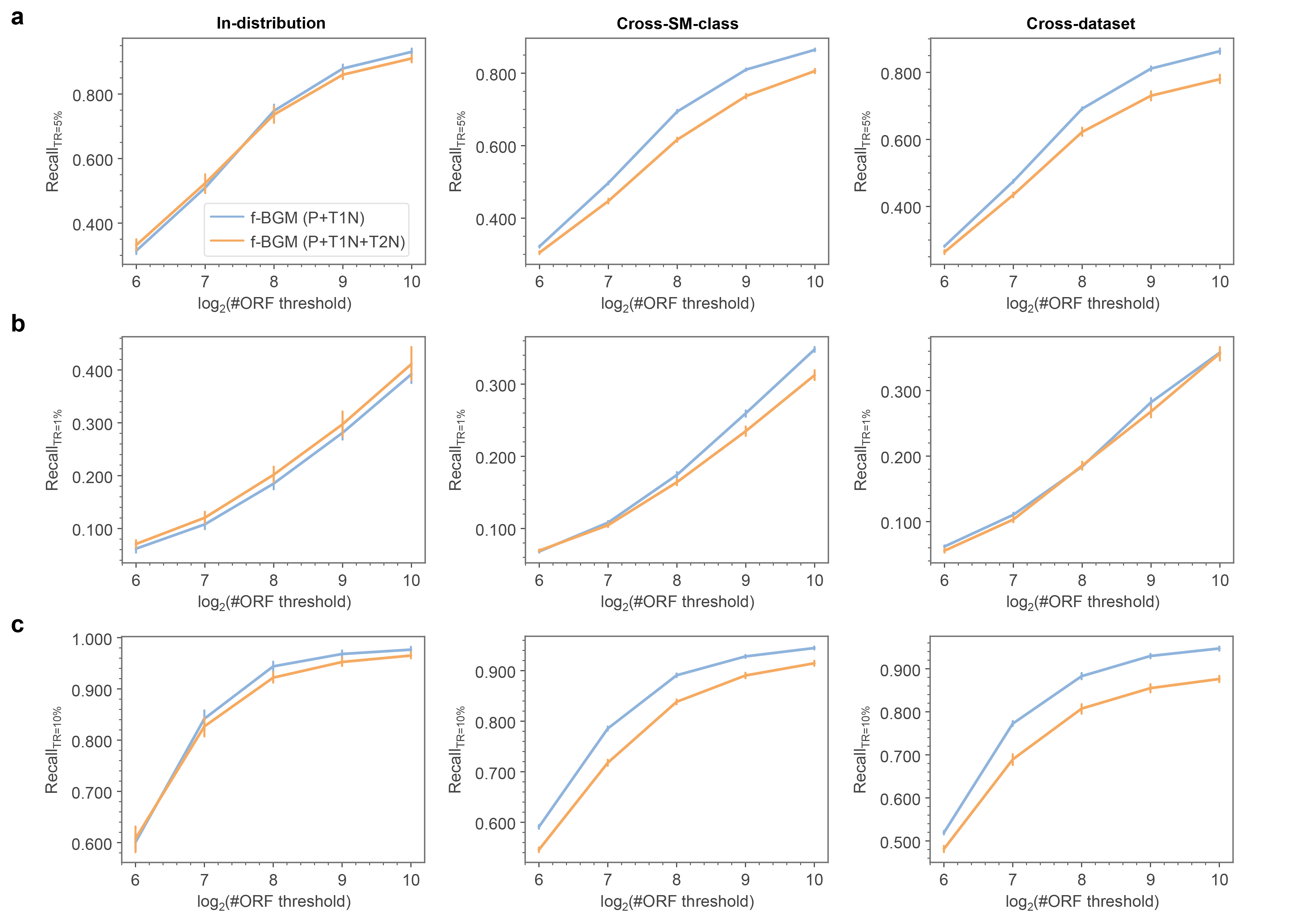

**Supplemental Figure S5. Performance of f-BGM in real BGC detection tasks with/without involvement of type-2 negative gene cluster samples**

**a-c** Self-performance comparison between f-BGMs trained with/without type-2 negative gene cluster samples with regard to (**a**) Recall_TR=5%_, (**b**) Recall_TR=1%_ and (**c**) Recall_TR=10%_ under in-distribution, cross-SM-class and cross-dataset schemes. f-BGM (P+T1N): f-BGM models trained from positive BGC samples + only type-1 negative gene cluster samples at a 1:16 ratio; f-BGM (P+T1N+T2N): f-BGM models trained from positive BGCs + type-1 negative samples + type-2 negative samples at a 1:12:4 ratio.

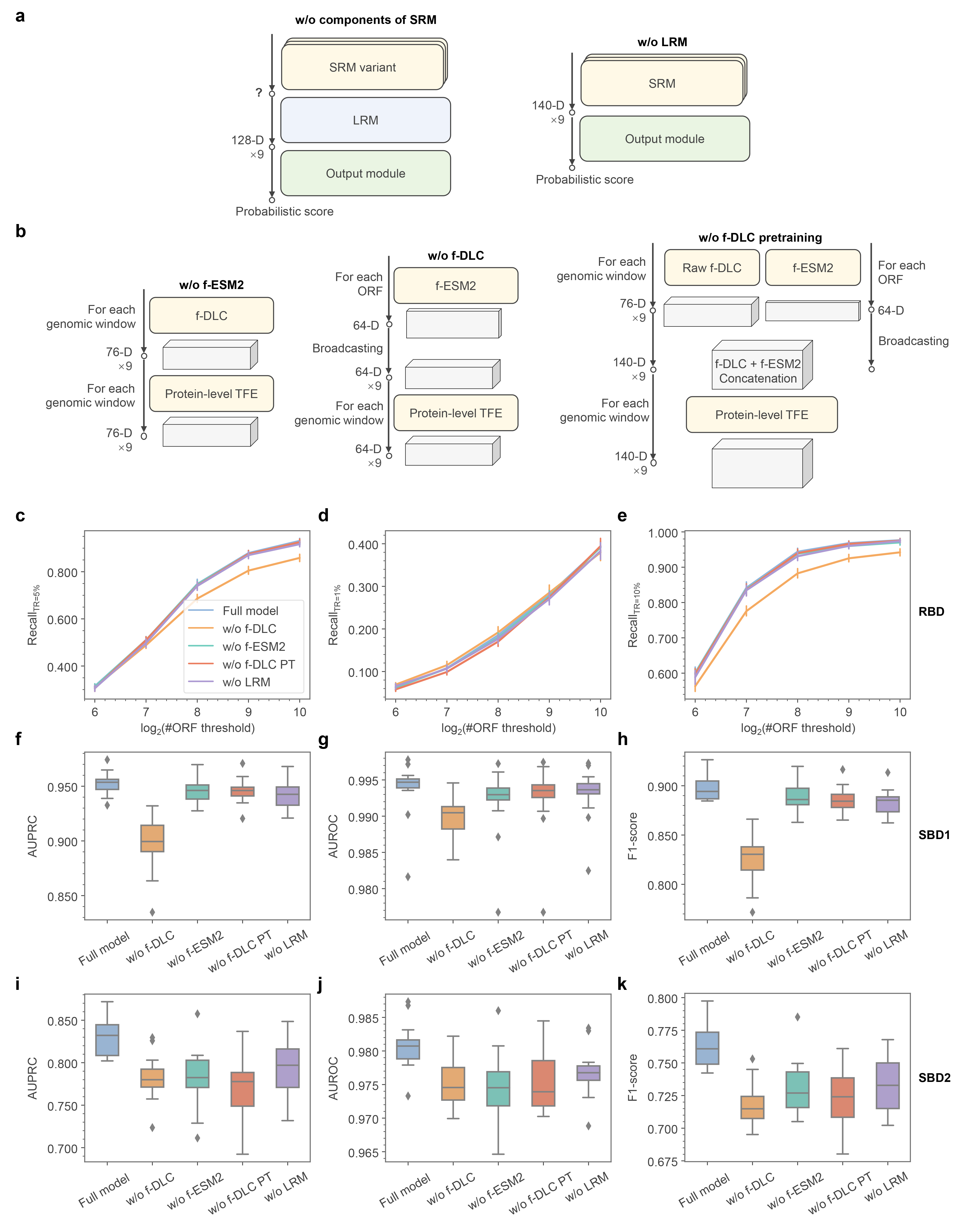

**Supplemental Figure S6. Ablation experiments of f-BGM in in-distribution BGC detection tasks**

**a** Illustration of f-BGM variants. **b** Detailed illustration of SRM variants in absence of individual components. **c-e** Performance comparison among the f-BGM full model and variants in real BGC detection tasks with regard to (**c**) Recall_TR=5%_, (**d**) Recall_TR=1%_ and (**e**) Recall_TR=10%_. **f-h** Performance comparison among the f-BGM full model and variants in standard simulated BGC detection tasks with regard to (**f**) AUPRC, (**g**) AUROC and (**h**) F1-score. **i-k** Performance comparison among the f-BGM full model and variants in challenging simulated BGC detection tasks with regard to (**i**) AUPRC, (**j**) AUROC and (**k**) F1-score. RBD: real BGC detection; SBD1: standard simulated BGC detection; SBD2: challenging simulated BGC detection.

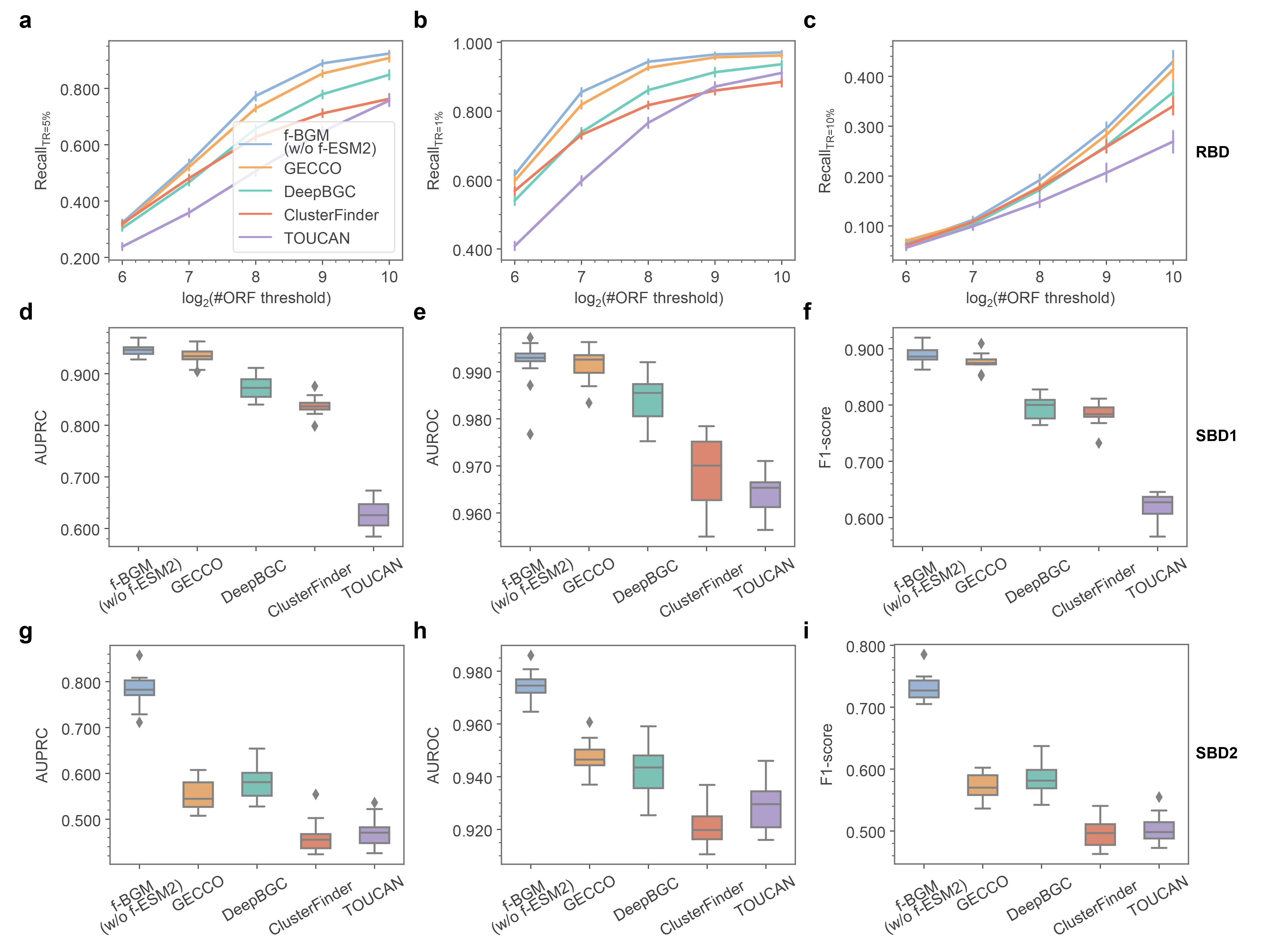

**Supplemental Figure S7. Performance of f-BGM variant with only Pfam domain features in in-distribution BGC detection benchmark tests**

**a-c** Performance comparison among the f-BGM variant without f-ESM2 and four baseline algorithms in real BGC detection tasks with regard to (**a**) Recall_TR=5%_, (**b**) Recall_TR=1%_ and (**c**) Recall_TR=10%_. **d-f** Performance comparison among the f-BGM variant without f-ESM2 and four baseline algorithms in standard simulated BGC detection tasks with regard to (**d**) AUPRC, (**e**) AUROC and (**f**) F1-score. **g-i** Performance comparison among the f-BGM variant without f-ESM2 and four baseline algorithms in challenging simulated BGC detection tasks with regard to (**g**) AUPRC, (**h**) AUROC and (**i**) F1-score. RBD: real BGC detection; SBD1: standard simulated BGC detection; SBD2: challenging simulated BGC detection.

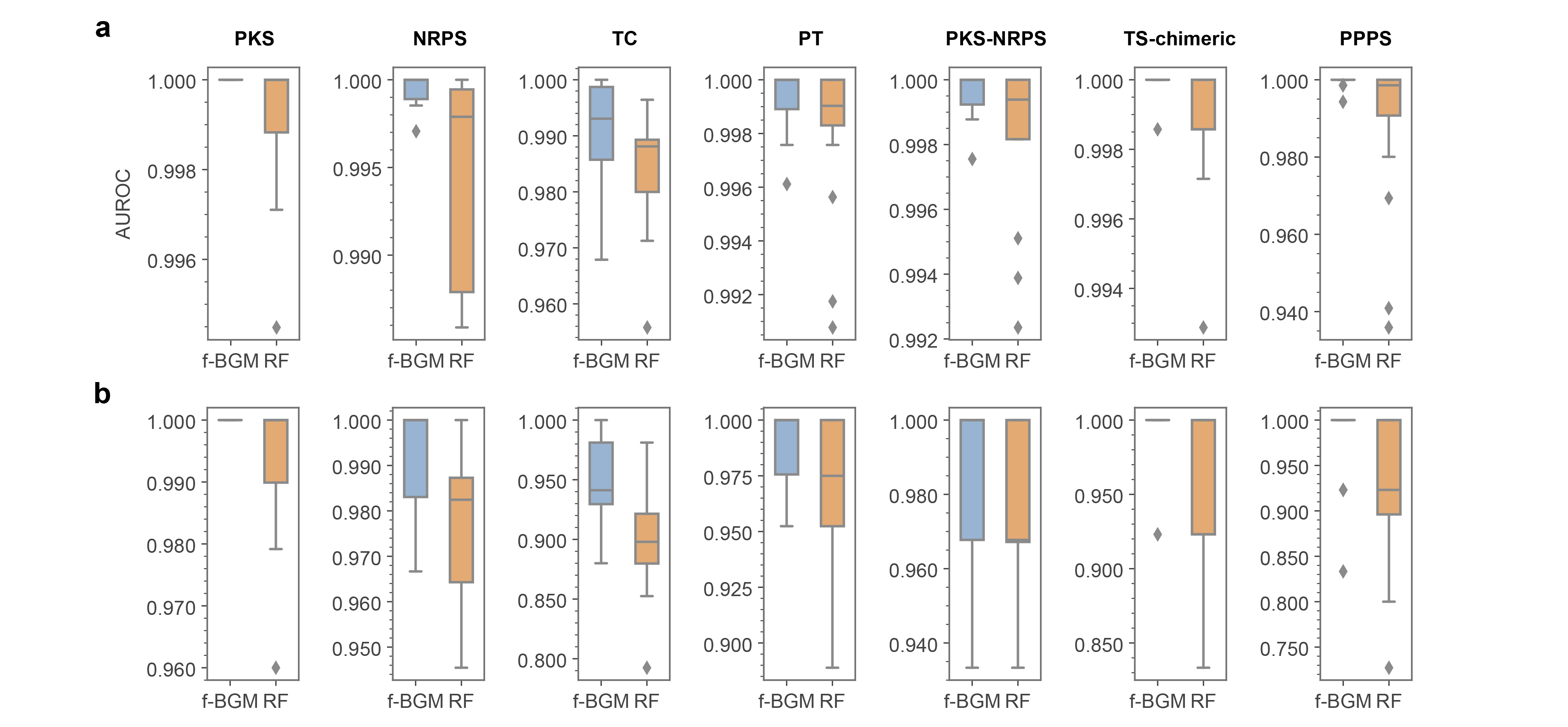

**Supplemental Figure S8. Performance of f-BGM in core enzyme identification benchmark tests (secondary metrics)**

**a, b** Performance comparison between f-BGM and RF in identifying seven major families of core enzymes with regard to (**a**) AUROC and (**b**) F1-score.

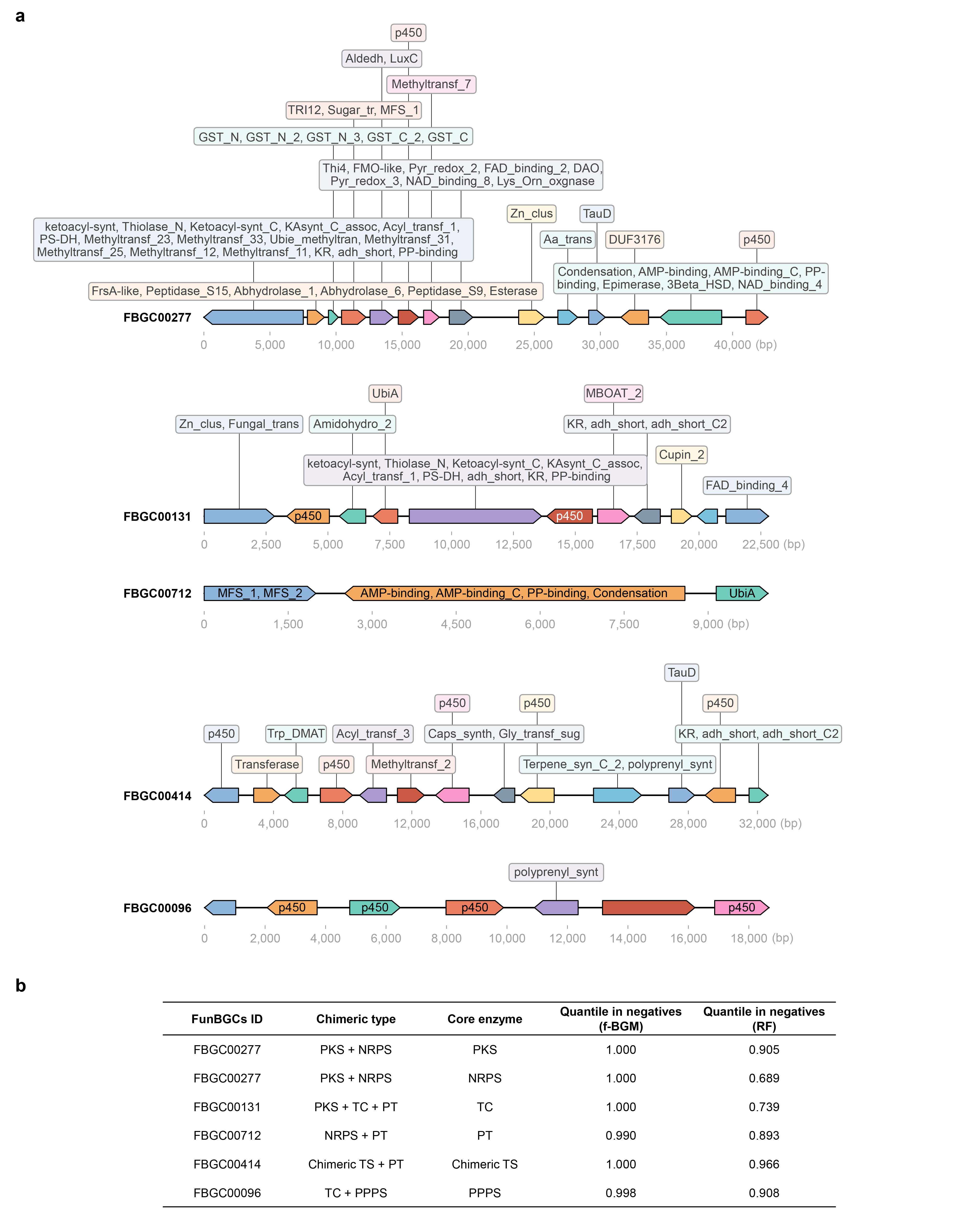

**Supplemental Figure S9. Representative chimeric BGCs prioritized by f-BGM over RF in core enzyme identification**

**a** Representative chimeric BGCs (harboring ≥2 core enzymes) prioritized by f-BGM over RF and their constitutions of Pfam domains. **b** Basic information of the BGCs shown in (**a**) including FunBGCs ID, chimeric type, the f-BGM-prioritized core enzyme and corresponding quantiles in negatives under f-BGM’s and RF’s scoring systems, respectively.

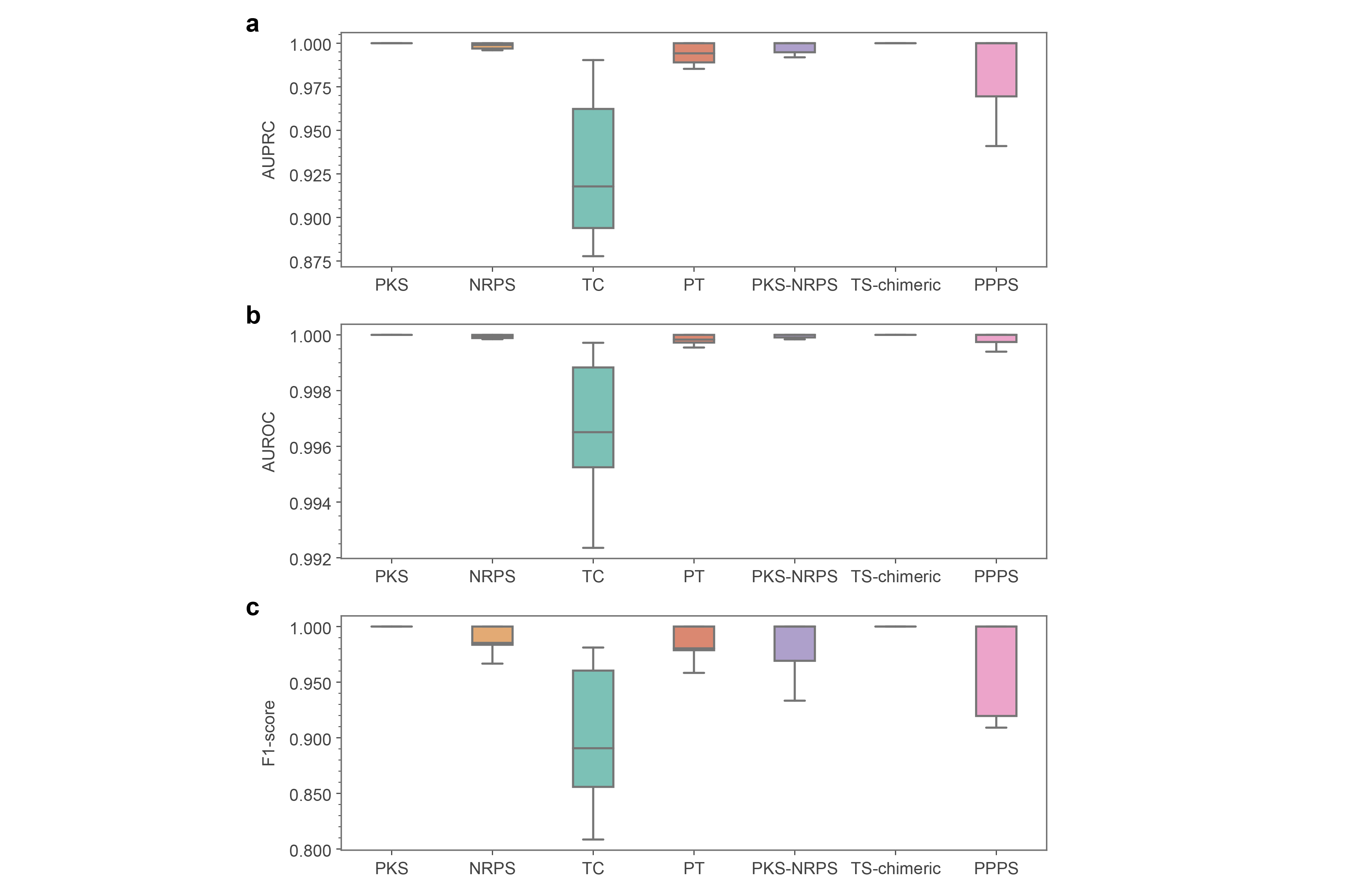

**Supplemental Figure S10. Performance of f-BGM in ORF-level core enzyme identification**

**a**-**c** Performance of f-BGM in high-resolution core enzyme identification at ORF level with regard to (**a**) AUPRC, (**b**) AUROC and (**c**) F1-score.

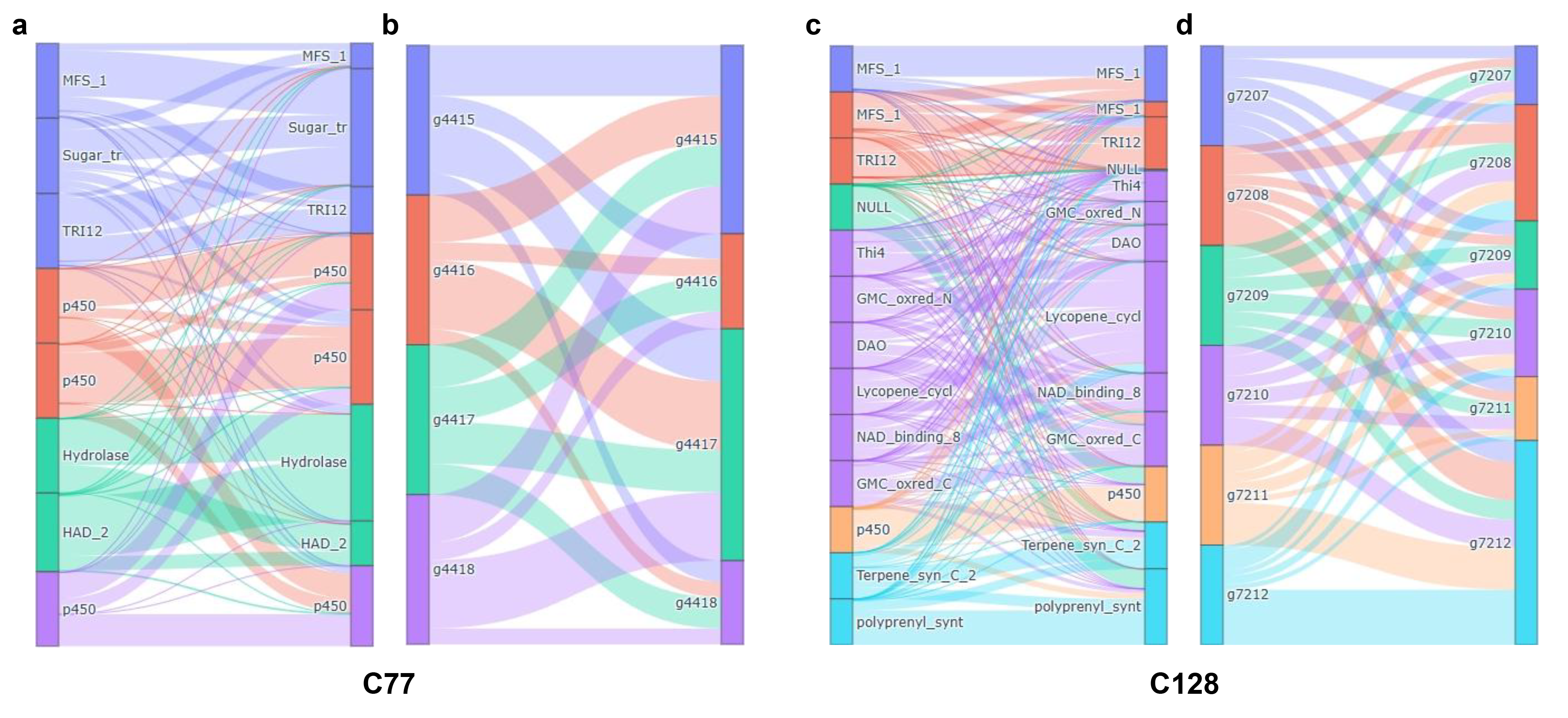

**Supplemental Figure S11. Attention weight visualization of LZDX-33-4-sourced gene clusters C77 and C128**

**a, c** Sankey plots illustrating inter-domain attention flows in (**a**) C77 and (**b**) C128, as revealed by multi-protein domain-level TFE. **b,** **d** Sankey plots illustrating inter-protein attention flows in (**b**) C77 and (**d**) C128, as revealed by protein-level TFE.

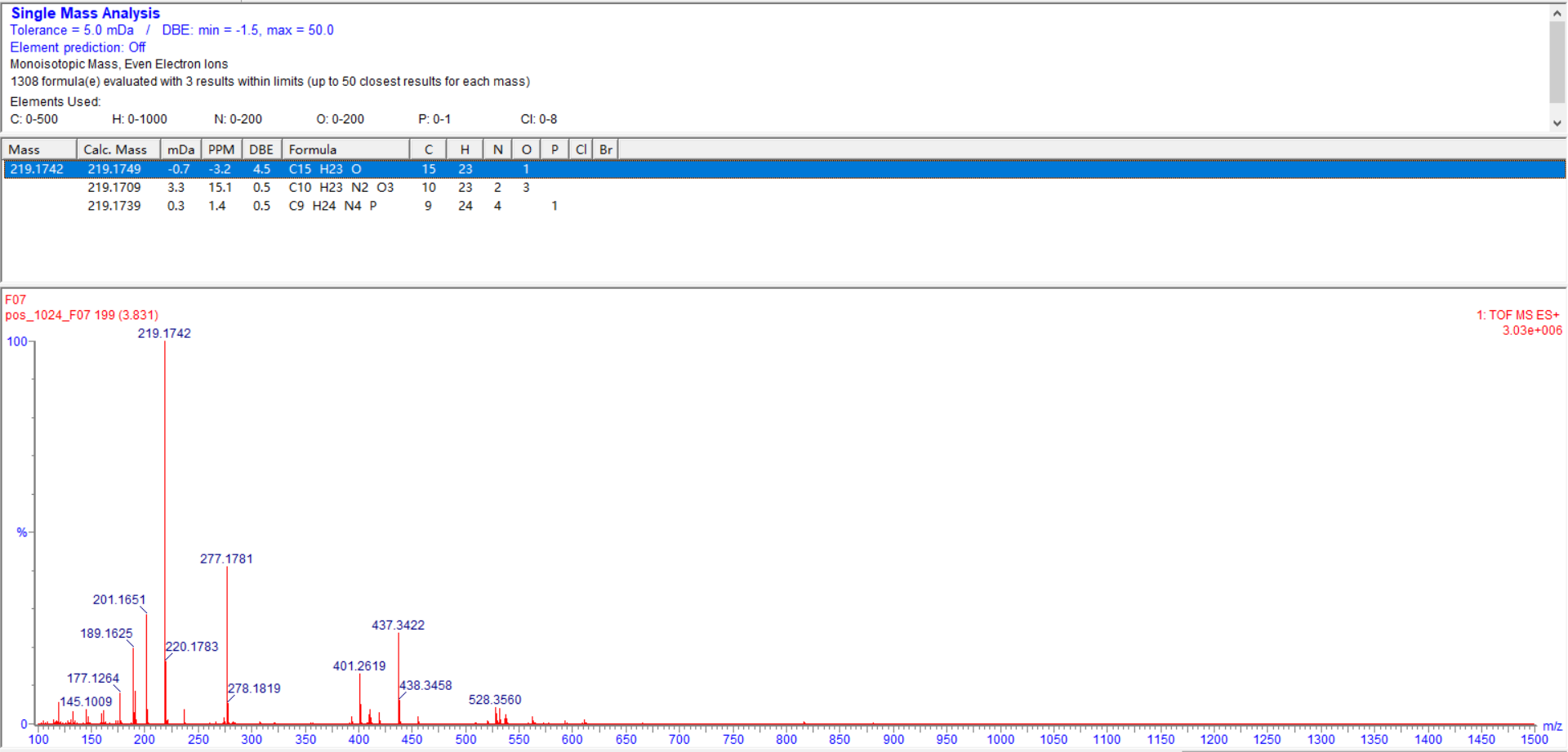

**Supplemental Figure S12. HRESIMS spectrum of compound 1**

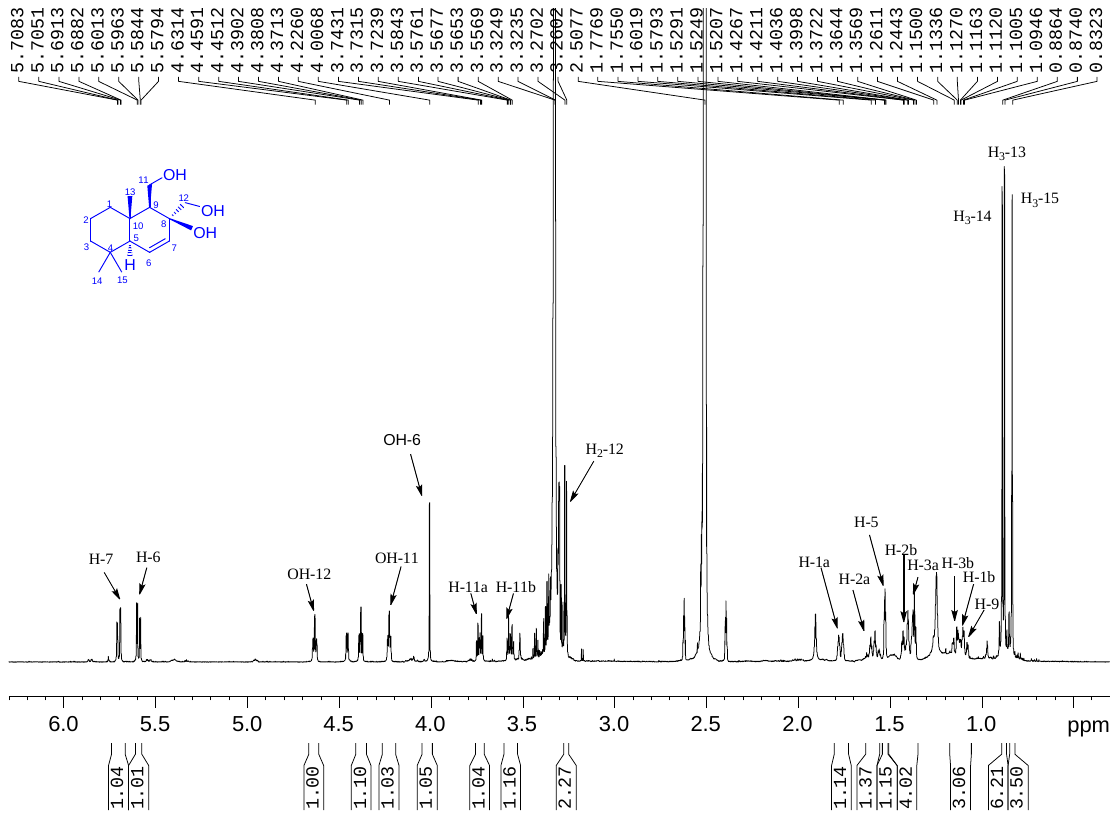

**Supplemental Figure S13. ^1^H NMR spectrum of compound 1 (600 MHz, DMSO-*d*_6_)**

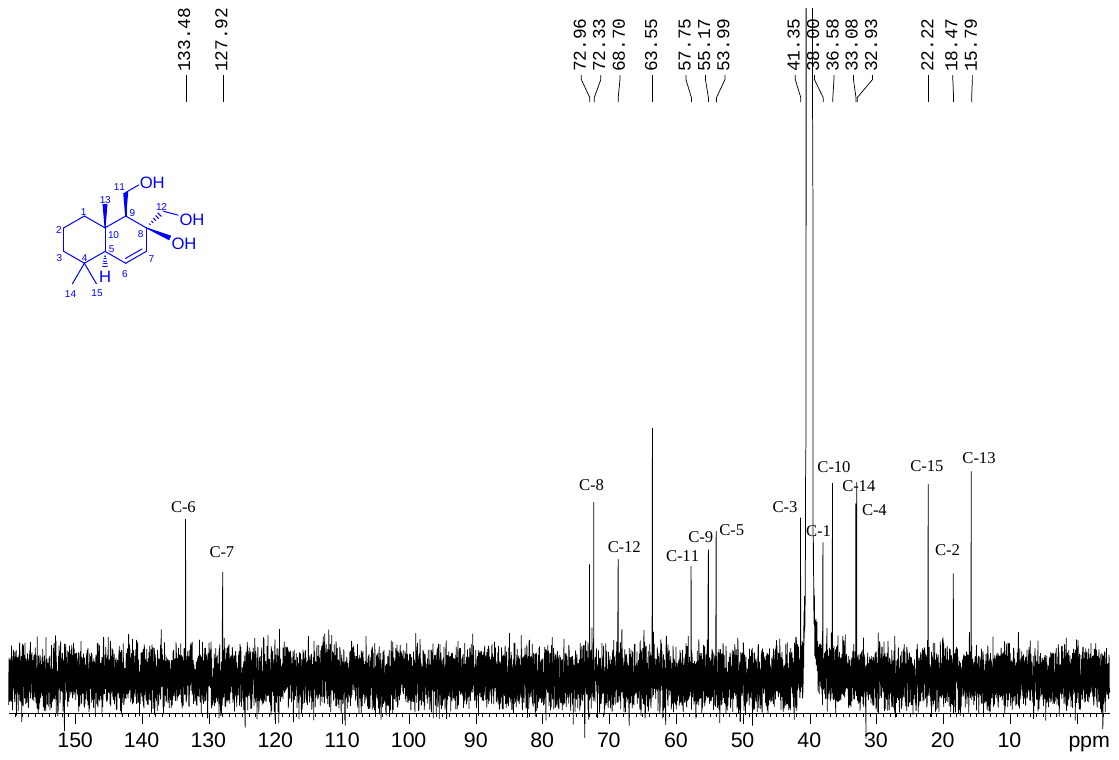

**Supplemental Figure S14. ^13^C NMR spectrum of compound 1 (150 MHz, DMSO-*d*_6_)**

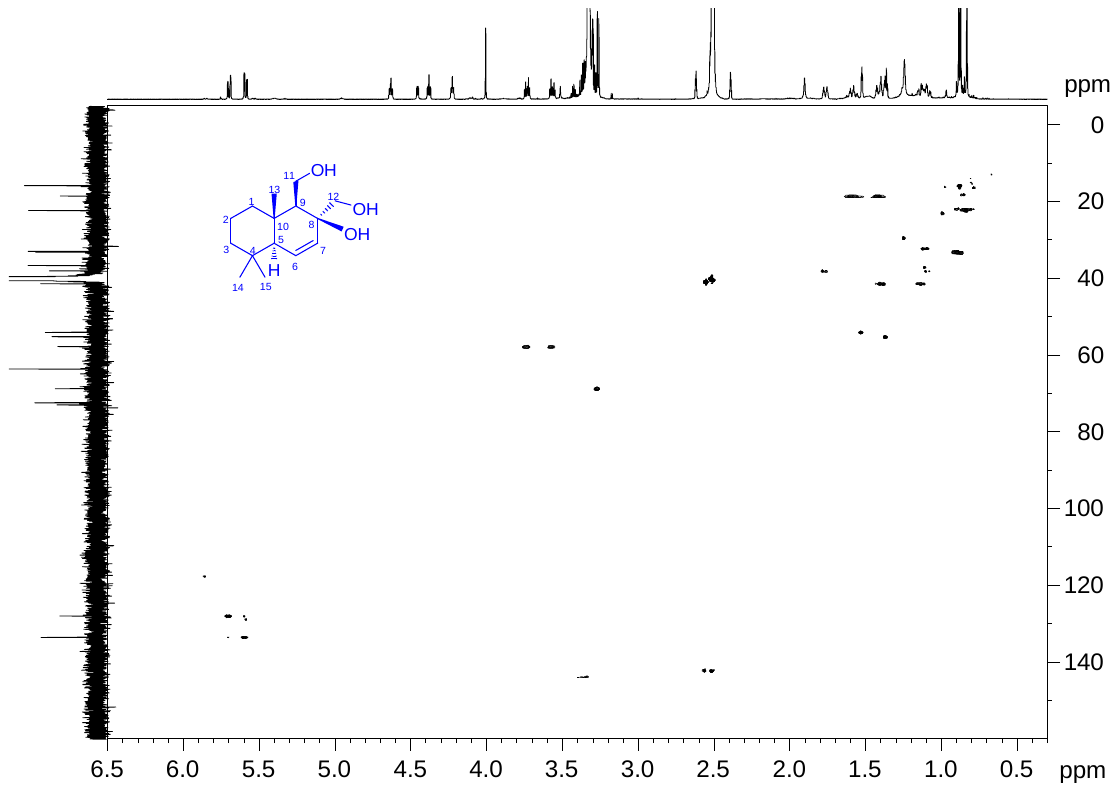

**Supplemental Figure S15. HSQC spectrum of compound 1 (DMSO-*d*_6_)**

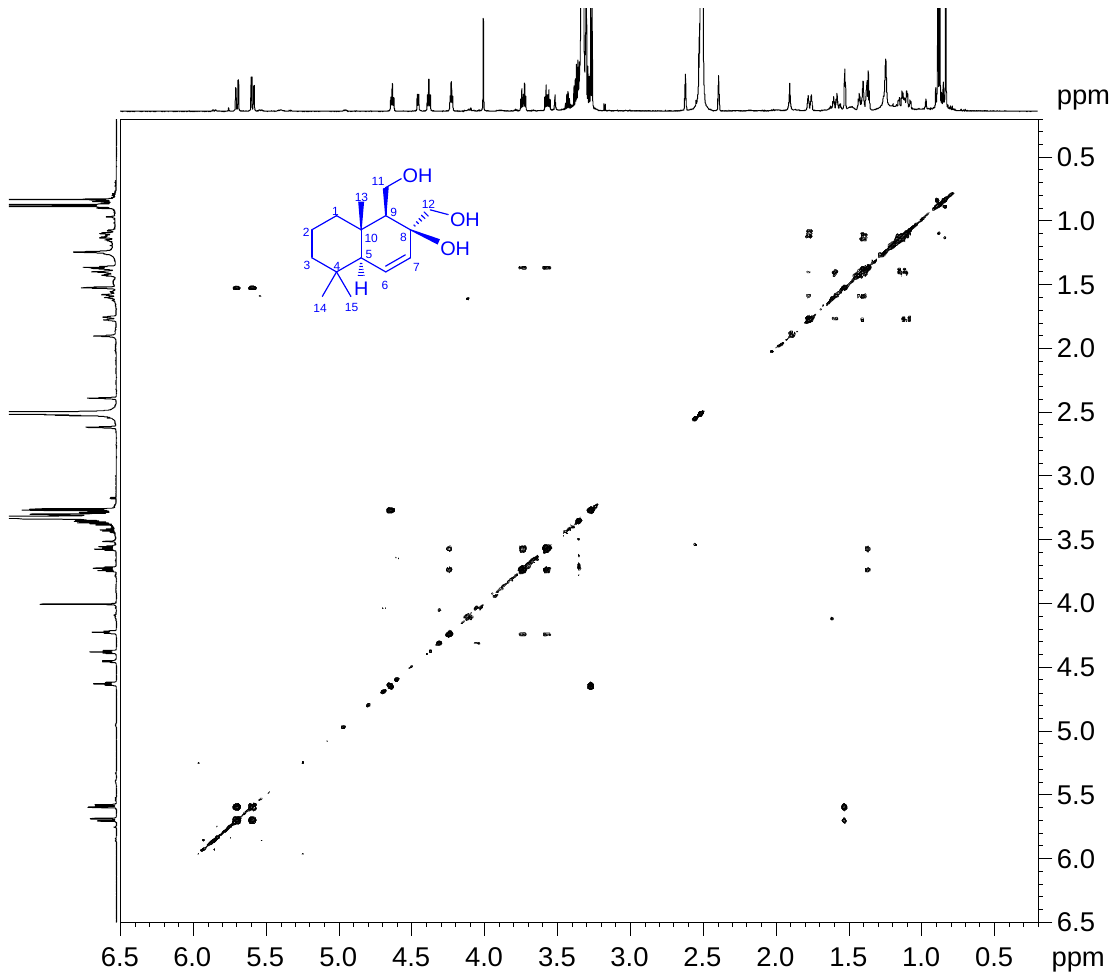

**Supplemental Figure S16. COSY spectrum of compound 1 (DMSO-*d*_6_)**

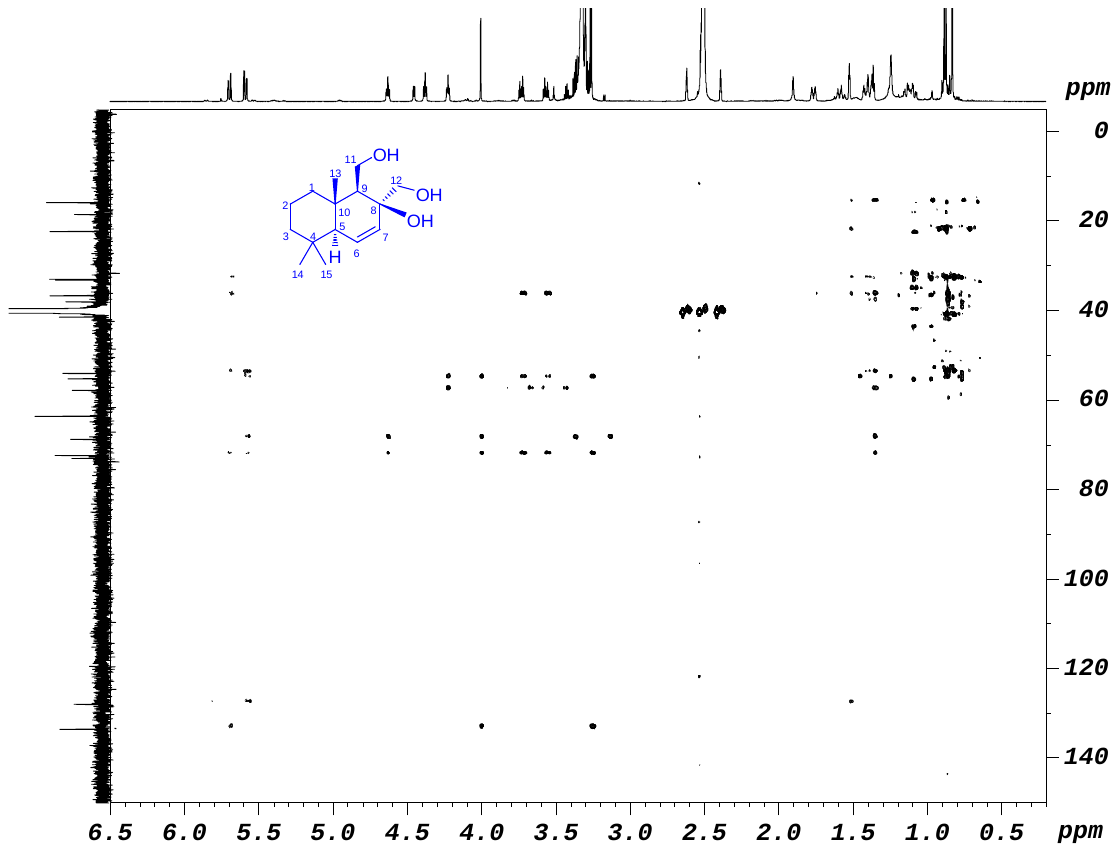

**Supplemental Figure S17. HMBC spectrum of compound 1 (DMSO-*d*_6_)**

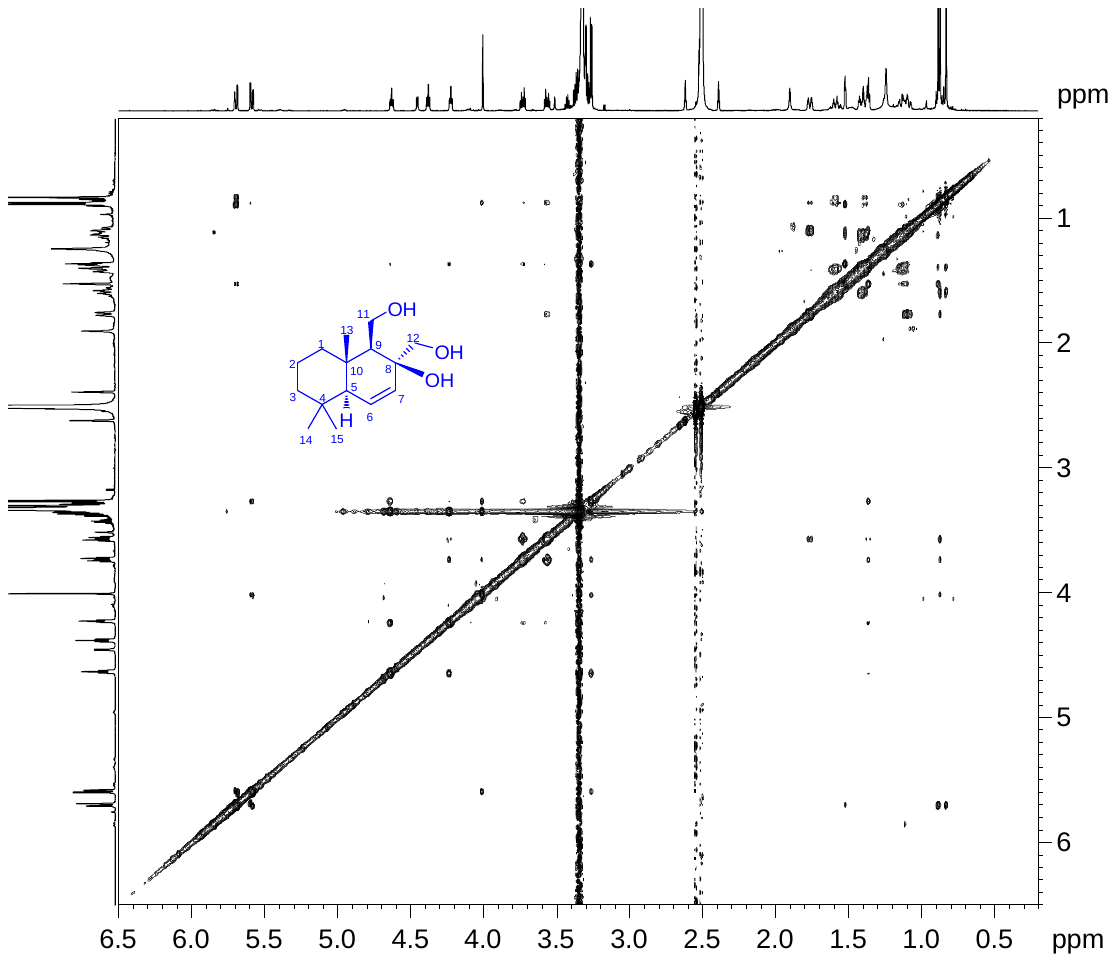

**Supplemental Figure S18. NOESY spectrum of compound 1 (DMSO-*d*_6_)**

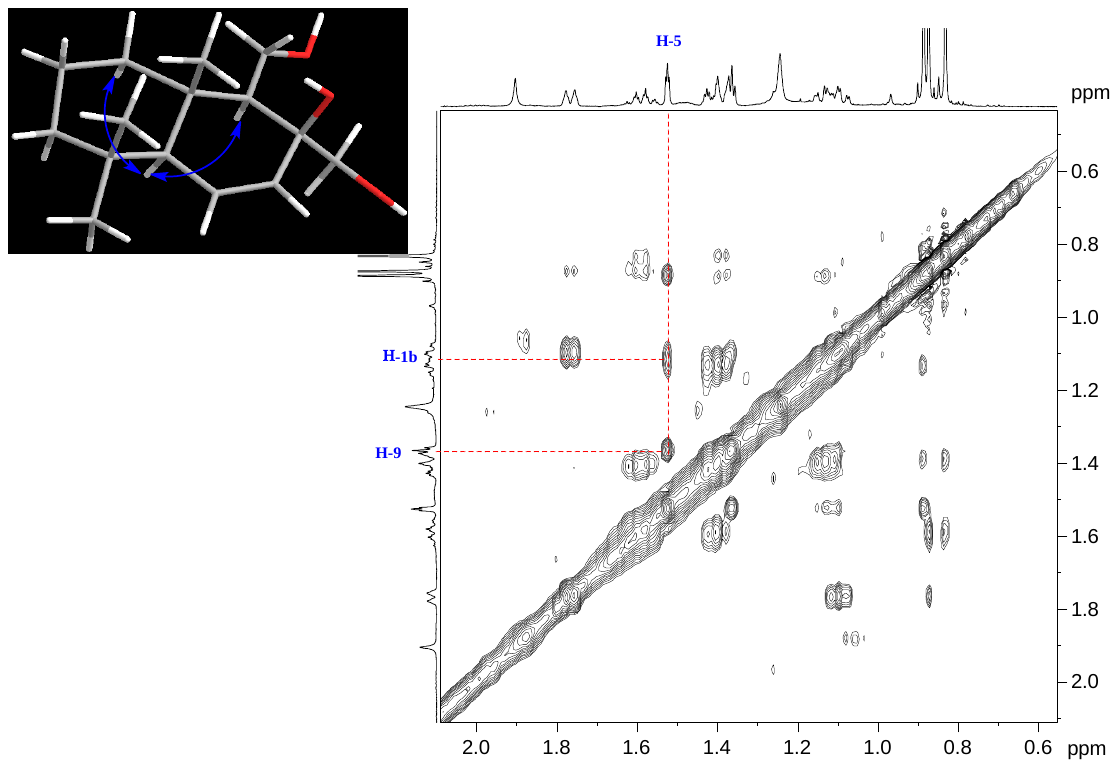

**Supplemental Figure S19. NOESY spectrum of compound 1 (DMSO-*d*_6_)**

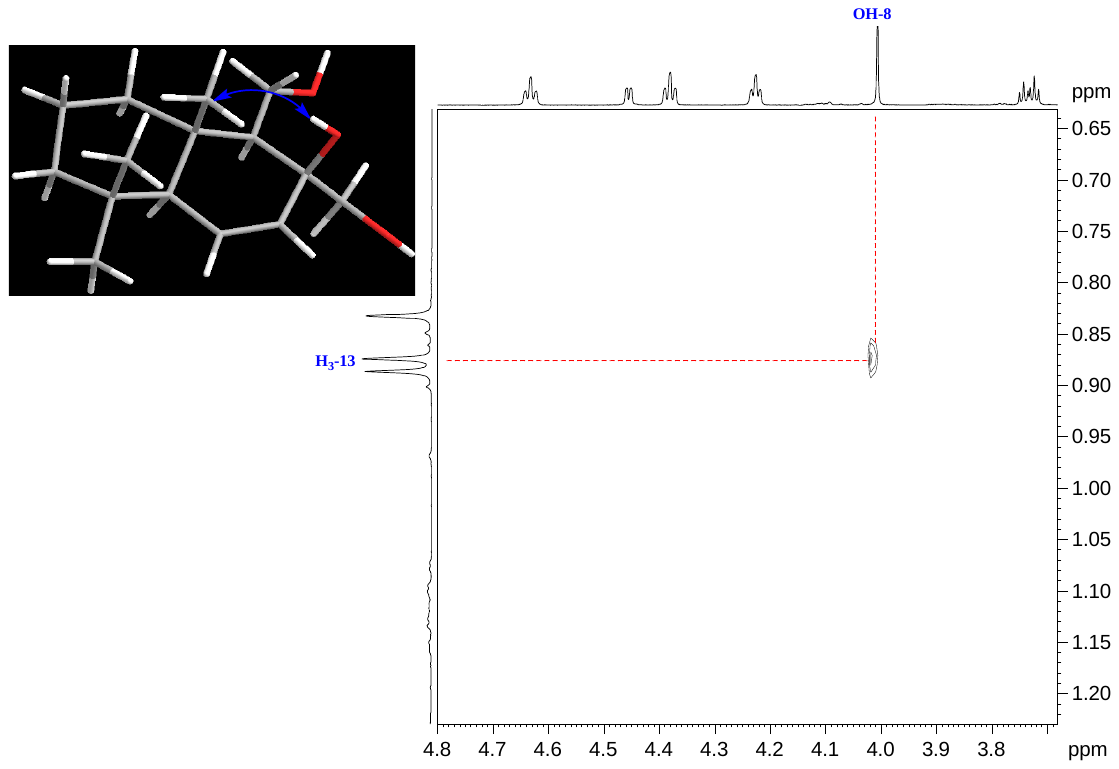

**Supplemental Figure S20. NOESY spectrum of compound 1 (DMSO-*d*_6_)**

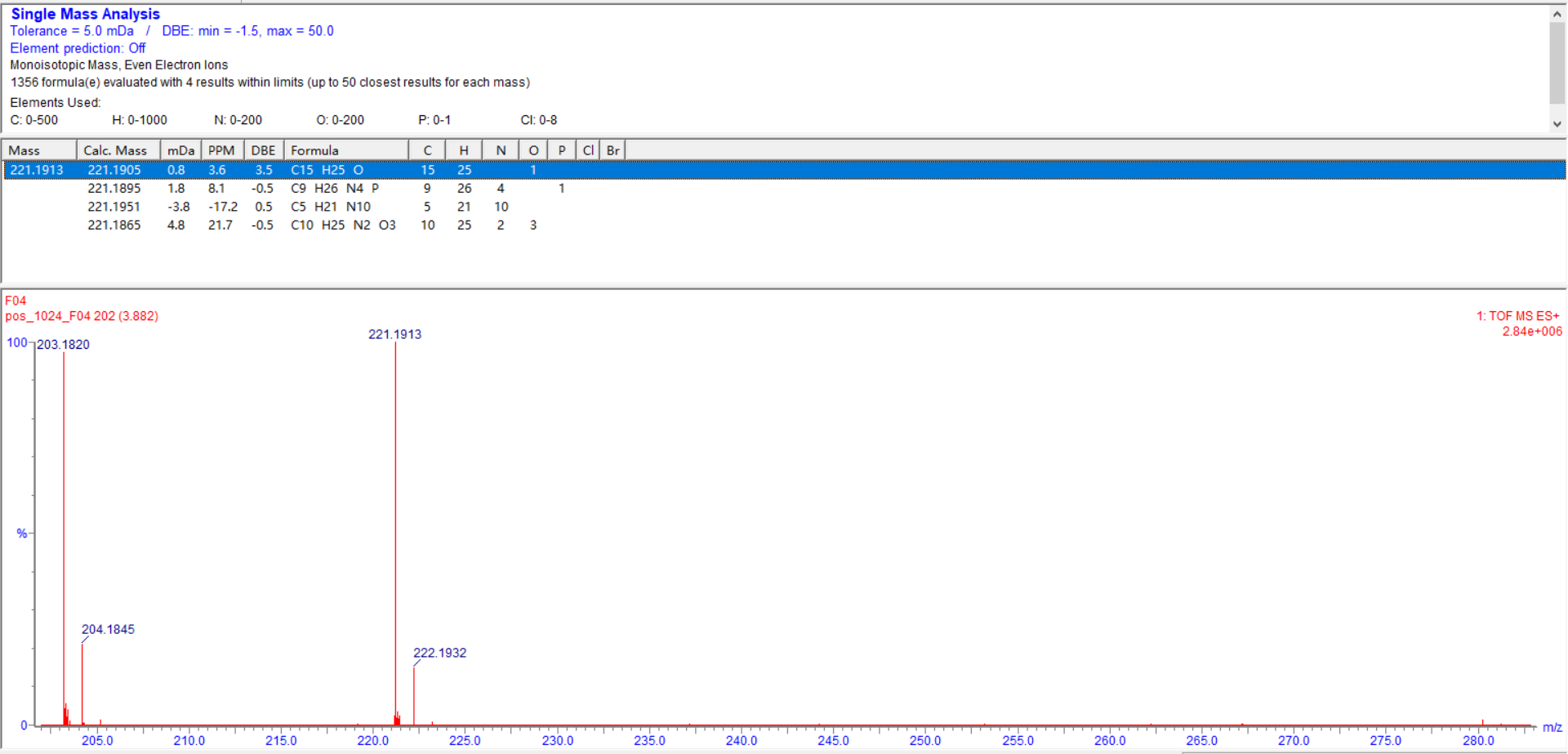

**Supplemental Figure S21. HRESIMS spectrum of compound 2**

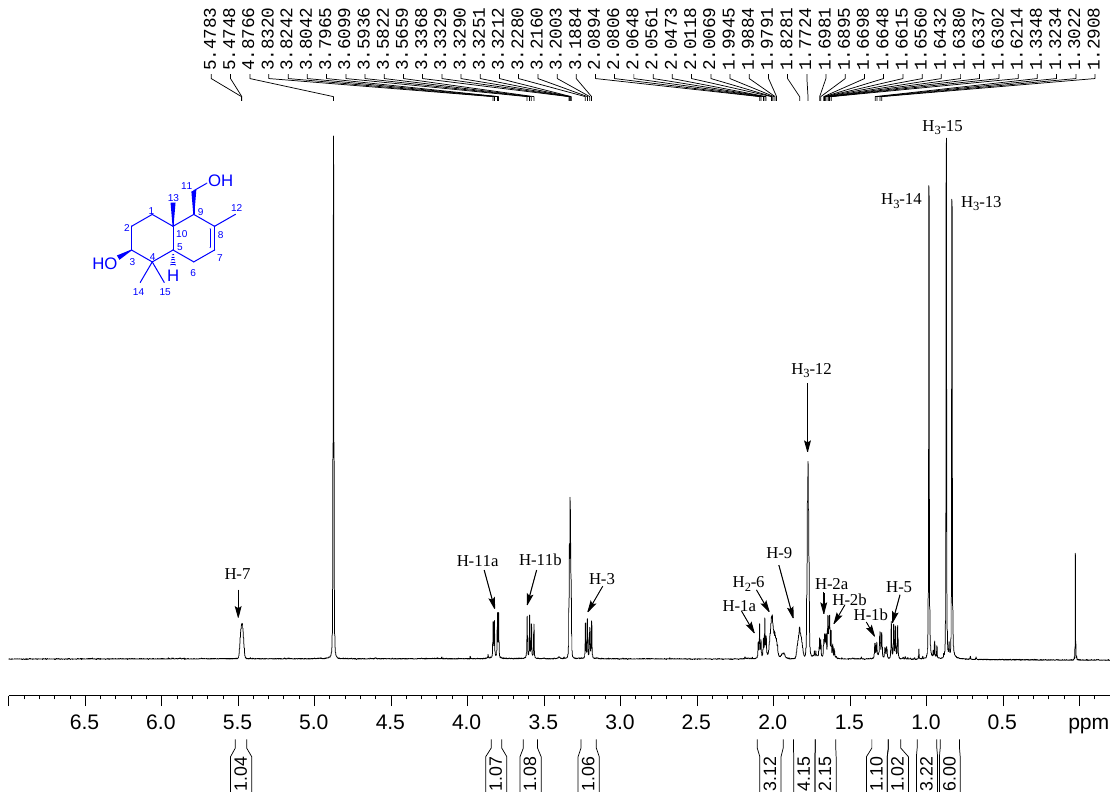

**Supplemental Figure S22. ^1^H NMR spectrum of compound 2 (400 MHz, CD_3_OD)**

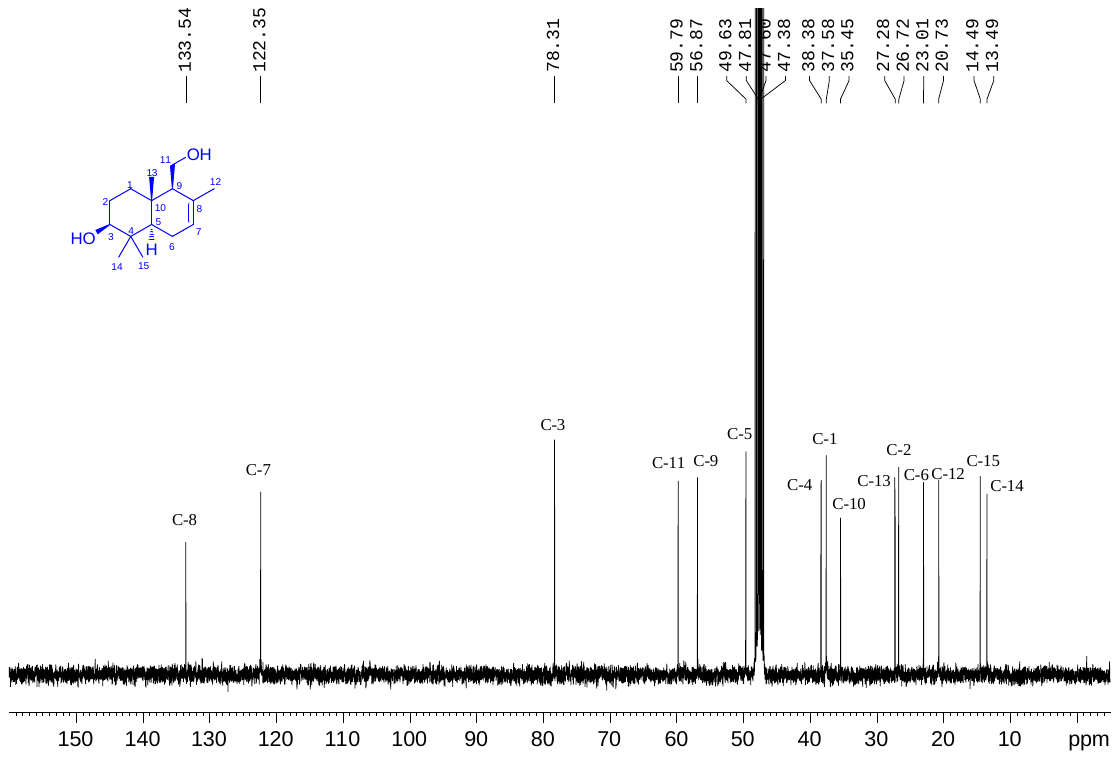

**Supplemental Figure S23. ^13^C NMR spectrum of compound 2 (100 MHz, CD_3_OD)**

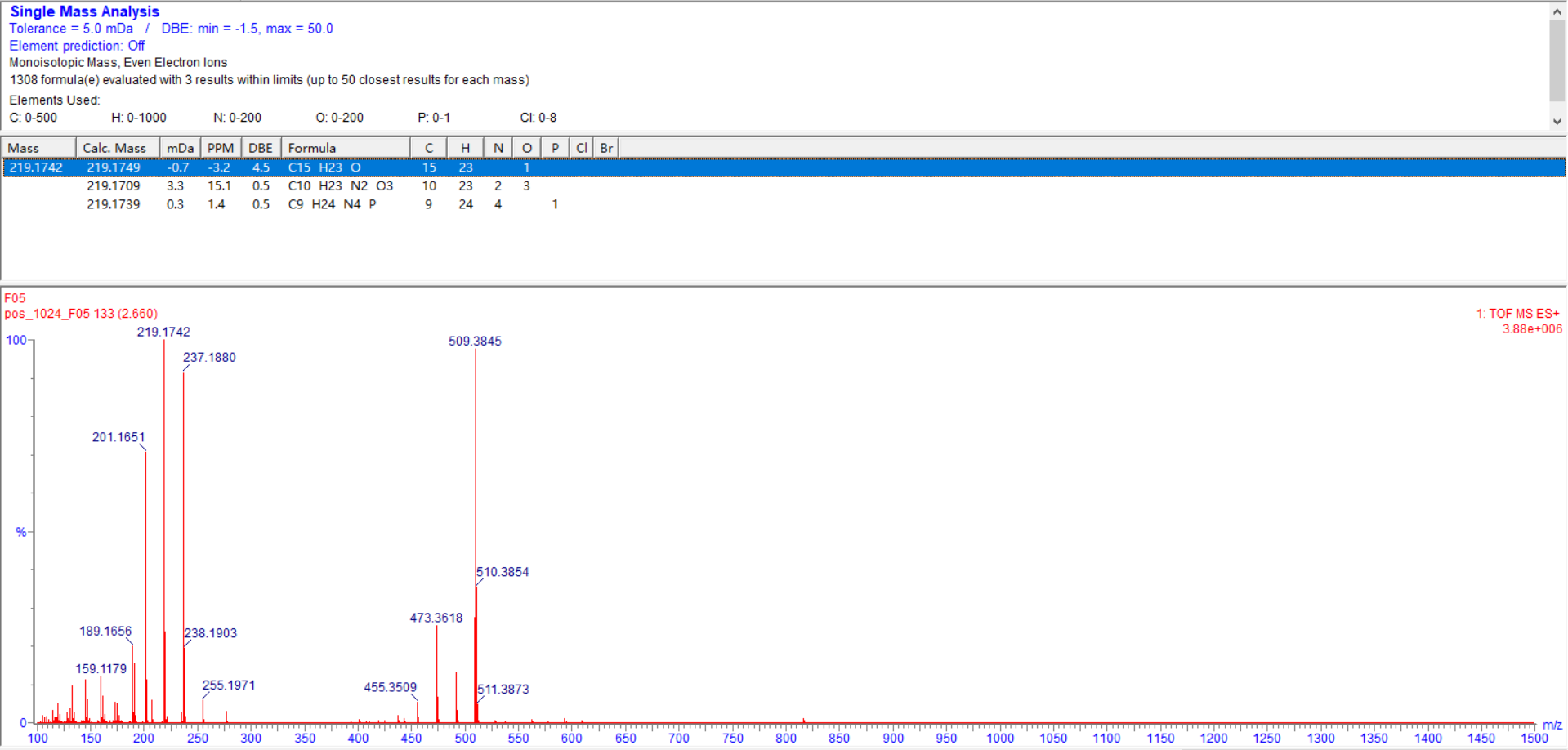

**Supplemental Figure S24. HRESIMS spectrum of compound 3**

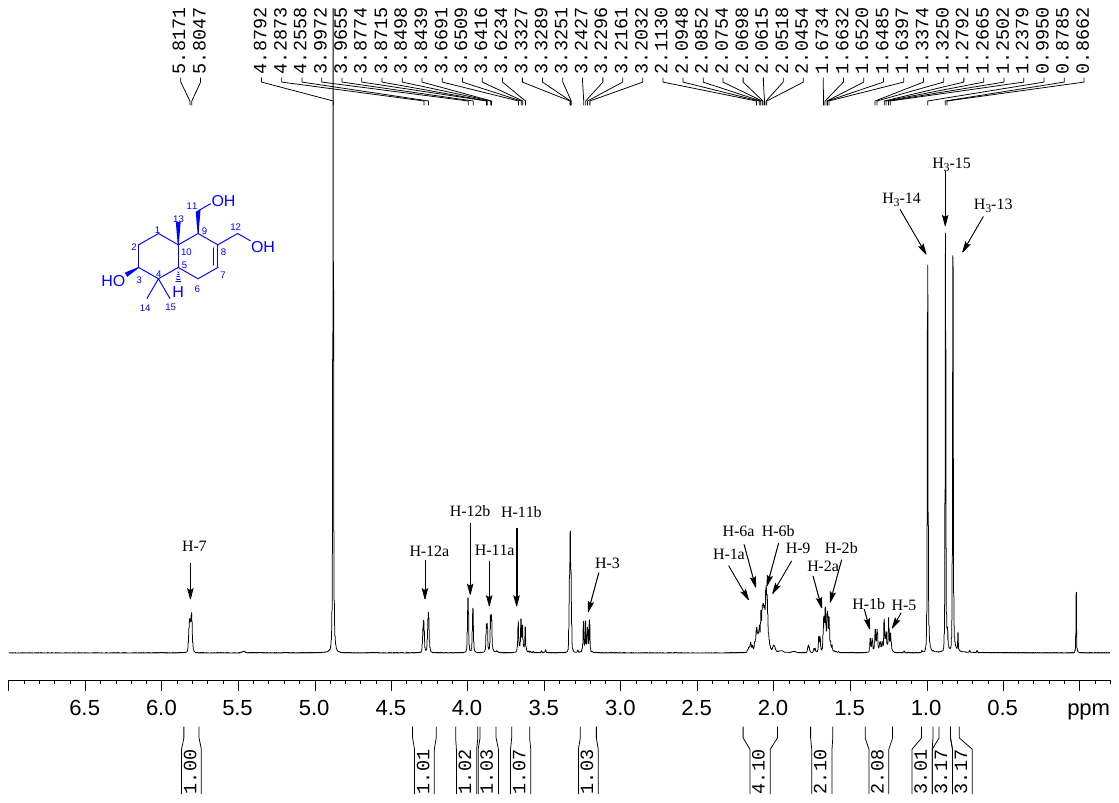

**Supplemental Figure S25. ^1^H NMR spectrum of compound 3 (400 MHz, CD_3_OD)**

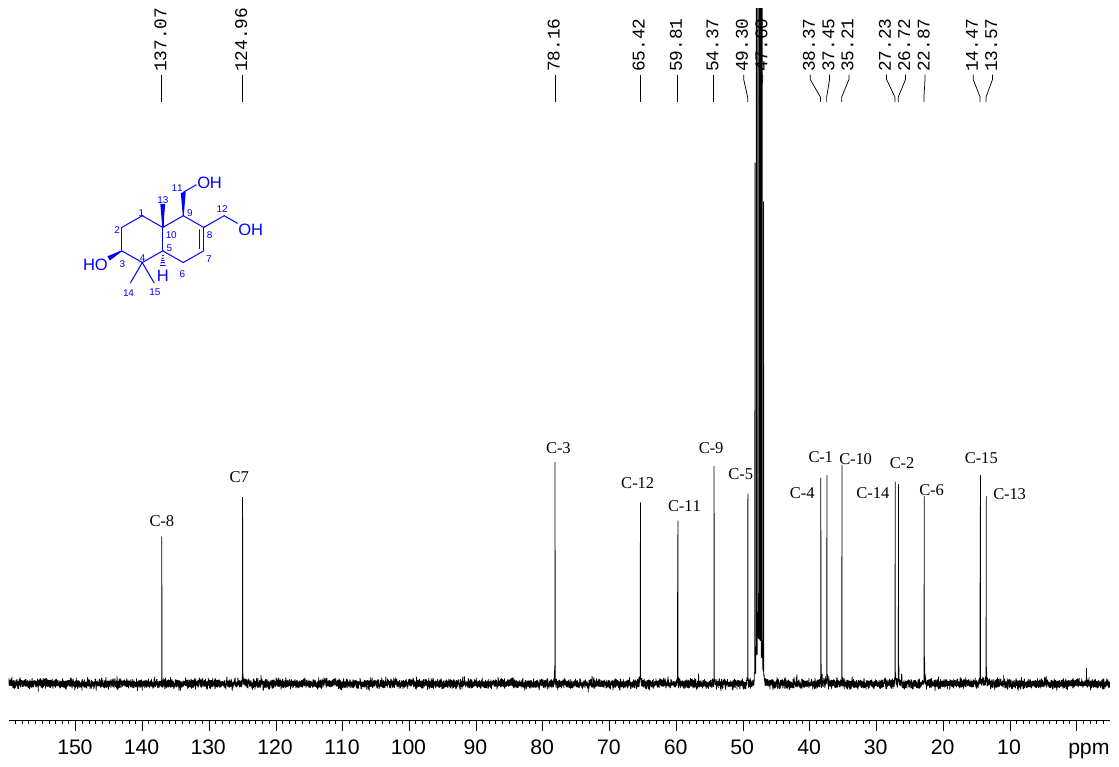

**Supplemental Figure S26. ^13^C NMR spectrum of compound 3 (100 MHz, CD_3_OD)**

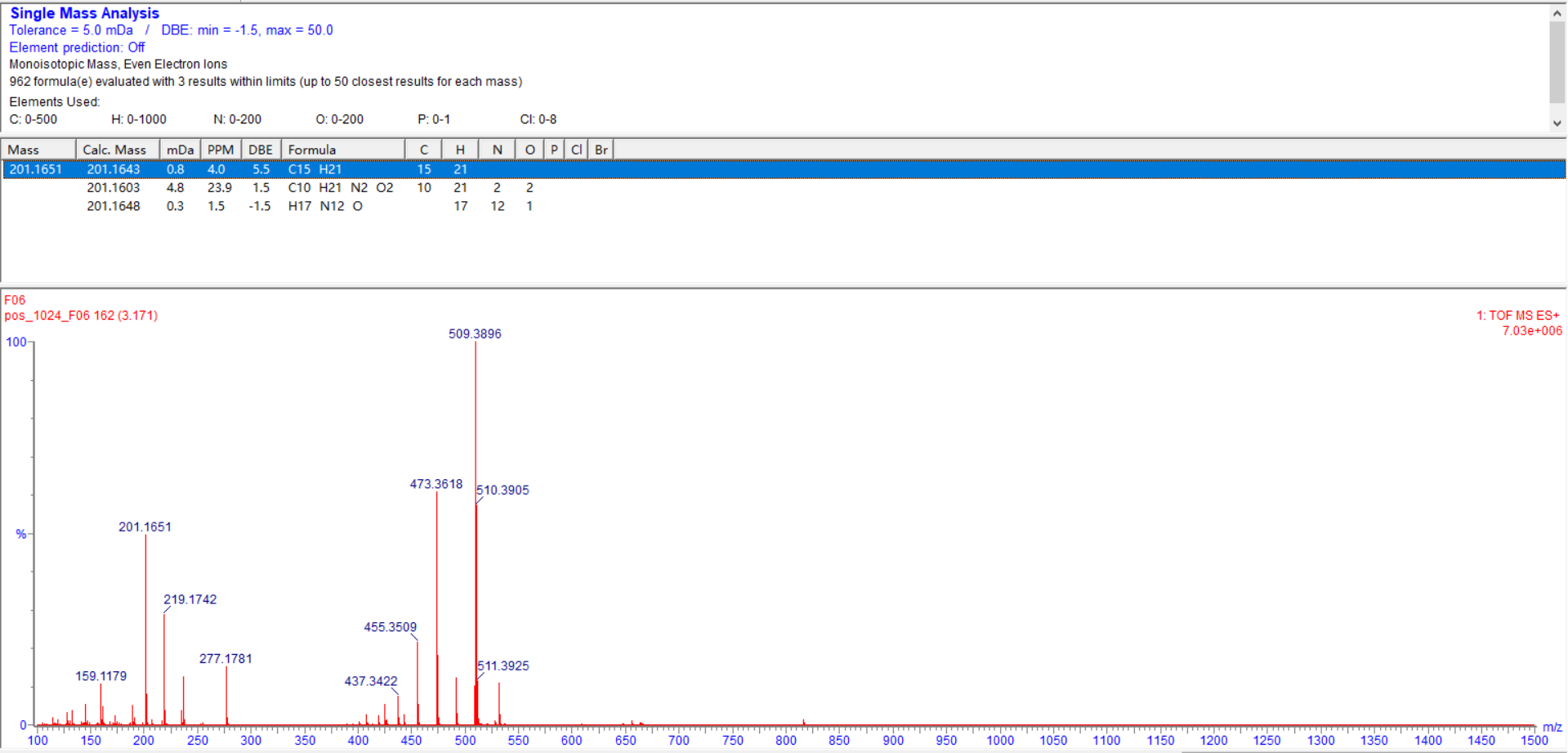

**Supplemental Figure S27. HRESIMS spectrum of compound 4**

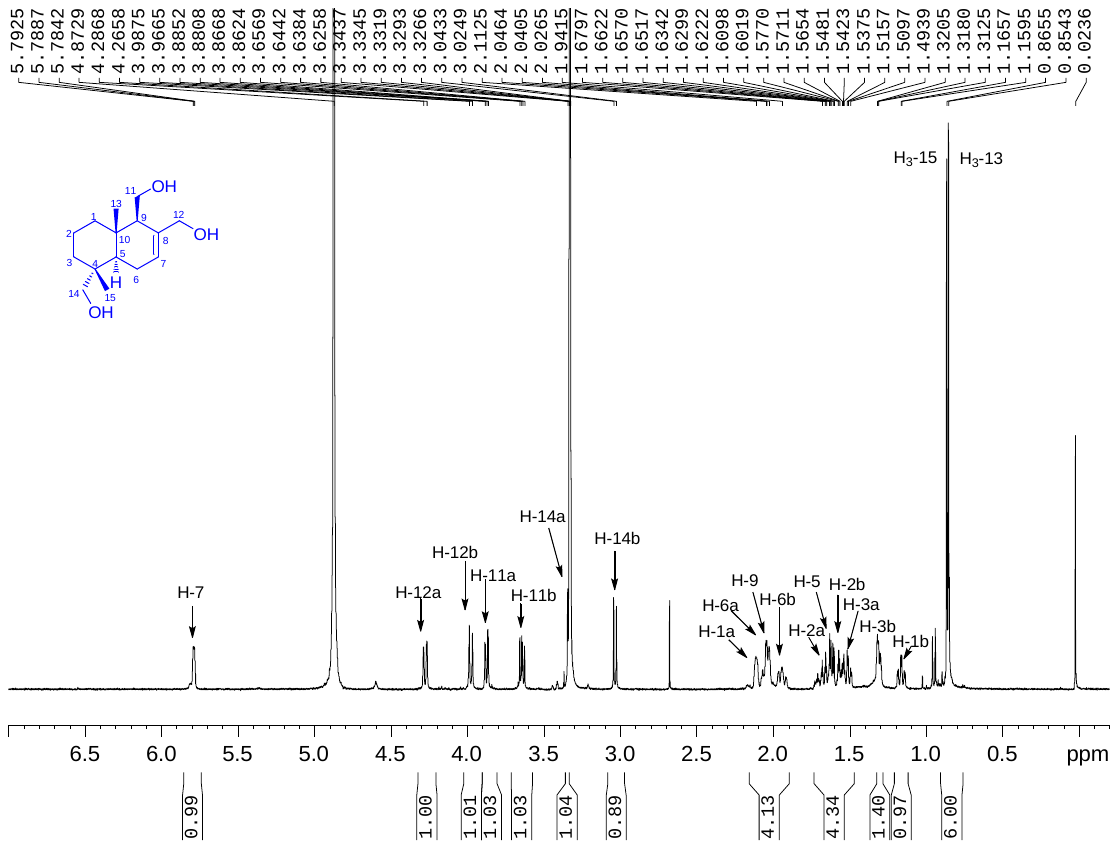

**Supplemental Figure S28. ^1^H NMR spectrum of compound 4 (400 MHz, CD_3_OD)**

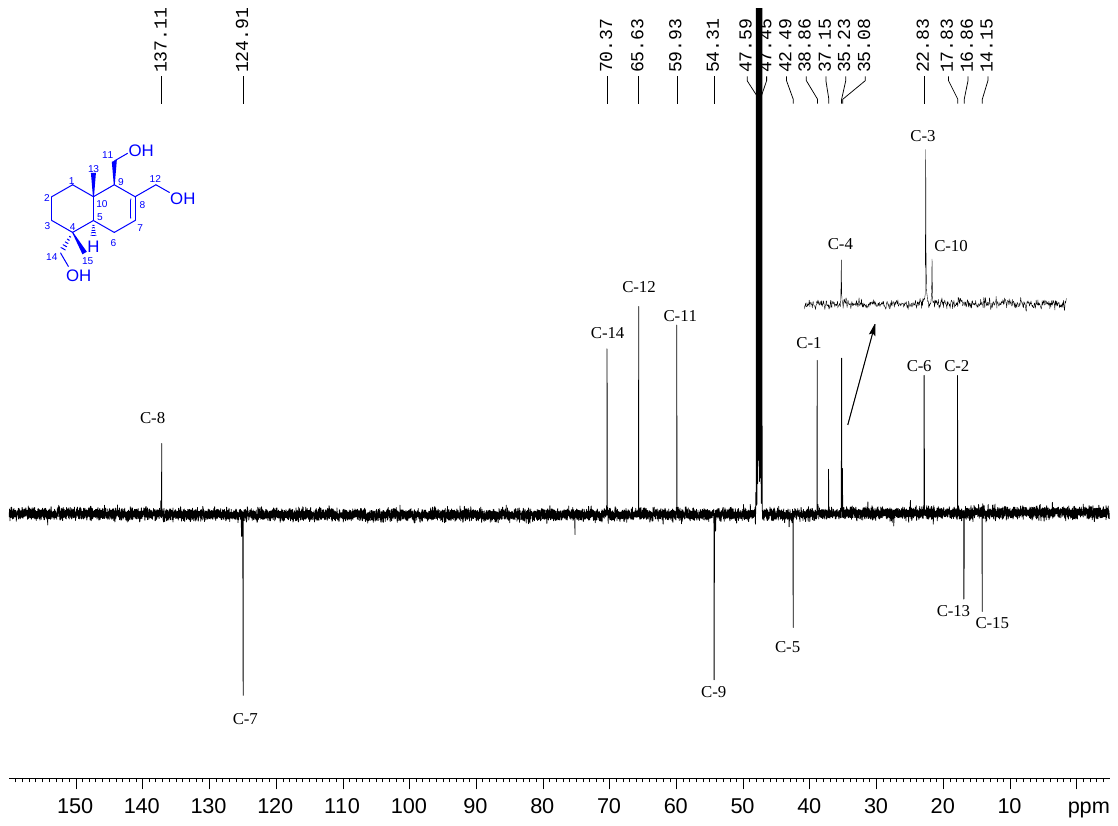

**Supplemental Figure S29. APT spectrum of compound 4 (100 MHz, CD_3_OD)**

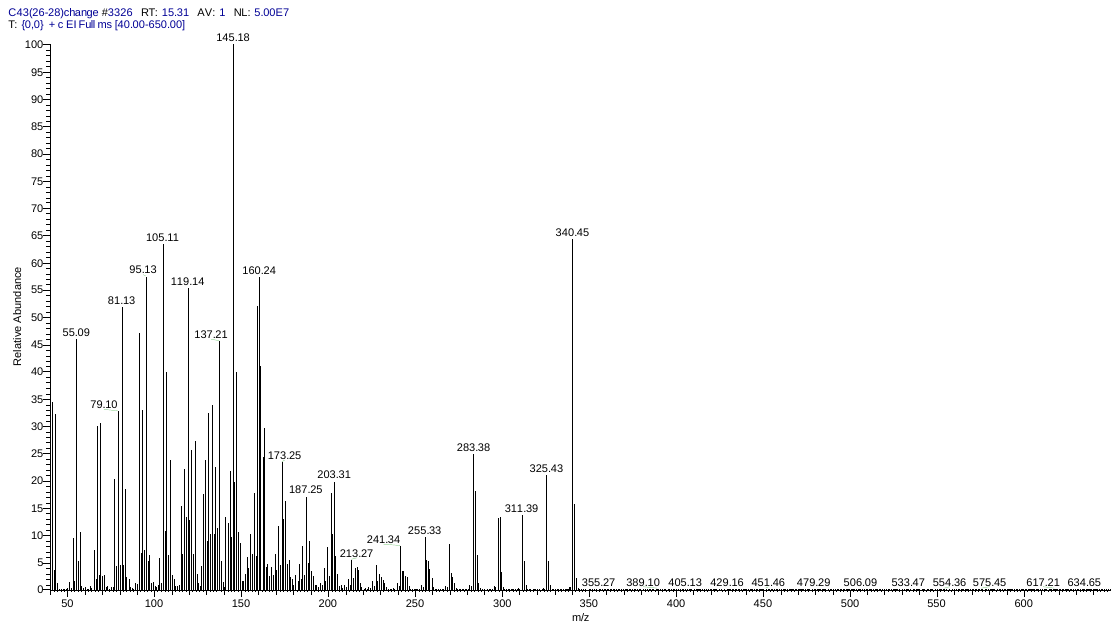

**Supplemental Figure S30. EIMS spectrum of compound 5**

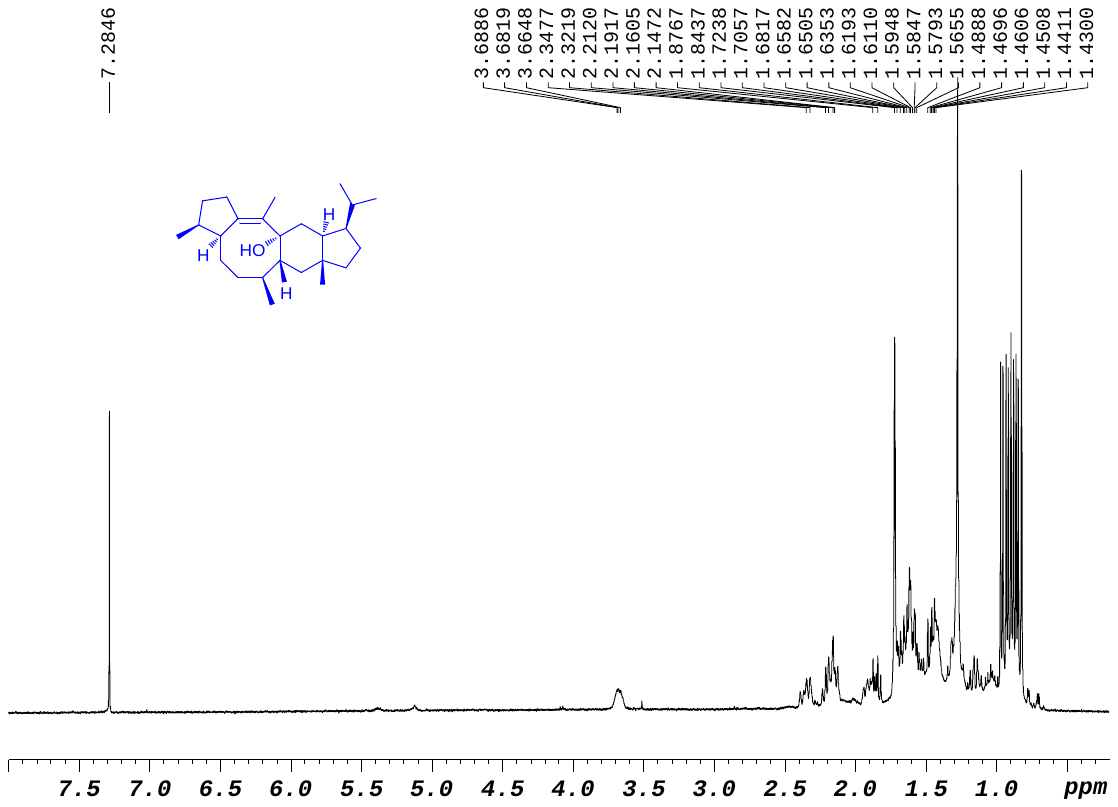

**Supplemental Figure S31. ^1^H NMR spectrum of compound 5 (400 MHz, CDCl_3_)**

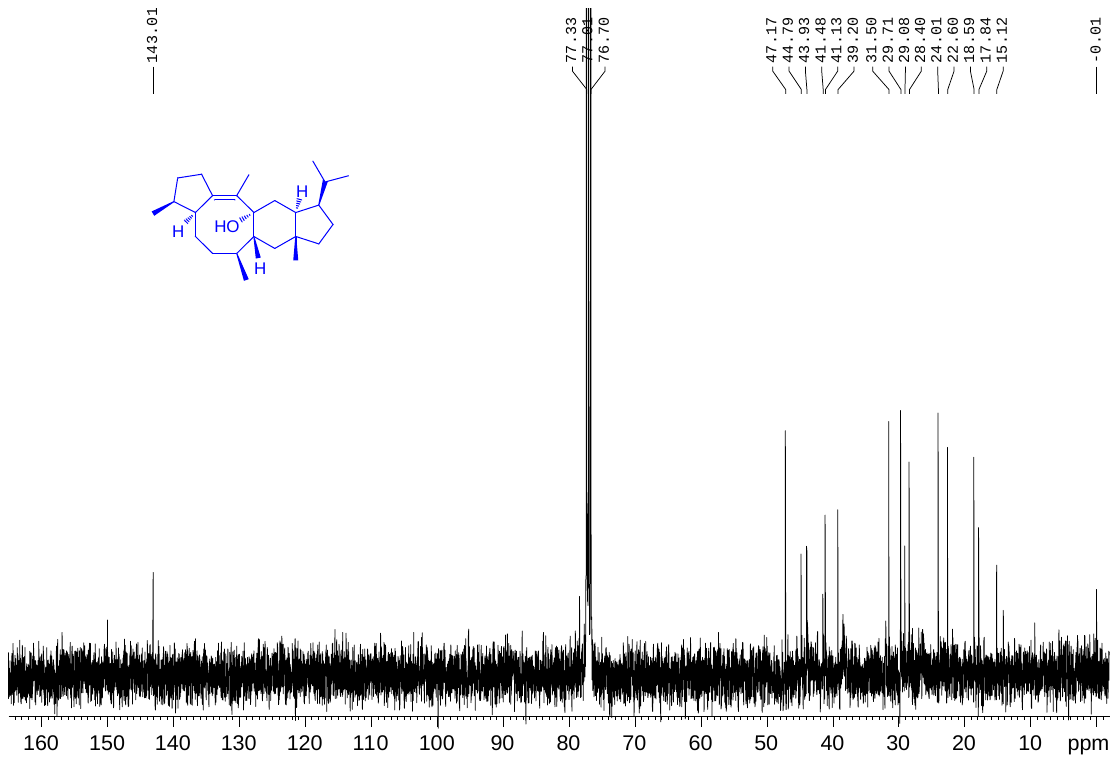

**Supplemental Figure S32. ^13^C NMR spectrum of compound 5 (100 MHz, CDCl_3_)**

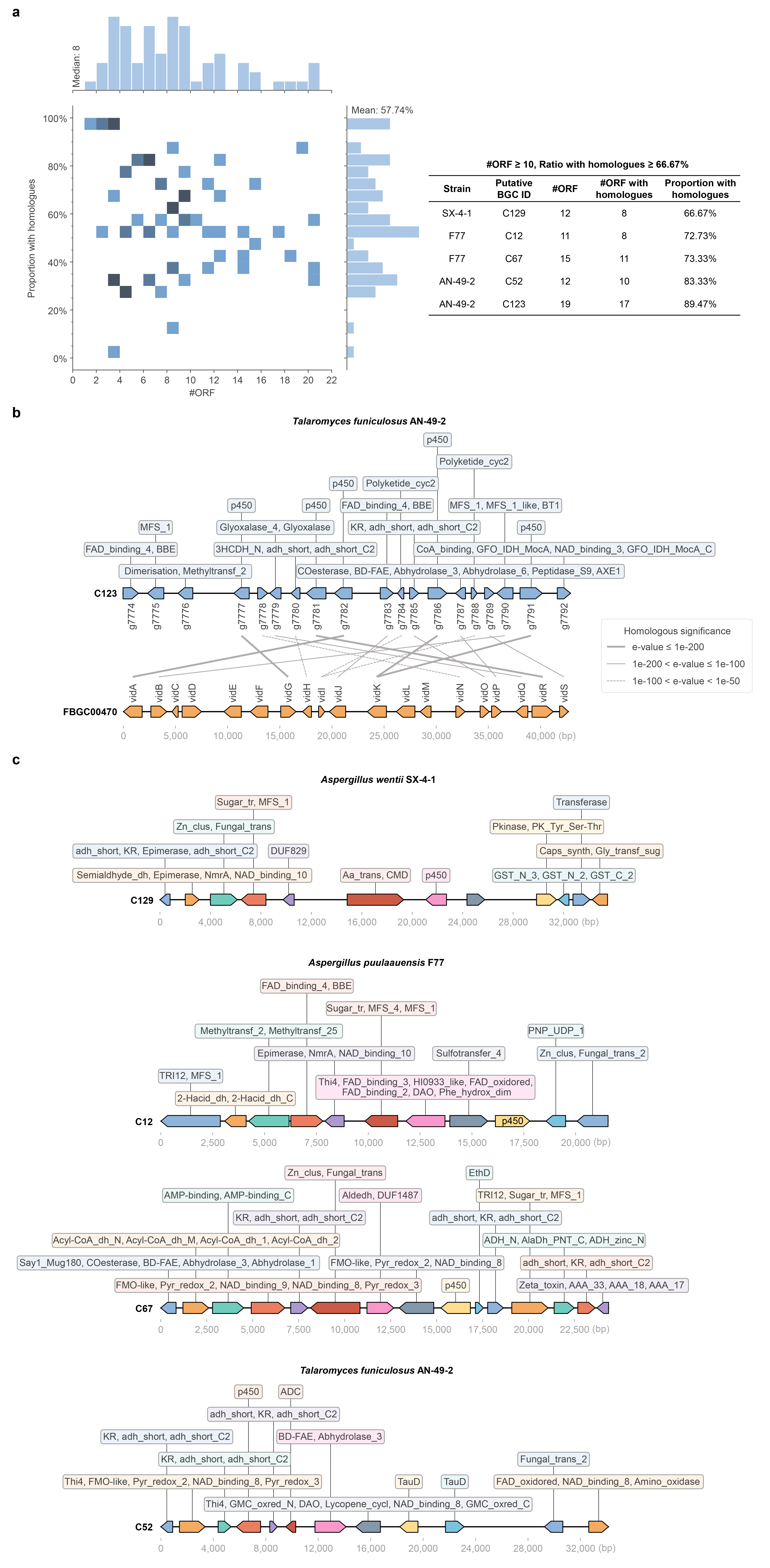

**Supplemental Figure S33. Non-canonical gene clusters exclusively predicted by f-BGM**

**a** Protein BLAST analysis between f-BGM-specific non-canonical gene clusters and FunBGCs-validated BGCs. The gene clusters are shown through the joint distributional histogram illustrating both ORF number (x-axis) and proportion of member ORFs with at least one homologue (e-value≤1×10^-30^) in FunBGCs (y-axis). Basic information of 5 representative gene clusters whose ORF number ≥10 and proportion with homologues ≥66.7% is listed in the right-sided table. **b** C129 sourced from the SX-4-1 genome exhibits high homology with a validated sub-BGC (FunBGCs ID: FBGC00470) involved in the biosynthesis of demethoxyviridin. The inter-ORF homologous linkages are distinguishably shown at three significance levels. **c** Other representative non-canonical gene clusters listed in (**a**) and their constitution of Pfam domains.
